# TNF inhibition reveals a reversible component of age-associated sinoatrial node dysfunction

**DOI:** 10.64898/2026.11.24.768532

**Authors:** Sacha Moore, Sami Al-Othman, Roman Tikhomirov, Sooraj Nair, Alice J Francis, Megan McKie, Matthew Smith, Yakun Wu, Eleonora Torre, Merve Keles, Xinning Xu, Steven A Niederer, Mark R Boyett, Luke B Roberts, Kazumasa Kanemaru, Pietro Mesirca, Matteo E Mangoni, Alicia D’Souza

## Abstract

**Background and Aims:** Age-related sinoatrial node (SAN) dysfunction is a principal indication for permanent pacing, yet its mechanisms remain poorly defined, and no disease modifying therapy exists. This study investigated whether age-related reorganisation of the local immune microenvironment suppresses the SAN pacemaker gene programme and investigated the functional and therapeutic relevance of implicated inflammatory signalling mediators.

**Methods:** Young and aged human SANs were profiled by spatial transcriptomics, integrated with physiological, molecular and immune phenotyping of aged mice. Candidate mediators were then tested through inducible expression restricted to HCN4⁺ pacemaker myocytes, pharmacological inhibition, and in living human SAN slices.

**Results:** The aged human SAN showed altered cellular composition, coordinated suppression of the pacemaker ion channel programme and inflammatory signalling concentrated in immune niches adjacent to pacemaker myocytes. Aged mice recapitulated these features, with SAN dysfunction, ion channel remodelling and immune expansion and activation. TNF emerged as a convergent candidate mediator. Subacute TNF exposure reduced HCN4 in living human SAN slices, and selective TNF induction in HCN4⁺ myocytes produced SAN dysfunction in structurally normal hearts, with slowed pacemaking and conduction attributable to reduced funny current (*I*_f_) and altered voltage dependence of the Ca_V_1.3 mediated component of the L-type Ca²⁺ current (*I*_Cav1.3_). Conversely, the TNF inhibitor etanercept reversed established electrical and molecular remodelling in aged mice, normalised the heart rate achieved during β-adrenergic stimulation, reduced frailty and improved survival during treatment.

**Conclusions:** Age-associated SAN dysfunction involves a therapeutically modifiable inflammatory component. Systemic TNF inhibition restores intrinsic SAN function in aged mice, identifying TNF signalling as a candidate therapeutic target.

**Translational Perspective:** Sinoatrial node (SAN) dysfunction is a major indication for permanent pacing, but the molecular basis of human SAN ageing and whether established disease can be modified remain poorly defined. We provide, to our knowledge, the first age-resolved spatial analysis of the human SAN at cellular resolution, complemented by studies in viable adult human SAN slices. TNF signalling emerged as a functionally important and therapeutically modifiable component of SAN dysfunction: TNF induction in HCN4⁺ pacemaker myocytes reproduced key features of SAN dysfunction, whereas TNF blockade with etanercept reversed established electrical and molecular abnormalities in aged mice, reduced frailty and improved survival. These findings show that age-associated SAN dysfunction has a reversible inflammatory component and identify TNF signalling within the nodal microenvironment as a candidate target for disease-modifying therapy.

**Summary:** Age-related sinoatrial node (SAN) dysfunction is a major cause of symptomatic bradyarrhythmia and permanent pacemaker implantation, but its molecular basis in the human SAN, and whether it is reversible, have not previously been investigated. Here we combine detailed studies of young and aged human SANs, with multiscale physiological, immunological, genetic and pharmacological investigation in mice to identify inflammatory remodelling as a functionally important feature of cardiac pacemaker ageing.

To our knowledge, this study provides the first age-resolved spatial characterisation of the human SAN at cellular resolution. Xenium profiling revealed altered cellular composition and local organisation in aged tissue, with increased representation and inflammatory activation of immune-cell populations alongside coordinated suppression of the pacemaker ion channel programme. Aged mice reproduced the principal electrical and molecular features of the human phenotype, including intrinsic bradycardia, prolonged SAN recovery time, slowed sinoatrial conduction and ion channel remodelling, and flow cytometry resolved selective expansion and activation of nodal T-cell and myeloid populations. TNF emerged as a conserved and experimentally tractable mediator. In a newly generated conditional line, TNF expression within HCN4⁺ pacemaker myocytes was sufficient to produce SAN disease in structurally normal hearts, including bradycardia, PR prolongation, sinus pauses, atrial ectopy, displacement of the leading pacemaker site, accompanied by remodelling of *I*_f_ and *I*_Cav1.3_ in through reduced current density and altered voltage dependence. Using a newly established viable human SAN slice preparation, we further showed that TNF suppresses HCN4 expression in native adult human nodal tissue. Importantly, eight weeks of systemic etanercept treatment restored key electrophysiological and molecular abnormalities in aged mice, reduced frailty and improved survival.

The ageing pacemaker therefore retains substantial biological plasticity, and TNF-dependent inflammatory signalling emerges as a candidate disease-modifying target in a condition currently managed only by device implantation.

## Introduction

Sinoatrial node dysfunction (SND) is a major cause of bradyarrhythmia in older adults and a leading indication for permanent pacemaker implantation.^1,2^ More than 1.2 million pacemakers are implanted worldwide each year. Although pacing relieves symptoms, it neither modifies the underlying disease nor restores native autonomic rate responsiveness, and it exposes patients to procedural and infection risks and the cumulative burdens of lifelong pacing, borne disproportionately by frail, multimorbid older adults; in Europe, device infection costs €22,000–34,000 and carries mortality of up to 11%.^3–5^

Despite this clinical burden, the mechanisms underlying human SND are poorly understood. Human studies have been dominated by histology and structural mapping, establishing the sinoatrial node (SAN) as a heterogeneous, fibrotically insulated pacemaker complex but leaving the impact of ageing itself to be inferred largely from low-resolution pathological analysis. Importantly, the largest histological series of human SANs did not support a simple monotonic increase in SAN collagen with age^6^ arguing against the commonly-accepted idea that human SAN ageing can be explained by progressive extracellular matrix (ECM) accumulation alone. What remains largely unknown is how ageing remodels the local SAN microenvironment, including the stromal and immune niches^7^ that surround and interact with pacemaker myocytes. Rodent studies have described intrinsic electrophysiological remodelling characterised by reduced expression or function of inward current carrying pacemaker ion channels, altered PKA-cAMP signalling and diminished βadrenergic sensitivity.^8–15^ These observations define important age-related defects in SAN automaticity, but leave open the questions of whether these apply to humans, what drives remodelling in the ageing SAN and whether this process is therapeutically tractable.

An important unresolved possibility is that, alongside structural and electrophysiological remodelling, the ageing SAN also develops a local inflammatory programme, especially as ‘inflammageing’^16–19^ and immune-inflammatory signalling are now recognised as important determinants of remodelling and arrhythmogenesis. Circulating cytokines such as IL-6, TNF and IL-1β rise with age and are associated with frailty, cognitive and cardiovascular decline and mortality.^20^ Extensive experimental evidence demonstrates that these same cytokines modulate cardiac electrical activity through direct and indirect effects on ionic currents, Ca^2+^handling processes, gap junctional coupling as well as ECM deposition.^21–27^ Furthermore, SND is common in systemic inflammatory states, including sepsis, autoimmune flares and COVID-19, and in cardiomyopathies with inflammatory aetiologies.^28,29^ Together, these observations implicate inflammation as a plausible upstream driver of age-associated SAN remodelling and dysfunction. However, this concept has not been directly investigated in the ageing SAN in humans or in complementary experimental models. We therefore combined spatial transcriptomics of young and aged human SAN with functional and molecular studies in living human SAN slices and complementary mouse models to test the hypothesis that ageing establishes a local inflammatory microenvironment with direct functional implications for SAN pacemaking and conduction.

## Methods

Full methodological detail, including ethical approvals, is provided in the **Supplemental Material**; all human and animal studies complied with applicable national and institutional regulations. SAN tissue from donor hearts unsuitable for transplantation was profiled by Xenium spatial transcriptomics.^30^ and viable human SAN slices were maintained ex vivo for intracellular action potential recording using sharp microelectrodes. Cardiac structure and global function were assessed by echocardiography. SAN function was characterised *in vivo* by continuous biotelemetry and anaesthetised surface electrocardiography, in Langendorff-perfused hearts by programmed electrical stimulation, and in isolated preparations and myocytes by optical and multielectrode-array mapping, sharp microelectrode recording³¹ and whole-cell patch clamp.^30–33^ Molecular, structural and immune remodelling was assessed by immunohistochemistry, immunoblotting, quantitative PCR and flow cytometry. *A priori* power calculations determined sample sizes, assuming 80% power and a two-sided α of 0.05, with spontaneous SAN beating rate as the primary endpoint. After assessment of data distribution, two-group comparisons used Welch’s t-test or the Mann-Whitney U test, matched comparisons used paired t-tests, three-group comparisons used one-way ANOVA with post hoc testing, categorical data used χ² or Fisher’s exact tests, and differential expression used DESeq2 with false discovery rate correction. All tests were two-tailed, significance was defined as P<0.05, and data are presented as mean ± SD unless otherwise stated.

## Results

### Spatial profiling of the aged human SAN reveals altered cellular organisation, suppression of the pacemaker electrical programme and immune cell remodelling

Human SAN tissue was obtained from acutely explanted donor hearts following non-cardiac causes of death. A discovery cohort of young (16–36 years, N=6) and aged (61–80 years, N=6) individuals was used for spatial profiling and histological analysis, with a further six young and six aged donors provided tissue for independent transcript validation. Available clinical histories are detailed in **Supplemental Table 1**. The SAN was localised in serial epicardial-to-endocardial sections using established anatomical landmarks^34,35^ and identified by low connexin 43 (Cx43) expression and enrichment of HCN4, the channel carrying the funny current (*I*_f_) (**Figure 1A**). A discrete HCN4⁺/Cx43⁻/Ca_V_1.3⁺ nodal region was reliably identified in three young and four aged hearts of the discovery cohort (**Supplemental Figure 1A**). These seven preparations underwent spatial transcriptomic profiling (10x Genomics Xenium In Situ) using a 480-gene panel (**Supplemental Table 3**) to resolve pacemaker cardiomyocytes and their local immune microenvironment within native tissue architecture. Key findings from cell-level analyses were assessed through donor-level pseudobulk analyses, protein and histological measurements, and transcript validation in an independent cohort in which the nodal region was identified using the same markers.

**Figure 1.**
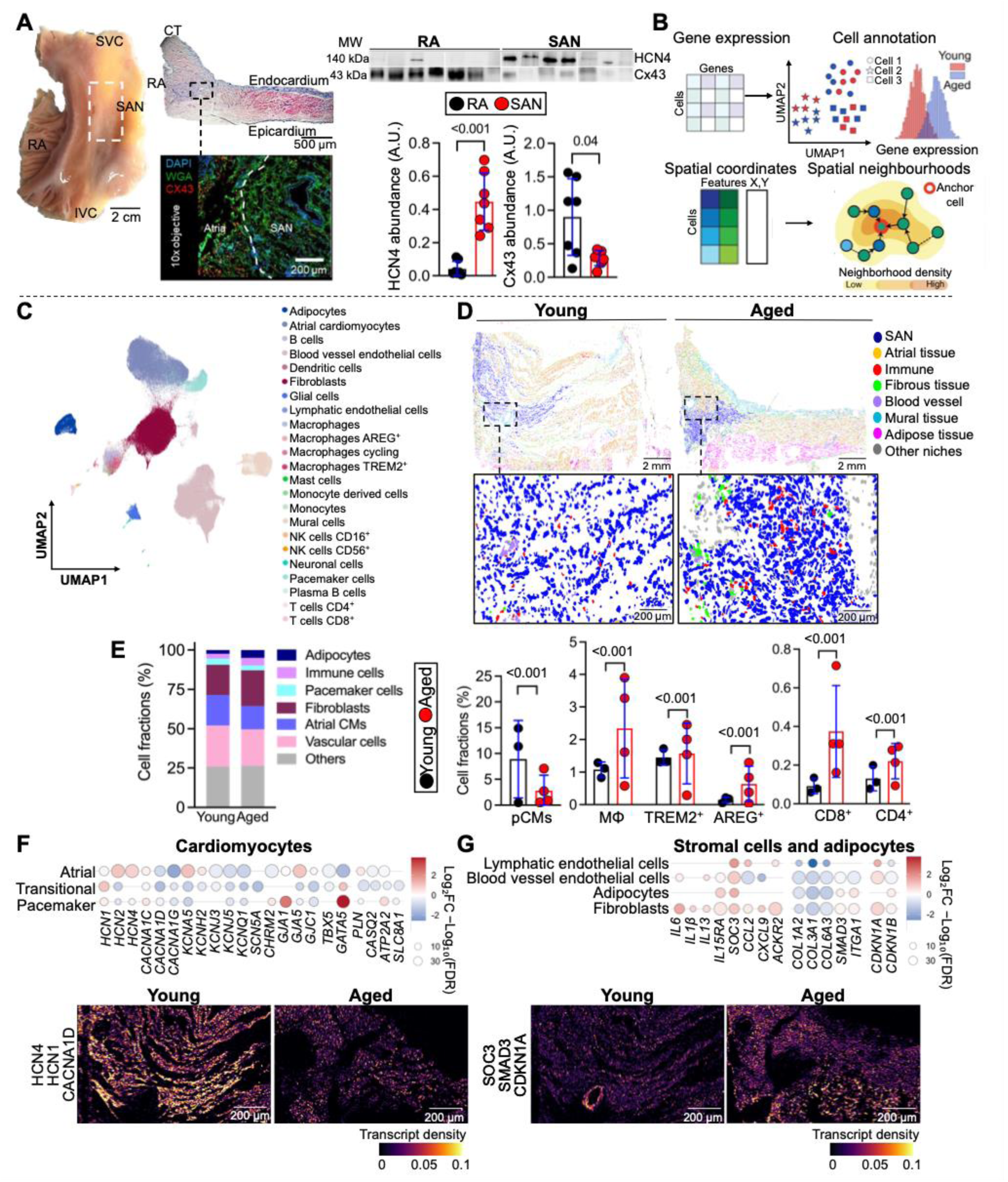
Spatially resolved profiling reveals age-associated cellular remodelling of the human SAN. **(A)** Left: representative anatomical dissection of the human SAN, Masson’s trichrome staining, and immunofluorescence images, acquired at 10x magnification, show nuclei (DAPI, blue), cardiomyocyte membranes (wheat germ agglutinin, WGA, green), connexin 43 (Cx43, red). Scale bars, 2 cm, 500 μm, 200 μm. Right: representative western blots and quantification of stain-free total protein-normalised HCN4 and Cx43 protein abundance in RA and SAN tissue (N=7). SVC, superior vena cava; RA, right atrium; IVC, inferior vena cava. **(B)** Xenium V1 spatial transcriptomic analysis workflow, showing gene expression-based cell annotation and spatial coordinate-based niche identification. **(C)** UMAP representation of annotated cell populations identified across young (N=3) and aged (N=4) human SAN tissue. **(D)** Representative spatial maps showing annotated cellular niches in young and aged human SAN tissue. Lower panels show enlarged SAN regions. Scale bars, 2 mm (upper panels) and 200 μm (lower panels). **(E)** Left: stacked bar plots showing the relative proportions of major cell populations in young and aged SAN tissue. Right: Relative abundance of pacemaker cardiomyocytes (pCMs), macrophages, TREM2⁺ macrophages, AREG⁺ macrophages, CD8⁺ T cells, and CD4⁺ T cells in young (N=3) and aged (N=4) donors. Cell-type composition was assessed using the underlying pooled cell counts, preserving the information contained in the number of cells contributing to each proportion. **(F)** Cell-level differential expression analysis of atrial, transitional and pacemaker cardiomyocytes. Dot colour denotes log₂FC (red, increased in aged; blue, decreased in aged) and dot size represents −log₁₀(FDR). Representative Xenium transcript-density maps for HCN4, HCN1, and CACNA1D are shown below. Scale bars, 200 μm. **(G)** Cell-level differential expression analysis of stromal cells and adipocytes, including fibroblasts, adipocytes, blood vessel endothelial cells, and lymphatic endothelial cells. Representative Xenium transcript-density maps of SOCS3, SMAD3, and CDKN1A are shown below. Warmer colours indicate greater local transcript abundance. Scale bars, 200 μm. P-values computed by paired t-test (A), two-sided Fisher’s exact test (E), or Mann-Whitney U test with Benjamini–Hochberg correction (F, G).

After multimodal morphology-based segmentation and reference-guided annotation with TACCO gene-count correction^36^ 272,715 cells were resolved, spanning pacemaker and atrial myocytes, fibroblasts, vascular cells, adipocytes, immune cells and neurons (**Figure 1C**, **Supplemental Figures 2A-D**). Spatial niches were then inferred by combining cell coordinates with intrinsic and neighbourhood gene expression across increasing aggregation depths, with cluster stability assessed over k = 5-20 (**Figure 1B**, **Supplemental Figure 3A-C**). This resolved discrete SAN, atrial myocardial, vascular, fibrous and immune-associated niches which separated by donor and by niche identity in reduced-dimensional space (**Figure 1D**, **Supplemental Figure 3E**). Within the Xenium-profiled regions, the aged SAN showed lower relative representation of pacemaker and other cardiomyocyte populations and enrichment of immune populations, including TREM2⁺ and AREG⁺ macrophages and CD4⁺ and CD8⁺ T cells annotated using canonical myeloid and lymphoid markers (Figure 1E, **Supplemental Figure 2E,F**). Donor-level immunofluorescence measurements corroborated these findings, demonstrating increased TNF^+^ cells, CD45⁺ leukocytes and CD8⁺ T cells, alongside directionally consistent increases in AREG⁺CD68⁺ macrophages and CD4^+^ T cells in aged SAN tissue (**Supplemental Figure 1D,E**). Spatial neighbourhood analysis also revealed altered cellular organisation, including greater fibroblast–macrophage association and reduced homotypic pacemaker cell neighbourhood enrichment in aged sections (**Supplemental Figure 3D**). Pacemaker cardiomyocyte density, nearest-neighbour distance and mean distance to the three and five nearest pacemaker cardiomyocytes did not differ between young and aged donors (**Supplemental Figure 3G**), indicating that these organisational differences do not reflect a change in pacemaker cell density or local clustering.

Cell-resolved Xenium analysis identified age-associated remodelling of transcripts encoding ion channels, gap junction proteins and Ca²⁺-handling machinery (**Figure 1F**). Pacemaker cardiomyocytes showed lower expression of HCN4, CACNA1D (Ca_V_1.3), CACNA1G (Ca_V_3.1) and SLC8A1 (NCX1), consistent with suppression of components contributing to diastolic depolarisation and automaticity. Donor-level pseudobulk analysis showed concordant age-associated expression changes across atrial (R=0.69), transitional (R=0.83) and pacemaker cardiomyocytes (R=0.74), and across all immune and stromal populations examined (R=0.78–0.89; **Supplemental Figure 4A,B**). Independent validation demonstrated reduced HCN4 protein abundance and Cav1.3 immunoreactivity, together with reduced *HCN4*, *HCN1* and *TBX3* transcript abundance and a non-significant reduction in *CACNA1D* (**Supplemental Figures 1B,C and 4C**). Atrial cardiomyocytes showed a more mixed pattern, with increased HCN4, HCN2, KCNA5 (K_v_1.5, *I*_K,ur_) and GJA5/Cx40 but reduced KCNJ3 and KCNJ5, encoding the GIRK1 and GIRK4 subunits of the acetylcholine-activated potassium channel (*I*_K,ACh_), together with its upstream muscarinic receptor CHRM2. At cell resolution, transitional cardiomyocytes showed the broadest de-specialisation, with reduced SCN5A (Na_v_1.5, *I*_Na_), CACNA1D, KCNQ1 and KCNH2 (*I*_Ks_ and *I*_Kr_), KCNA5, GJC1, SLC8A1, ATP2A2 (SERCA2a), PLN and CASQ2, indicating combined impairment of excitability, sarcoplasmic reticulum Ca²⁺ handling and intercellular coupling. Cell-level analysis of fibroblasts showed increased expression of the senescence-associated markers CDKN1A and CDKN1B, together with elevated SMAD3, and the proinflammatory cytokines IL6 and IL1b. Collagen deposition was increased in aged relative to young SAN tissue (32.88±4.85% vs 14.92±2.11%; P=0.03), despite downregulation of the corresponding fibrillar collagen transcript COL3A1 and unchanged COL1A1 (**Figure 1G**, **Supplemental Figure 1F**).

Cell-resolved differential expression and pathway-enrichment analyses identified inflammatory transcriptional programmes across myeloid and lymphoid populations (**Figure 2A,B**). The most prominent signature was in AREG^+^ macrophages, with increased AREG, IL1B, CXCL3 and CDKN1A with enrichment for TNF signalling via NF-κB (FDR = 3.8 × 10⁻⁴) and IL6-JAK-STAT3 signalling (FDR = 0.038). TREM2⁺ macrophages showed a more restricted antigen-presentation programme, CD4⁺ T cells upregulated NFKBIA and CCL5 with TNF-NF-κB enrichment, and CD8⁺ T cells showed a cytotoxic programme (KLRK1, GZMK, EOMES). We next mapped the spatial distribution of this inflammatory signal across the aged SAN. Transcript-density mapping showed focal enrichment of TNF signalling and AREG rather than a uniform rise (**Figure 2C),** verified at the protein level (**Supplemental Figure 1D,E**). Spatial neighbourhood analysis identified AREG⁺ macrophages as the immune population showing the most pronounced age-associated change in spatial organisation relative to pacemaker cardiomyocytes, shifting from relative depletion in young SAN sections to enrichment in aged tissue (**Figure 2D**). To investigate the associated transcriptional changes, we compared pacemaker cardiomyocytes proximal and distal to AREG⁺ macrophages (**Figure 2E**). Cell-level analysis identified a transcriptional signature strongest at 60 μm, with 31 genes shared across the 30, 60 and 90 μm thresholds (**Supplemental Figure 3F**). Protein–protein interaction analysis revealed reduced expression of pacemaker and nodal identity genes (*HCN4*, *HCN1*, *TBX3*, *ATP2A2*, *CACNA1G*, *CASQ2*), alongside increased inflammatory and matrix-associated signalling (**Figure 2F**). Importantly, donor-level pseudobulk analysis confirmed significantly lower expression of *HCN4* (log₂FC=−0.63; FDR=0.009), *HCN1* (log₂FC=−0.68; FDR=0.022) and *CACNA1D* (log₂FC=−0.93; FDR=0.045) in macrophage-proximal pacemaker cardiomyocytes (**Supplemental Figure 4D,E**). This combination of inflammatory activation, loss of electrical identity and gain of a matrix-associated programme resembles partial dedifferentiation.^37,38^

**Figure 2.**
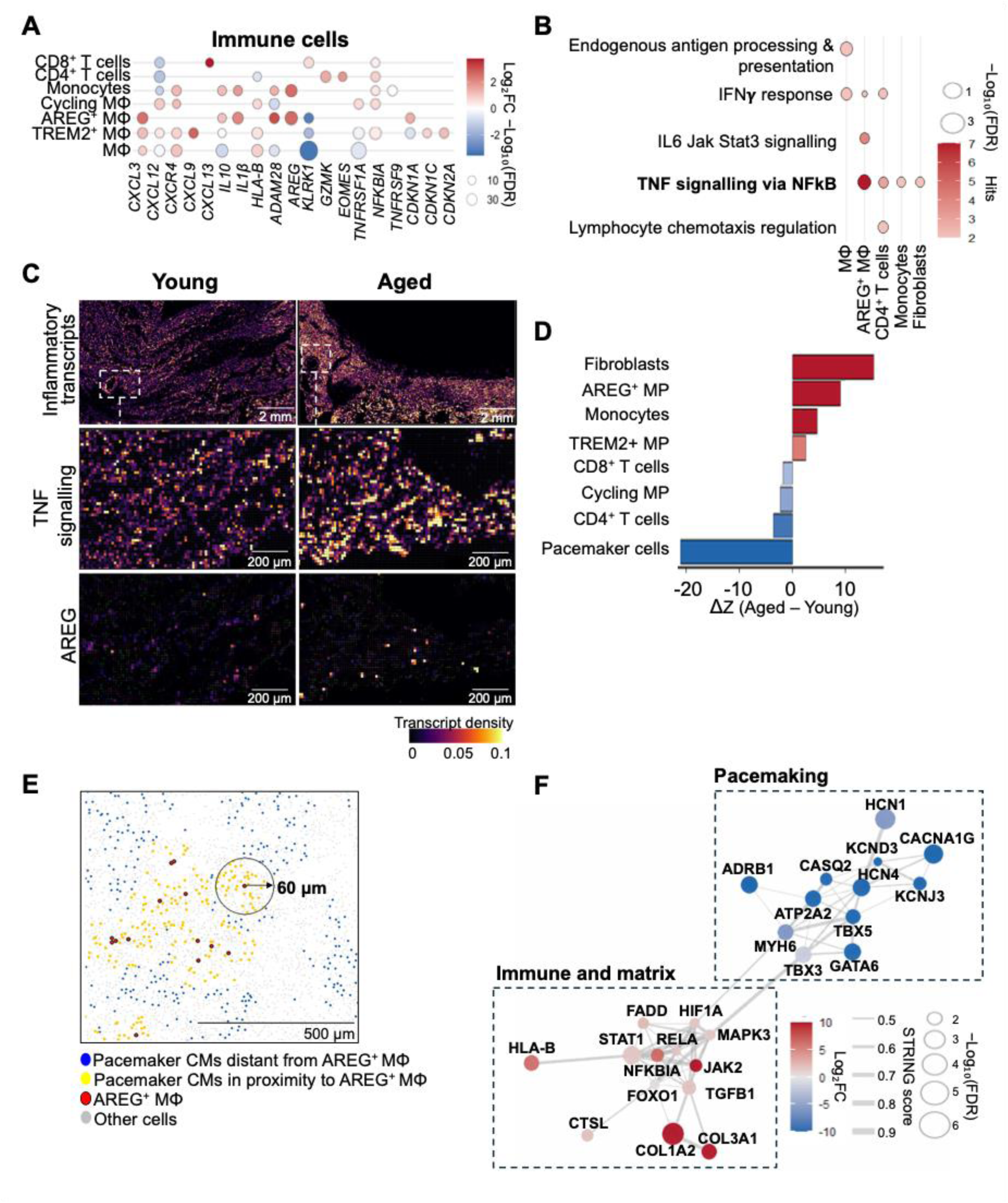
Spatial inflammatory niches and niche-associated pacemaker cell remodelling in the ageing human SAN. (**A)** Cell-level differential expression analysis of immune cell populations, including macrophages, TREM2⁺ macrophages, AREG⁺ macrophages, cycling macrophages, monocytes, CD4⁺ T cells and CD8⁺ T cells. Dot colour denotes log₂FC (red, increased in aged; blue, decreased in aged) and dot size represents −log₁₀(FDR). Only genes with FDR < 0.05 are shown. **(B)** Gene set enrichment analysis of cell-level differential expression findings using Gene Ontology and Hallmark pathway databases. Dot colour denotes the number of genes (hits) associated with each gene set and dot size represents −log₁₀(FDR). **(C)** Representative Xenium transcript-density maps showing inflammatory transcript density, TNF signalling, and AREG in young (N=3) and aged (N=4) human SAN tissue. Warmer colours indicate greater local transcript abundance. Scale bars, 2 mm (upper panels) and 200 μm (lower panels). **(D)** Differences in spatial neighbourhood-enrichment Z-scores (ΔZ) between aged and young SAN sections. Positive values indicate greater spatial association in aged tissue and negative values indicate reduced association. These permutation-derived scores characterise spatial organisation within the sampled sections rather than donor-level statistical significance. Cell-type pairs with |ΔZ| > 2.5 were considered to show prominent changes in neighbourhood enrichment. **(E)** Spatial proximity analysis of pacemaker cardiomyocytes and AREG⁺ macrophages. Pacemaker cardiomyocytes located within 60 μm of an AREG⁺ macrophage are shown in yellow, distal pacemaker cardiomyocytes in blue, AREG⁺ macrophages in red and remaining cells in grey. Scale bar, 500 μm. **(F)** STRING protein–protein interaction network of genes identified by cell-level differential-expression analysis of pacemaker cardiomyocytes proximal and distal to AREG⁺ macrophages. Node colour denotes log₂ fold change and node size represents −log₁₀(FDR) from the cell-level analysis. Donor-level pseudobulk analysis of the principal proximity-associated ion-channel genes is presented in **Supplemental** Figure 4D**, E.** Functional subnetworks associated with pacemaking, immune signalling, and extracellular matrix are highlighted. Cell-level differential expression was assessed using Mann– Whitney U tests with Benjamini–Hochberg correction (A, F). Pathway enrichment (B) and spatial neighbourhood enrichment (D) were assessed using permutation-based methods, with Benjamini–Hochberg correction where applicable. Neighbourhood enrichment Z-scores describe spatial associations within the sampled sections rather than donor-level significance.

### Aged mice recapitulate SND and inflammatory remodelling with expansion of immune cell populations

To determine whether inflammation accompanies age-related SAN dysfunction and underpins it mechanistically, we studied the naturally aged mouse, arguably the best characterised model of mammalian ageing,^39^ in which the electrophysiology, molecular profile and immune compartment of the node can be resolved together and interrogated genetically and pharmacologically. Aged mice (22 months, equivalent to 60–70 human years^40^) showed SND consistent with previous reports.^13,41^ Heart rate *in vivo* was lower than in young mice (by 52.11 BPM, 95% CI 5.17-99.06; P=0.03). Spontaneous beating rate of the isolated and denervated SAN was reduced by 24.3% and corrected SAN recovery time was prolonged (by 36.46 ms, 95% CI 7.90-65.03; P=0.02; **Figure 3A**). Optical mapping of isolated preparations showed slower conduction from the SAN to both atria (**Figure 3B**). *I*_f_ density was reduced in isolated pacemaker cells (**Figure 3C**; age × voltage interaction, P<0.001). At physiologically relevant diastolic potentials (−55 to −75 mV), mean inward current density was lower in aged than young myocytes, although these differences were not significant after multiple-comparison correction (**Supplemental Table 9**).

**Figure 3.**
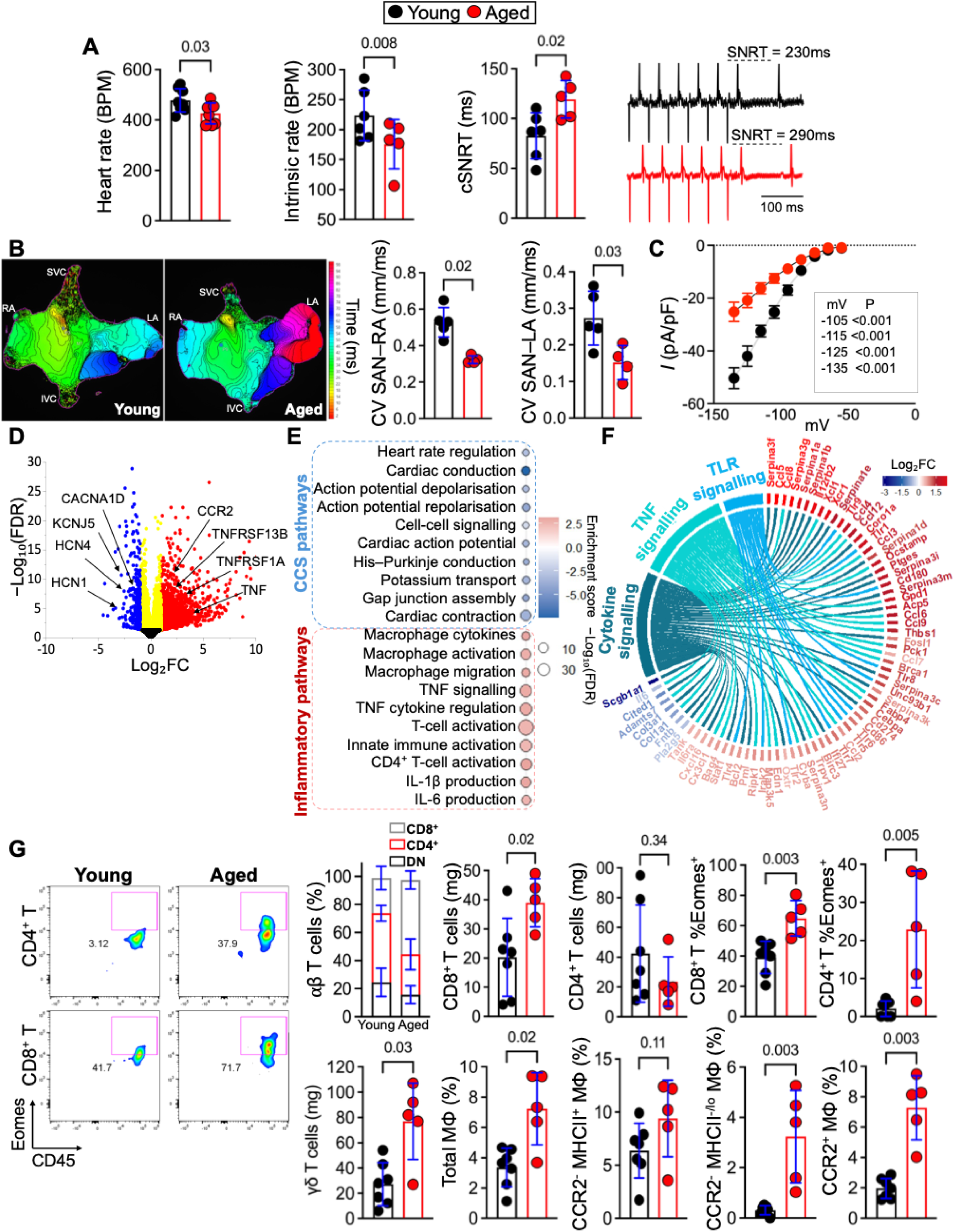
Aged mice recapitulate SND and inflammatory remodelling observed in the ageing human SAN. **(A)** Quantification of heart rate in vivo, spontaneous beating rate of the isolated SAN, and corrected sinoatrial node recovery time (cSNRT) in young (2–3 months, N=5-7) and aged (22 months, N*=*5-7) mice. Representative SNRT recordings are shown on the right. **(B)** Representative conduction maps from young (N=5) and aged (N=4) SAN. Quantification of SAN-to-RA and LA conduction velocity (CV). SVC, superior vena cava; RA, right atrium; LA, left atrium; IVC, inferior vena cava. **(C)** Current-voltage relationships of the hyperpolarization-activated funny current (*I*_f_) measured by whole cell patch clamp in isolated SAN myocytes from young (N=6 mice, n=16 cells) and aged (N=5 mice, n=16 cells) mice. Current amplitude was normalised to membrane capacitance (pA/pF). Current-voltage relationships were analysed using a mixed-effects model (REML) with age, test voltage and their interaction, followed by Šídák-adjusted multiple comparisons between groups at individual voltages. Mixed-effects analysis: age, P<0.001; voltage, P<0.001; age×voltage interaction, P<0.001. Voltage-specific statistics are provided in **Supplemental Table 9. (D)** Volcano plot showing differentially expressed genes in aged (N=8) relative to young (N=8) SAN tissue. Upregulated genes are shown in red, downregulated genes in blue, genes exceeding the log₂FC threshold alone in yellow, and non-significant genes in black. Representative inflammatory-and pacemaker-associated genes highlighted. **(E)** Gene ontology enrichment analysis of differentially expressed genes. Dot colour denotes enrichment score and dot size represents −log₁₀(FDR). Cardiac conduction-and inflammatory-related pathways enriched with ageing highlighted. **(F)** Chord diagram showing relationships between differentially expressed genes and enriched inflammatory pathways, including TNF, cytokine and toll-like receptor (TLR) signalling. Connecting ribbons denote genes contributing to each pathway. Gene labels are coloured according to log₂FC (red, upregulated; blue, downregulated). **(G)** Flow cytometric analysis of immune cell populations in young (N=7) and aged (N=5) SAN. Representative pseudocolour plots showing Eomes and CD45 expression within gated CD4⁺ and CD8^+^ T cells; values denote the percentage of cells within the parent population. Quantification of αβ T-cell subsets (CD8⁺, CD4⁺, and double-negative (DN)), CD8⁺, and CD4⁺ T cells, Eomes⁺ CD8⁺ and CD4⁺ T cells, γδ T cells, total macrophages (MΦ), CCR2⁻MHCII⁺ macrophages, CCR2⁻MHCIIlo macrophages, and CCR2⁺ macrophages. Data are presented as mean ± SD with individual biological replicates. P-values computed by Welch’s t-test (A), Mann–Whitney U test (B, G), mixed-effect analysis (C), or DESeq2 with Benjamini–Hochberg correction (D), permutation testing with Benjamini–Hochberg correction (E), or DESeq2 FDR < 0.05 (F).

This phenotype provided a defined functional context in which to examine proinflammatory signalling and the immune compartment. RNA sequencing of aged nodal tissue showed coordinated suppression of the pacemaker electrical programme, with downregulation of *Hcn4*, *Hcn1*, *Cacna1d* and *Kcnj5* and enrichment of downregulated pathways related to cardiac conduction, ion channel activity and regulation of heart rate (**Figure 3D,E**). The same tissue showed concurrent proinflammatory remodelling, with enrichment of macrophage and T-cell activation and migration pathways and upregulation of cytokines, chemokines and TNF superfamily members (**Figure 3F**). Annotation against curated ligand-receptor pairs^42^ identified twelve cytokine and TNF-family pairs upregulated with age, both partners increased and none reduced, of which seven were TNF superfamily pairs with *Tnf* and *Lta* both signalling through *Tnfrsf1a* (**Supplemental Figure 5A**). TNF upregulation was validated at the protein level by immunofluorescence, with a 6.5-fold increase in TNF^+^ cells in the aged SAN (**Supplemental Figure 5B**). Flow cytometry was used to define the immune compartment (**Supplemental Figure 5C**). The most prominent changes were expansion of γδ T cells (2.8-fold; P=0.03) and CD8⁺ αβ T cells (1.9-fold; P=0.02), with a shift of the αβ compartment towards CD8⁺ predominance. Total CD4⁺ numbers were unchanged, but both CD4⁺ and CD8⁺ T cells showed increased Eomes expression (P=0.003 and P=0.005), consistent with an activated effector phenotype. The myeloid compartment was remodelled through expansion of CCR2^-^MHCIIlo/^-^ and CCR2^+^ macrophage populations (**Figure 3G**), whereas B cells, dendritic cells and neutrophils were unchanged (**Supplemental Figure 5D**). Intracellular cytokine staining confirmed TNF production capacity across multiple SAN-associated leucocyte lineages in young mice (**Supplemental Figure 6A**). Immunofluorescence in young and aged mice then showed that TNF colocalisation with CD68⁺ macrophages increased with age, whereas colocalisation with CD8⁺ T cells did not (**Supplemental Figure 6B**). Suppression of the pacemaker programme alongside immune activation is therefore conserved in the aged mouse SAN, where it can additionally be linked to measurable pacemaker dysfunction.

### Selective TNF overexpression establishes a proinflammatory nodal microenvironment and impairs pacemaker function

Across human and mouse datasets, TNF emerged as a convergent candidate mediator of age-associated SAN dysfunction. In aged human SAN, TNF–NF-κB signalling was strongly enriched in AREG⁺ macrophages, the population whose proximity to pacemaker cardiomyocytes increased with age, while nearby pacemaker cells showed the greatest loss of pacemaker ion-channel transcripts (**Figure 2B,D,F**; **Supplemental Figure 4D,E**). TNF protein was increased in aged human SAN and colocalised most strongly with macrophages, while aged mouse SAN showed increased *Tnf* expression, TNF-family signalling and TNF protein (**Figure 3F**; **Supplemental Figures 1D, 5A,B, 7A–C**). Human pacemaker cardiomyocytes expressed both TNF receptors (**Supplemental Figure 7D–F**). This cross-species, transcript-to-protein convergence prompted us to test whether TNF is sufficient to impair pacemaker function and whether its inhibition can restore established dysfunction.

To address sufficiency, we generated tamoxifen-inducible *Hcn4-CreERT2;Tnf*^LSL^ mice (HCN4/TNF^glo^), in which TNF expression is induced in HCN4^+^ expressing cells, i.e. pacemaker myocytes (**Figure 4A**; **Supplemental Figure 8A**). This model tests the consequences of a locally generated TNF-rich environment rather than identifying the endogenous cellular source of TNF in the ageing SAN. Selective SAN TNF expression was confirmed at 6 weeks following tamoxifen administration by immunostaining and expression analysis in the SAN (**Figure 4A-C, Supplemental Figure 8C-E**). Cardiac structure and function were preserved on echocardiography, with no overt behavioural or gross physiological abnormalities (**Supplemental Figure 8B**).

**Figure 4.**
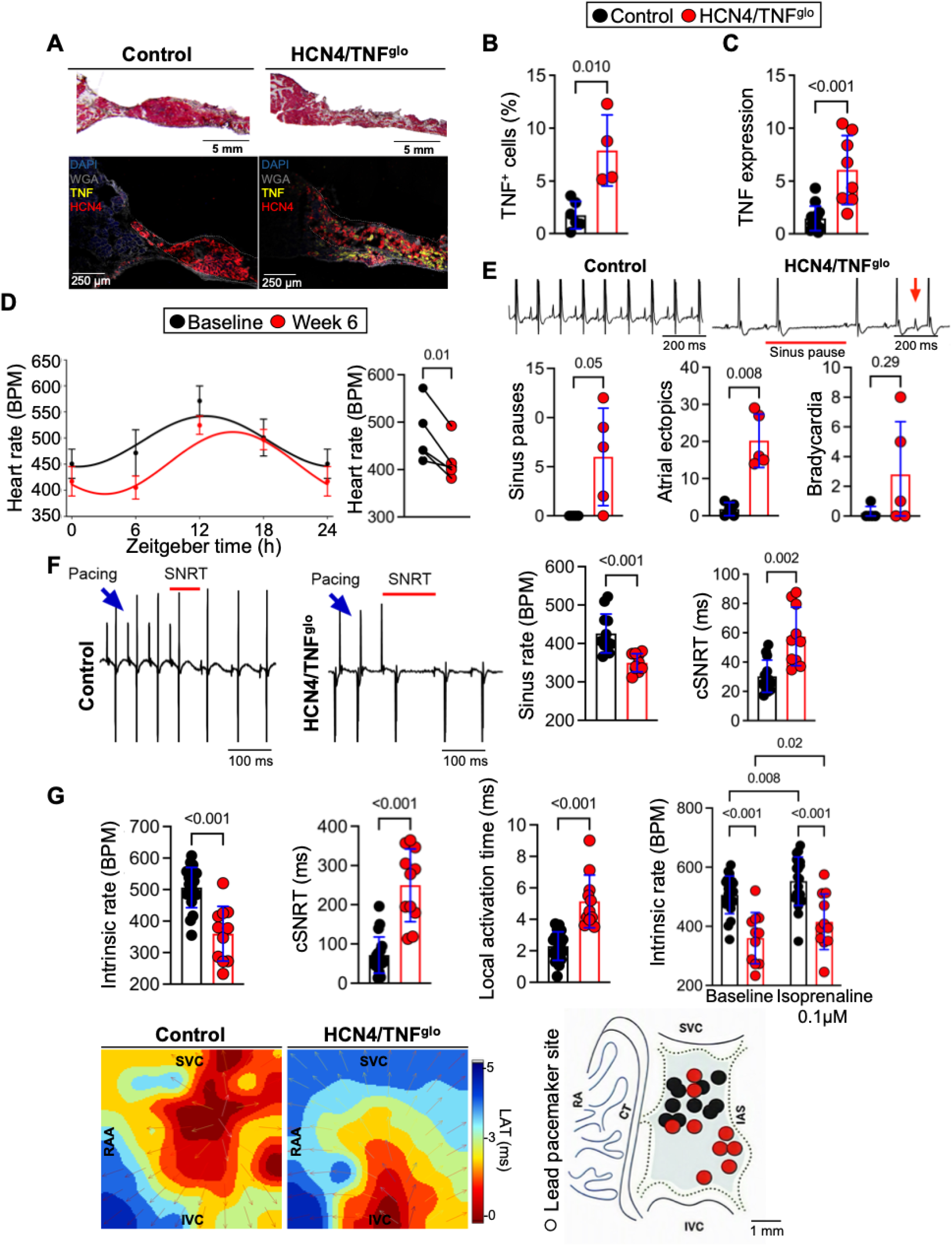
**Selective TNF overexpression in HCN4**⁺ **pacemaker myocytes establishes a proinflammatory nodal microenvironment, impairing pacemaker function**. HCN4-CreERT2 mice were crossed with TNF^loxSTOPlox^ mice to generate inducible HCN4/TNF^glo^ mice. Tamoxifen administration (150 mg kg⁻¹ i.p. on two consecutive days) induced Cre-mediated activation of TNF expression selectively within HCN4⁺ pacemaker cardiomyocytes. TNF^glo^ mice and vehicle-treated HCN4/TNF^glo^ littermates were pooled as a single control group (**Supplemental** Figure 8). **(A)** Representative Masson’s trichrome staining (upper panels) and immunofluorescence images (lower panels) acquired at 20x of SAN tissue from control and tamoxifen-injected HCN4/TNF^glo^ mice, showing TNF (yellow), HCN4 (red), WGA (grey), and DAPI (blue). Scale bars, 5 mm (upper panels) and 250 μm (lower panes). **(B)** Quantification of the percentage of TNF^+^ cells in SAN tissue from control (N=6) and HCN4/TNF^glo^ (N=4) mice. **(C)** Relative TNF mRNA expression in SAN tissue from control (N=8) and HCN4/TNF^glo^ (N=8) mice. Gene expression was normalised to IPO8 and calculated using the 2^−ΔΔCt^ method. **(D)** In vivo ECG telemetry recordings showing circadian heart rate profiles across 24 hours plotted by Zeitgeber time (ZT). Quantification of median active-phase heart rate at baseline and 6 weeks following tamoxifen administration (N=4). Data are presented as mean ± SEM. **(E)** Representative telemetry traces showing sinus pauses (red bars) and atrial ectopic events (red arrows) in control (N=5) and HCN4/TNF^glo^ (N=5) mice. Quantification of sinus pauses, atrial ectopic beats, and bradycardic events obtained from 24-hour telemetry recordings. **(F)** Representative pacing protocols from Langendorff-perfused hearts used to determine sinoatrial node recovery time (SNRT; pacing stimuli (blue arrows), recovery interval (red bars)). Quantification of intrinsic sinus rate and corrected SNRT (cSNRT) in control (N=10) and HCN4/TNF^glo^ (N=10) mice. **(G)** Flexible multi-electrode array (MEA) recordings from isolated SAN preparations of control (N=18) and HCN4/TNF^glo^ (N=11) mice. Quantification of intrinsic SAN rate, cSNRT, local activation time (LAT), and intrinsic rate at baseline and following 0.1 μM isoprenaline. Representative isochronal activation maps showing impulse initiation and propagation, with earliest activation in red and latest activation in blue. Lead pacemaker site localisation is summarised schematically. SVC, superior vena cava; RA, right atrium; CT, crista terminalis; IAS, interatrial septum; IVC, inferior vena cava. Scale bar, 1 mm. Data are presented as mean ± SD with individual biological replicates. P-values computed by Mann–Whitney U test (B, C, E), paired Students t-test (D), Welch’s t-test (F, G), or two-way repeated-measures ANOVA (G: baseline vs isoprenaline).

Six weeks after induction, biotelemetry revealed an SND phenotype with a sustained reduction in heart rate across the 24-h cycle. Cosinor analysis showed a 41.4 BPM reduction in mesor (95% CI 8.4– 74.4; P=0.025), while rhythm amplitude and acrophase were unchanged. Consistent with this, active-phase resting rate was reduced by 55.08 BPM (baseline 473.6±57.8 vs week 6 418.6±40.7 BPM; 95% CI 19.15–91.01; P=0.01; **Figure 4D**). HCN4/TNF^glo^ mice also showed more frequent sinus pauses (7 vs 0; 95% CI 0–12; P=0.05) and atrial ectopic beats (16 vs 2; 95% CI 11–27; P=0.008; **Figure 4E**). *Ex vivo* interrogation confirmed an intrinsic defect in SAN function. Langendorff-perfused HCN4/TNF^glo^ hearts showed a lower intrinsic sinus rate (by 76.4 BPM, 95% CI 41.5-111.3; P<0.001) and prolonged corrected SAN recovery time (by 27.14 ms; 95% CI 12.00-42.28; P=0.002; **Figure 4F**), and MEA mapping showed slowed intercaval conduction with inferior displacement of the leading pacemaker site, consistent with failure of the superior pacemaker complex and recruitment of subsidiary caudal foci as reported in other SND models (**Figure 4G**).^33,34^ Sharp microelectrode recordings in isolated SAN preparations showed a lower spontaneous firing rate (by 98.7 BPM, 95% CI 182.7-14.62; P=0.02), with reduced diastolic depolarisation rate (by 25.4 mV/s, 95% CI 34.95-15.77; P<0.001) and prolonged APD_90_ (by 18.03ms, 95% CL 3.90-32.15; P=0.02; **Figure 5A**), also observed in the peripheral SAN and atrial regions (**Supplemental Figure 9A, B**). Similarly, isolated HCN4/TNF^glo^ myocytes under current clamp showed reduced spontaneous firing rate (by 78.82 BPM, 95% CI 11.9-145.7; P=0.03; **Figure 5B**) and reduced *I*_f_ density (**Figure 5C**, genotype×voltage interaction, P<0.001), including across the physiological diastolic range from −75 to −45 mV (**Supplemental Table 9**). *I*_f_ activation was also shifted by 8.31 mV towards more hyperpolarised potentials (**Figure 5C**, inset), further reducing current availability during diastolic depolarisation, comparable to that reported in pacemaker myocytes from aged mice.^13,43^ Total *I*Ca,L showed no overall genotype effect but a significant genotype×voltage interaction (P=5.05×10⁻⁵⁰), with current density 34–70% lower between −35 and −10 mV. (**Figure 5C**; **Supplemental Table 9**), comparable to that reported in aged SAN myocytes.^44^ Component analysis showed the deficit to be predominantly Ca_V_1.3-mediated (**Figure 5C**, bottom panels): *I*_Cav1.3_ was reduced by 68% at −20 mV and activation shifted 16.77 mV towards more depolarised potentials, whereas *I*_Cav1.2_ did not exhibit voltage-specific differences or change in activation. Together, these findings indicate that TNF induction reproduces key ionic features of the aged SAN, with the L-type Ca²⁺-current deficit arising predominantly through Ca_V_1.3.

**Figure 5.**
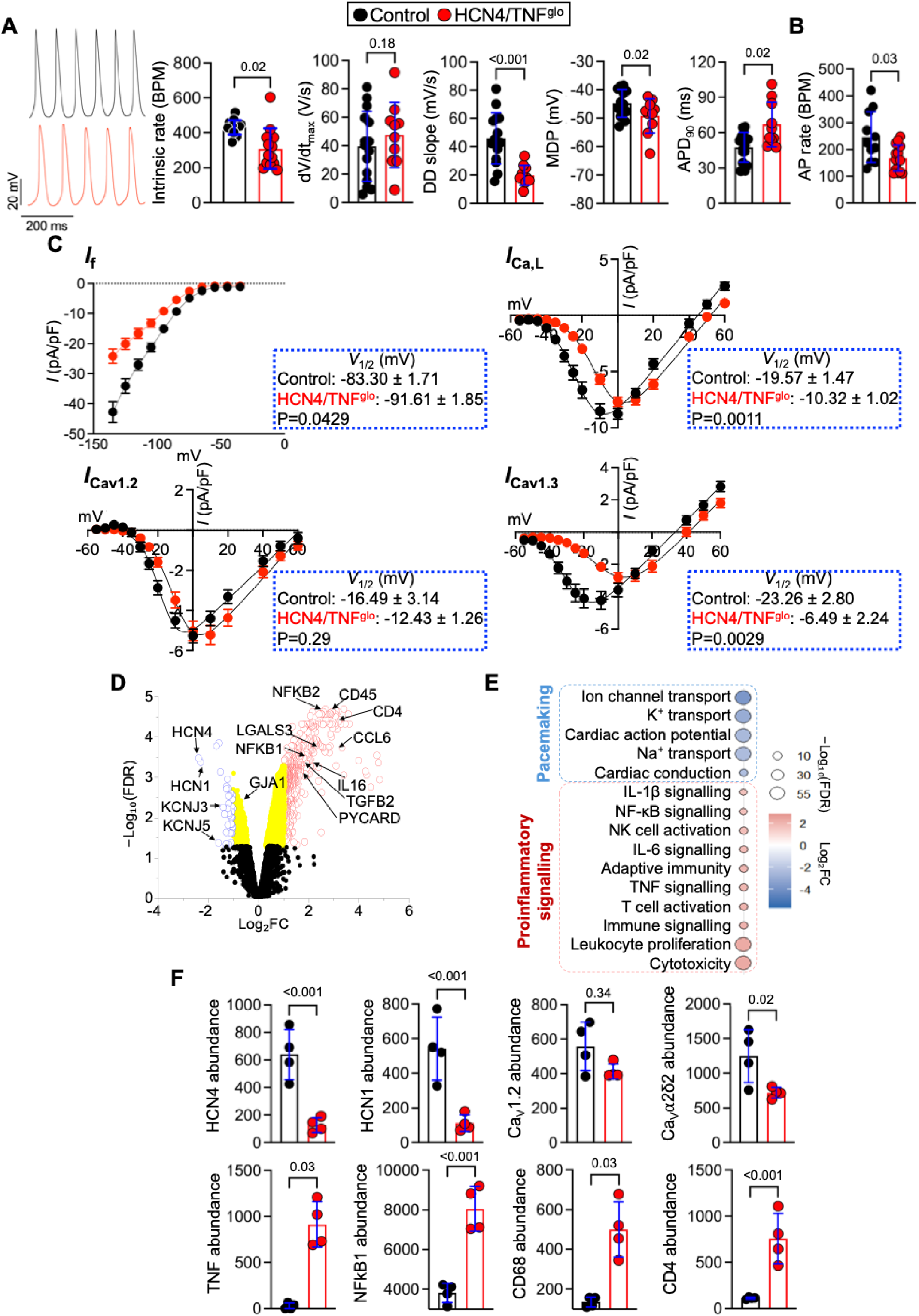
TNF overexpression impairs intrinsic pacemaker function and remodels the SAN proteome. **(A)** Representative spontaneous intracellular action potentials recorded from central SAN myocytes from control (n=53/N=15) and HCN4/TNF^glo^ (n=40/N=10) mice. Quantification of intrinsic firing rate, maximal upstroke velocity (dV/dt_max_), diastolic depolarisation (DD) slope, maximum diastolic potential (MDP), and action potential duration at 90% repolarisation (APD_90_). Each dot represents the mean value obtained from averaged impalements per animal. **(B)** Quantification of spontaneous action potential firing rate from perforated patch clamp recordings from control (N=5) and HCN4/TNF^glo^ (N=7) mice. **(C)** Current-voltage relationships of the hyperpolarisation-activated funny current (*I*_f_), total L-type Ca^2+^ current (*I*_Ca,L_), and the Ca_V_1.2-(*I*_CaV1.2_) and Ca_V_1.3 (*I*_CaV1.3_)-mediated components of *I*_Ca,L_, measured by whole-cell patch clamp in isolated SAN myocytes from control and HCN4/TNF^glo^ mice. Current amplitudes were normalised to membrane capacitance and are expressed as pA/pF. Insets show the voltage of half-maximal activation (*V*_1/2_; mean±SEM) and corresponding *P*-values from nested two-tailed *t*-tests. Current-voltage data are mean±SEM. Current-voltage relationships were analysed using linear mixed-effects models with genotype, test voltage, and their interaction as fixed effects, repeated measurements across voltage within cells, and random intercepts for animals and cells nested within animals. Full voltage-specific statistics are provided in **Supplemental Table 9**. For *I*_f_: control, n=46 cells/N=9 mice; HCN4/TNF^glo^, n=54 cells/N=13 mice. Mixed-effects analysis: genotype, P<0.001; voltage, P<0.001; genotype×voltage interaction, P<0.001. For total *I*_Ca,L_: control, n=27 cells/N=5 mice; HCN4/TNF^glo^, n=35 cells/N=6 mice. Mixed-effects analysis: genotype, P=.0809; voltage, P<0.001; genotype×voltage interaction, P<0.001. For *I*_Cav1.2_: control, n=23 cells/N=4 mice; HCN4/TNF^glo^, n=27 cells/N=5 mice. Mixed-effects analysis: genotype, P=.9895; voltage, P<0.001; genotype×voltage interaction, P<0.001. For *I*_Cav1.3_: control, n=23 cells/N=4 mice; HCN4/TNF^glo^, n=29 cells/N=5 mice. Mixed-effects analysis: genotype, P=.0170; voltage, P<0.001; genotype×voltage interaction, P<0.001. **(D)** Proteomic profiling of SAN tissue from control (N=4) and HCN4/TNF^glo^ (N=4) mice. Volcano plot showing differentially expressed proteins identified by mass spectrometry, with selected pacemaker-, ion channel-, inflammatory-, and immune-associated proteins highlighted. Upregulated proteins are shown in red, downregulated proteins in blue, proteins exceeding the log₂FC threshold alone in yellow, and non-significant proteins in black. **(E)** Gene set enrichment analysis of differentially expressed proteins (Gene ontology database). Dot colour denotes log_2_FC and dot size represents −log10(FDR). Pacemaking-and pro-inflammatory signalling-related pathways highlighted. **(F)** Normalised protein abundance of HCN4, HCN1, Cav1.2, Ca_V_α2δ2, TNF, NFKB1, CD68, and CD4 in SAN tissue from control (N=4) and HCN4/TNF^glo^ (N=4) mice. Data are presented as mean ± SD with individual biological replicates. P-values computed by nested t-test (A), Mann– Whitney U test (B), mixed-effect analysis (C), Limma test with Benjamini–Hochberg correction (D, F) or permutation testing with Benjamini–Hochberg correction (E).

Unbiased proteomic profiling revealed extensive molecular remodelling following TNF induction. Of 5,108 proteins detected, 2,010 were differentially abundant at FDR ≤ 0.05, with 1,558 increased and 452 decreased (**Figure 5D**), and TNF itself increased 34.7-fold (**Figure 5F**). Within the pacemaker programme, HCN4 and HCN1 were reduced 5.2-fold and 4.9-fold, consistent with the reduction in *I*_f_, alongside reductions in GIRK1, GIRK4, SERCA2a, calsequestrin-2 and the auxiliary Ca²⁺-channel subunit Ca_v_α2δ2. Ca_v_1.2 abundance was unchanged, consistent with the absence of an overall genotype effect on *I*_Cav1.2_, whereas Ca_v_1.3 was not detected. These changes occurred alongside marked immune remodelling, with increased CD68, CD4 and NF-κB1 and the chemoattractants CCL6, CCL9, IL-16 and IL-33 (**Figure 5F**, **Supplemental Figure 9E**). Immunostaining showed concordant increases in CD45⁺ leukocytes, AREG⁺CD68⁺ macrophages and CD8⁺ and CD4⁺ T cells (**Supplemental Figure 9F,G**), while fibrillar collagens were unchanged (**Supplemental Figure 9H**). Pathway analysis reinforced this convergence, showing loss of ion channel gene regulatory programmes and gain of inflammatory, chemotactic and phagocytic pathways (**Figure 5E**; **Supplemental Figure 9C,D**). Taken together, a TNF-rich SAN microenvironment is sufficient to recapitulate the multi-scale phenotype of age-related SND, from loss of the molecular machinery of automaticity through to SND in vivo, without fibrosis or overt myocardial disease. To investigate potential downstream mechanisms, PROGENy analysis was performed and identified NF-κB as the pathway showing the greatest predicted increase in activity in aged mouse SAN (**Supplemental Figure 10A,B**), with increased NF-κB p65-positive area and NFKB1 and TNF transcript expression in aged human SAN (**Supplemental Figure 10F**). Conserved predicted NF-κB binding motifs were present at the HCN4, HCN1, CACNA1D and TBX3 loci in both species, including one within the HCN4 proximal promoter (**Supplemental Figure 10C**). In TNF-exposed mouse SAN preparations, IKKβ inhibition increased *Hcn4*, *Hcn1* and *Tbx3* expression relative to vehicle, whereas *Cacna1d* expression was unaffected (**Supplemental Figure 10E**).

### TNF induction suppresses HCN4 in the human sinoatrial node

To determine whether TNF induces comparable remodelling in human tissue, we established organotypic cultures of 500 μm SAN slices from seven donor hearts spanning a range of ages and clinical backgrounds (**Supplemental Table 1**) and recorded spontaneous action potentials using intracellular sharp microelectrodes (**Figure 6A**). Freshly prepared slices yielded 18 impalements across the seven donors. Spontaneous cycle length and resting membrane potential were consistent with published values for adult human SAN ^45,46^ and isoprenaline (100 nM) shortened cycle length by 47.9±25.56% indicating preserved β-adrenergic responsiveness (**Supplemental Table 2)**.

**Figure 6.**
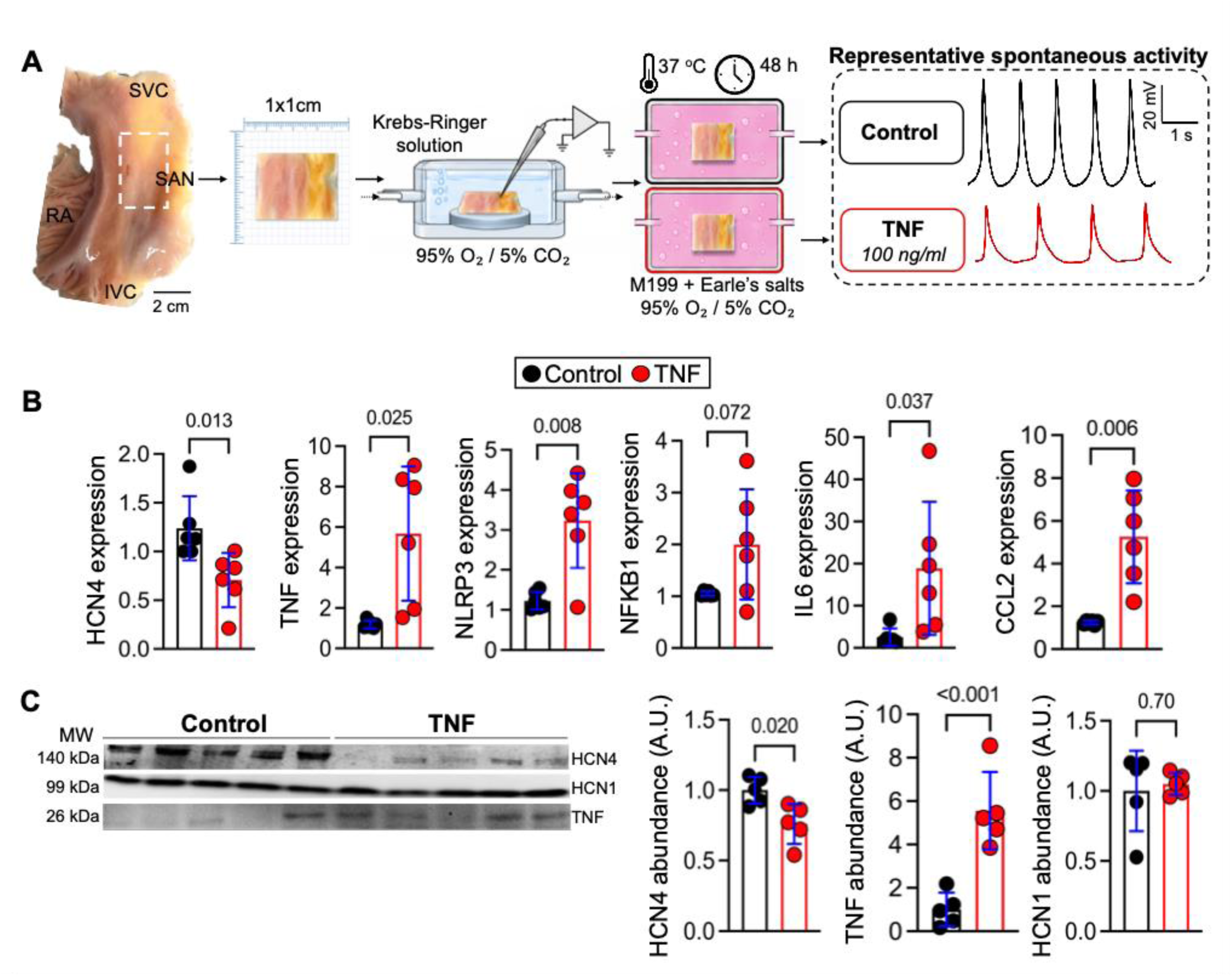
TNF activates inflammatory signalling and suppresses HCN4 in human SAN slices. **(A)** Schematic of human SAN slice preparation. The SAN region was dissected from the superior right atrial wall and sectioned into 500 μm-thick tissue slices. Slices were maintained in culture for 48h at 37°C in M199 + Earle’s salts with either control medium or TNF (100 ng/mL). Representative spontaneous action potential recordings. Scale bar, 2 cm. SVC, superior vena cava; RA, right atrium; IVC, inferior vena cava. Representative recordings illustrate persistence of spontaneous activity after 48 h culture and were not used to infer TNF-specific electrophysiological remodelling. **(B)** Relative mRNA expression of HCN4, TNF, NLRP3, NFKB1, IL6 and CCL2 in human SAN slices following 48 h culture from control (N=6) or TNF (N=6). Gene expression was normalised to GAPDH and calculated using the 2^-ΔΔCt^ method. **(C)** Representative western blots and quantification of stain-free total protein-normalised HCN4, TNF, and HCN1 protein abundance in human SAN slices following 48 h culture from control (N=6) or TNF (N=6). Data are presented as mean ± SD with individual donors. P-values computed by paired students t-test (B, C).

Slices from the same donor were then cultured for 48 h with recombinant TNF (100 ng/mL) or in parallel as time-matched controls. Spontaneous activity persisted throughout 48 h of organotypic culture (**Supplemental Figure 11A–D**), establishing the functional viability of the preparation over the experimental interval. TNF increased CCL2, IL6 and NLRP3 transcript abundance and there was a trend towards increase in NF-κB (**Figure 6B**). It also reduced HCN4 transcript abundance by 43% (P=0.013) and HCN4 protein abundance by 24% (P=0.020; **Figure 6C**), recapitulating the loss identified in the murine models. HCN1, Cx45, and KCNH2 protein abundance were unchanged (**Figure 6C, Supplemental Fig 11E**) whereas and Ca_v_1.2 and Ca_v_1.3 could not be reliably detected in western blot experiments with the available antibodies. Cleaved caspase-3 positivity did not differ between paired control and TNF-treated slices, nor between young and aged donor SAN tissue (**Supplemental Figures 11F and 1G**), arguing against apoptotic cell loss as an explanation for the reduction in HCN4. These experiments nevertheless show that TNF activates inflammatory signalling and suppresses HCN4 in living adult human SAN tissue, conserving a central molecular component of the murine remodelling.

### Systemic TNF inhibition restores intrinsic SAN function in aged mice

Having established that TNF is sufficient to induce SAN dysfunction, we investigated whether its neutralisation could reverse established dysfunction in aged mice. Aged mice (22 months) received etanercept (2.5 mg kg⁻¹, i.p., twice weekly for 8 weeks) and were compared with vehicle-treated aged controls and young mice (2 months). On surface ECG, ageing reduced heart rate and prolonged PR, QRS and QTc intervals, with a trend towards RR prolongation (**Figure 7A**). Etanercept shortened the RR and PR intervals (by 30 ms and 5.40 ms vs aged controls, 95% CI 8.3-52 and 0.51-10.29), restoring heart rate towards that of young mice. QRS was not changed; QTc was partially improved but not normalised. *Ex vivo*, cSNRT did not differ significantly between groups, while age-associated prolongation of WBCL and AVERP persisted despite treatment (**Supplemental Figure 12D**).

**Figure 7.**
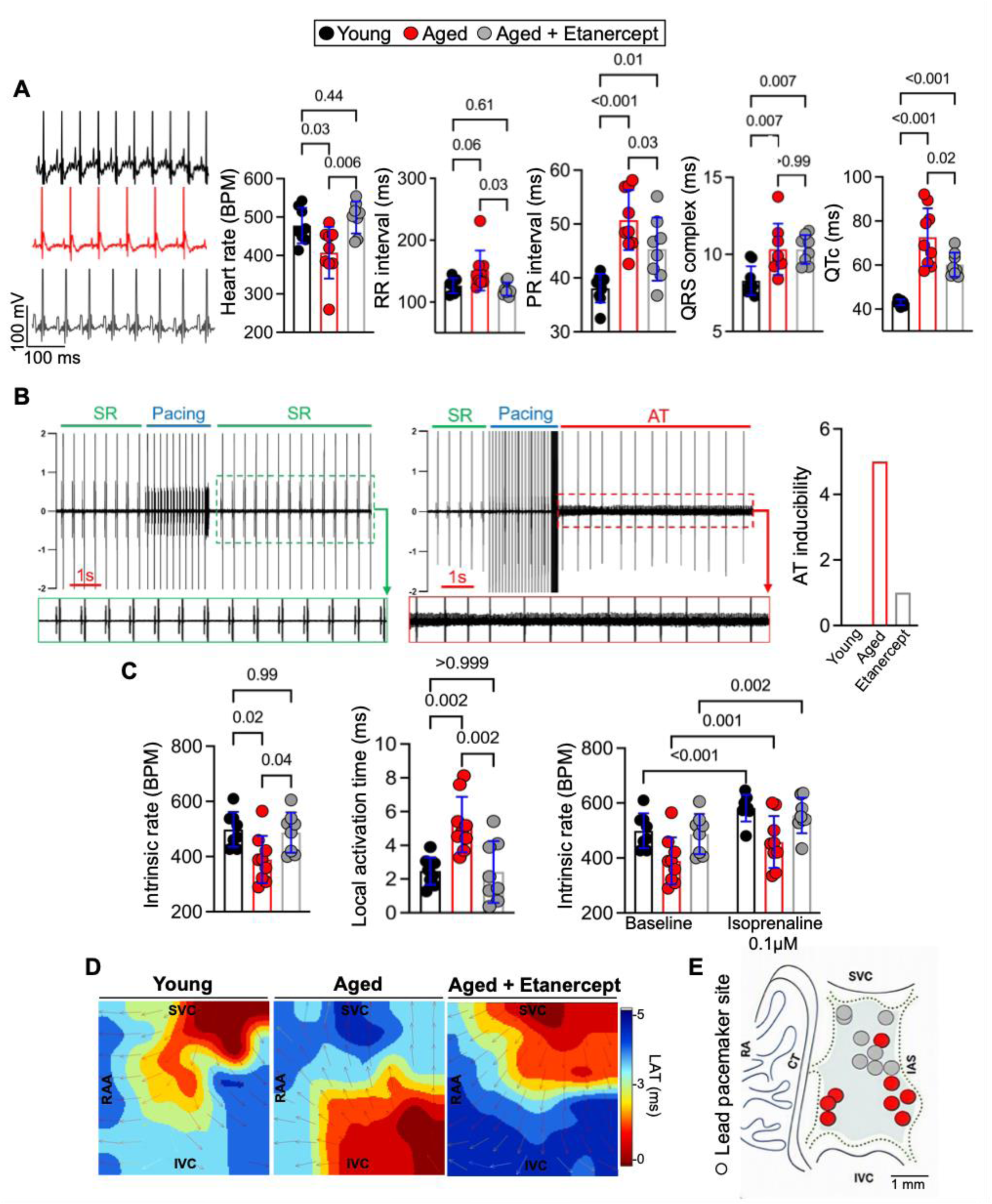
Pharmacological TNF blockade reverses age-associated SND. **(A)** Representative anaesthetised ECG recordings from young (N=8), aged (N=8), and aged + etanercept (N=8) mice. Quantification of heart rate, RR interval, PR interval, QRS complex, and corrected QT interval (QTc). **(B)** Ex vivo Langendorff assessment of atrial rhythm stability. Representative electrogram traces showing sinus rhythm (SR) maintained following programmed electrical stimulation and atrial tachycardia (AT) inducibility with S1-S11 pacing. Quantification of AT inducibility in young (N=8), aged (N=8), and aged + etanercept (N=8) mice. **(C)** Flexible multi-electrode array (MEA) recordings from isolated SAN preparations. Quantification of intrinsic SAN rate, local activation time (LAT), and intrinsic rate at baseline and following 0.1 μM isoprenaline in young (N=8), aged (N=8), and aged + etanercept (N=8) mice. **(D)** Representative isochronal activation maps from young, aged, and aged + etanercept SAN preparations, with earliest activation shown in red and latest activation sites in blue. **(E)** Lead pacemaker site localisation summarised schematically for aged (red) and aged + etanercept (grey) SAN. SVC, superior vena cava; RA, right atrium; CT, crista terminalis; IAS, interatrial septum; IVC, inferior vena cava. Scale bar, 1 mm. Data are presented as mean ± SD with individual biological replicates. Pvalues computed by one-way ANOVA with Šídák’s multiple-comparisons test (A, C) or two-way repeated-measures ANOVA (C: baseline vs isoprenaline).

Interestingly, etanercept also reduced the inducibility of atrial arrhythmia by extrastimulus pacing in Langendorff-perfused hearts from 5 of 8 aged controls to 1 of 8 treated animals (**Figure 7B**). To test whether recovery occurred within the SAN itself, we examined isolated preparations. Ageing reduced intrinsic sinus rate, prolonged local activation time, slowed intercaval conduction, displaced the leading pacemaker site inferiorly and normalised the heart rate achieved during β-adrenergic stimulation with isoprenaline (**Figure 7C-E**; **Figure 3A,B**). Each was reversed. Etanercept raised intrinsic rate by 97.65 BPM over aged controls (95% CI 3.36-191.9; P=0.04) and shortened local activation time by 2.80 ms (95% CI 0.95-4.65; P=0.002), restoring both to values indistinguishable from young mice (**Figure 7C**), returned the leading pacemaker site to a superior position **(Figure 7E**).Isoprenaline increased spontaneous rate in all groups, and although the magnitude of the increase was similar in aged and treated preparations, the lower stimulated rate in aged mice was restored by etanercept (458.0±90.5 to 554.8±67.0 BPM; P=0.03), to a value indistinguishable from young mice (P=0.864; **Figure 7C**).These functional gains were accompanied by restoration of the pacemaker apparatus and reduced SAN inflammation; Etanercept increased HCN4 transcript abundance by 79% whereas Ca_v_1.3 other nodal and ion channel transcripts were unchanged (**Supplemental Figure 12E**). Treatment reduced *TNF* expression, while *Nfkbia* expression showed a non-significant increase (**Figure 8A**), consistent with attenuation of the inflammatory programme, and lowered TNF⁺ cell fraction (from 16.62±1.28% to 3.65±1.96%, 95% CI 9.25-16.70; P<0.001), AREG⁺CD68⁺ macrophages accumulation (from 78.59±11.57% to 22.40±7.04%, 95% CI 35.22-77.16; P<0.001), and CD45^+^ leukocyte infiltration (74±29.10% to 15.33±9.07%, 95% CI 11.46-105.9; P=0.02); CD8⁺ and CD4⁺ cell fractions and collagen area showed concordant but non-significant reductions (**Figure 8B,C**). The benefits extended beyond the SAN. Etanercept attenuated hepatic inflammatory gene expression (**Supplemental Figure 12F**), markedly reduced frailty, which was also apparent on gross inspection (**Supplemental Figure 12A**) and prevented the mortality observed in aged controls during treatment. All treated mice reached the mandated 24-month endpoint, compared with only 50% of vehicle-treated aged mice (log-rank P=0.007, **Figure 8D, E**). Left ventricular ejection fraction and fractional shortening were unchanged (**Supplemental Figure 12B,C**), arguing against improved global systolic function as the explanation for recovery of intrinsic pacemaker activity. Together, these findings demonstrate that TNF blockade reverses established molecular and functional features of SAN ageing while conferring broader frailty and survival benefits in aged animals.

**Figure 8.**
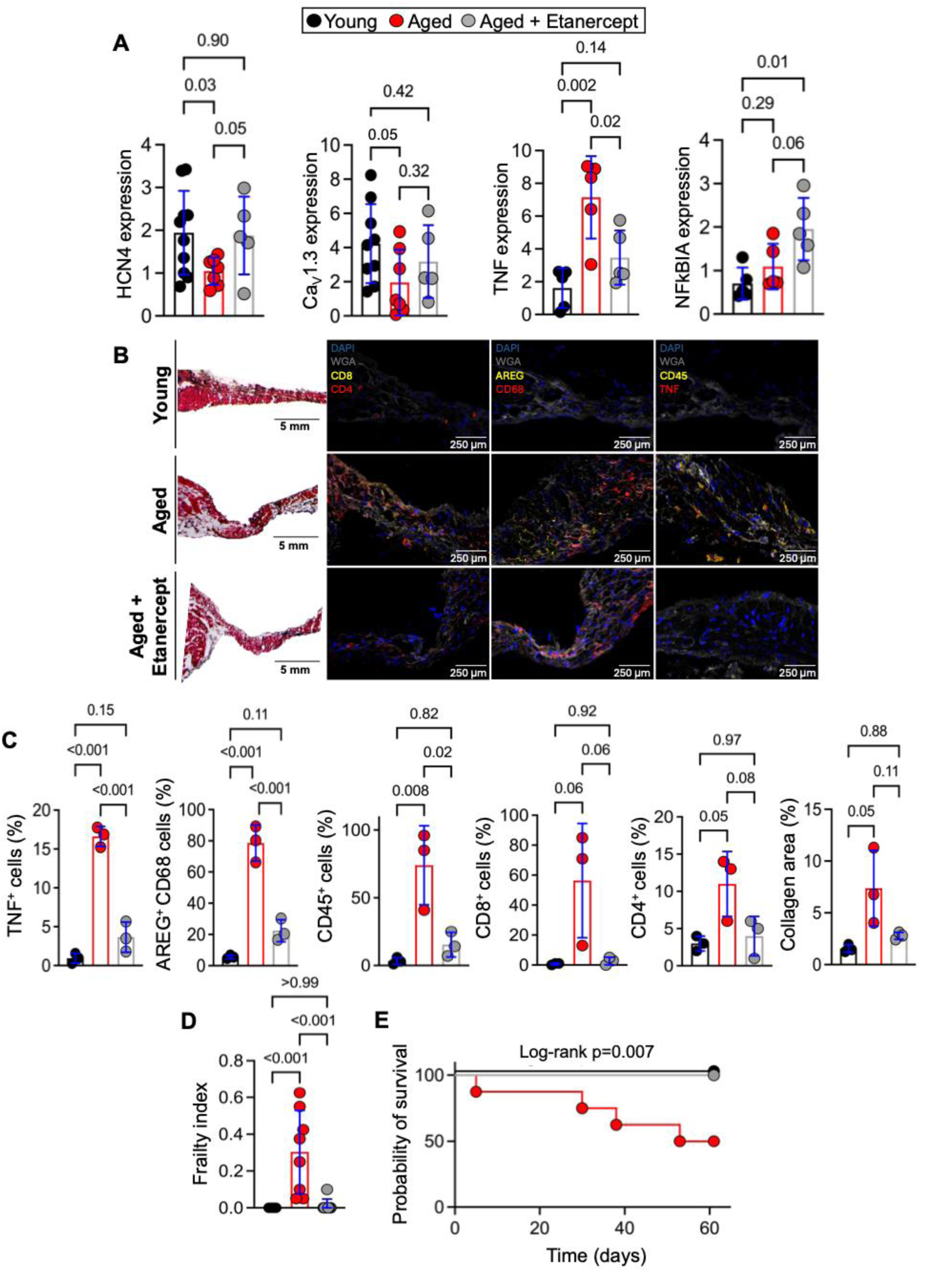
Pharmacological TNF blockade attenuated age-associated inflammatory remodelling of the SAN. **(A)** Relative mRNA expression of HCN4, CACNA1D (Ca_V_1.3), TNF, and NFkBIA in SAN tissue from young (N=8), aged (N=8), and aged + etanercept (N=8) mice. Gene expression was normalised to IPO8 and calculated using the 2^−ΔΔCt^ method. **(B)** Representative Masson’s trichrome staining and immunofluorescence images acquired at 20x of SAN tissue from young, aged, and aged + etanercept mice showing CD8 (yellow) and CD4 (red), AREG (yellow) and CD68 (red), CD45 (yellow) and TNF (red), WGA (grey), and DAPI (blue). Scale bars, 5 mm (left) 250μm (right). **(C)** Quantification of the percentage of TNF^+^ cells, AREG⁺CD68⁺ macrophages, CD45⁺ leukocytes, CD8⁺ T cells, CD4⁺ T cells, and collagen area fraction in young (N=3), aged (N=3), and aged + etanercept (N=3) mice. **(D)** Frailty index scores in young (N=8), aged (N=8), and aged + etanercept (N=8) mice during the treatment period. **(E)** Kaplan-Meier survival curves from young (N=8), aged (N=8), and aged + etanercept (N=8) mice during the treatment period. Survival distributions were compared using log-rank (Mantel-Cox) test. Data are presented as mean ± SD with individual biological replicates. P-values computed by one-way ANOVA with Šídák’s multiple-comparisons test (A, C, D) or log-rank (Mantel-Cox) test (E).

## Discussion

This study identifies inflammatory remodelling of the SAN microenvironment as an important component of human pacemaker ageing. To our knowledge, it provides the first age-resolved spatial analysis of the human SAN at cellular resolution and the first molecular characterisation of pacemaker ion channel remodelling in the ageing human node. Within this broader inflammatory response, convergent human and mouse data identify TNF signalling as a functionally important and therapeutically modifiable contributor to suppression of the pacemaker programme and nodal dysfunction.

Age-dependent remodelling of pacemaker ion channels has been extensively defined in animals^8–15,41^ but its relevance to human SAN ageing has remained uncertain. Here, aged human pacemaker cardiomyocytes showed coordinated suppression of genes required for automaticity, including HCN4, HCN1, CACNA1D, CACNA1G, and SLC8A1, while transitional cardiomyocytes showed broader loss of ion channel, Ca²⁺-handling, and connexin programmes. Spatial analysis revealed that this de-specialisation was not uniform. AREG⁺ macrophages shifted from relative exclusion from pacemaker cardiomyocytes in young tissue to enrichment around them with age, and neighbouring pacemaker cells showed the clearest loss of HCN4, HCN1, and TBX3 together with inflammatory and matrix-associated signalling. This extends the concept of specialised cardiac immune niches^7^ to SAN ageing and suggests that pacemaker cell identity is shaped by the local cellular environment. Amphiregulin, which can regulate Cx43^47^ provides one possible paracrine mediator, although whether it is causal or simply marks the macrophage state associated with this niche remains unresolved. This is distinct from direct electrical coupling, which we excluded by the complete absence of Cx43 gap junctions between macrophages and HCN4⁺ pacemaker cells, consistent with our previous work^48^ and that of others.^7^

The enrichment of TNF signalling within these niches raised the more direct question of whether cytokine elaboration could itself drive remodelling of the SAN pacemaker complex. Selective TNF expression in HCN4⁺ myocytes reproduced the cardinal features of clinical SND, including bradycardia, sinus pauses, and conduction slowing, in structurally normal hearts, establishing that a TNF-rich nodal environment is sufficient to impair pacemaker function. The underlying ion channel changes involved both altered abundance and biophysical regulation. Reduced HCN4 abundance provides a plausible basis for the lower *I*_f_ density, while the hyperpolarising shift in activation and the marked depolarising shift in *I*_Cav1.3_ activation would each limit inward current across the diastolic voltage range, indicating that TNF acts through both transcriptional and biophysical mechanisms. Our additional analyses implicate IKK/NF-κB signalling in suppression of the pacemaker programme, with IKK inhibition increasing *Hcn4, Hcn1 and Tbx3* expression in TNF-exposed SAN preparations (**Supplemental Figure 10**). The accompanying shifts in *I*_f_ and Ca_V_1.3 voltage dependence suggest additional effects on channel gating, potentially mediated by TNF-responsive pathways such as PI3K–Akt, which can regulate HCN activation and L-type Ca²⁺-channel activity.^49–51^ The intervening signalling steps, and the mechanisms underlying these biophysical changes, remain to be established.

The conservation of this response in part at least in human tissue strengthens its relevance to human SAN ageing. In viable adult human SAN slices, TNF activated inflammatory signalling and reduced HCN4 transcript and protein, reproducing a central molecular feature observed in the aged human SAN and in the mouse models. These experiments do not establish downstream functional consequences but they show that native adult human nodal tissue retains a conserved molecular response to TNF and provide an experimentally accessible bridge between the spatial observations in human tissue and mechanistic studies in mice.

Fibrosis accompanied ageing in the human SAN, but is unlikely to fully account for the associated functional decline. The largest quantitative histological series found no consistent increase in SAN collagen across adult life,^6^ and contemporary reviews similarly conclude that the contribution of fibrosis to age-related intrinsic pacemaker slowing remains uncertain.^52^ Our mouse experiments provide a functional distinction: TNF induction produced pronounced electrophysiological dysfunction without overt collagen accumulation, whereas etanercept restored automaticity, conduction, and chronotropic responsiveness in aged mice without a detectable reduction in ECM deposition. These findings do not exclude a contribution of fibrosis to the aged SAN substrate, but demonstrate that substantial deterioration and recovery of pacemaker function can occur independently of detectable ECM remodelling.

The most clinically relevant finding is that established age-associated SAN dysfunction retains substantial capacity for functional recovery. Earlier studies catalogued age-dependent defects in nodal ion channels, Ca²⁺ signalling and conduction, but whether an already dysfunctional node retained the capacity to recover was unknown. In aged mice with established disease, etanercept restored intrinsic rate, activation across the node, superior pacemaker site recruitment and normalised rate during β-adrenergic stimulation. Recovery persisted in isolated SAN preparations and occurred without improvement in global left ventricular systolic function, consistent with intrinsic recovery accompanied by restoration of HCN4 and attenuation of local inflammatory signalling and AREG⁺ macrophage accumulation. Etanercept also reduced frailty and extended survival, placing the SAN within the broader emerging evidence that selective cytokine inhibition can improve healthspan in ageing.^53^ These findings define a therapeutic direction distinct from the negative anti-TNF heart-failure trials,^54,55^ which targeted heterogeneous populations with advanced systolic dysfunction, whereas the phenotype rescued here is a localised disorder of automaticity in hearts with preserved systolic function. As with IL-1β blockade in atherosclerosis,^56^ these results argue for precision approaches directed at the inflammatory microenvironment of the SAN rather than broad TNF antagonism. Circulating TNF already tracks SND severity and predicts progression to tachycardia-bradycardia syndrome in patients, identifying it as a candidate stratification biomarker.^57^ The next challenge is to test this prospectively and to develop pathway-selective or SAN-directed strategies capable of modifying progression before pacing becomes necessary.

The following limitations apply: The human cohort was necessarily modest and cross-sectional, and spatial proximity cannot establish the direction or mediators of communication between macrophages and pacemaker cells. Spatial profiling used a targeted gene panel and may therefore have missed additional pathways relevant to SAN ageing. The HCN4/TNF^glo^ model establishes the sufficiency of local TNF elevation but not the necessity of endogenous TNF signalling during SAN ageing. The relative contributions of individual TNF-producing populations remain to be established, while systemic etanercept treatment cannot distinguish local nodal from broader anti-inflammatory effects. The persistence of age-associated AV nodal conduction and refractory abnormalities despite etanercept treatment suggests that the mechanisms governing AVN ageing may differ from those contributing to SAN dysfunction, warranting further investigation. Finally, the durability of recovery after treatment withdrawal and its translation to human SND remain to be established.

## Conclusion

In sum this study identifies a modifiable inflammatory-electrical component within the ageing human SAN and establishes TNF-dependent signalling as a candidate route towards disease modification in age-associated SND.

## Data availability statement

Data supporting the findings of this study are available from the corresponding author upon reasonable request.

## References

1 Defaye, P. et al. Cardiac pacing and lead devices management: 25 years of research at EP Europace journal. Europace 25 (2023). 10.1093/europace/euad202

2 Raatikainen, M. J. P. et al. A Decade of Information on the Use of Cardiac Implantable Electronic Devices and Interventional Electrophysiological Procedures in the European Society of Cardiology Countries: 2017 Report from the European Heart Rhythm Association. Europace 19, ii1–ii90 (2017). 10.1093/europace/eux258

3 Gutierrez-Carretero, E. et al. Mortality and Costs of Cardiac Implantable Electronic Device (CIED) Infections According to the Therapeutic Approach: A Single-Center Cohort Study. Microorganisms 12 (2024). 10.3390/microorganisms12030537

4 Mlynarska, A., Mlynarski, R. & Golba, K. S. Influence of frailty on the quality of life patients qualified for pacemaker implantation. J Clin Nurs 27, 555–560 (2018). 10.1111/jocn.13897

5 Bhatia, N. & El-Chami, M. Leadless pacemakers: a contemporary review. J Geriatr Cardiol 15, 249– 253 (2018). 10.11909/j.issn.1671-5411.2018.04.002

6 Bois, M. C. et al. Age-related histologic features of the sinoatrial node from normal human hearts during the first 10 decades of life: a study of 200 cases. Cardiovasc Pathol 52, 107327 (2021). 10.1016/j.carpath.2021.107327

7 Kanemaru, K. et al. Spatially resolved multiomics of human cardiac niches. Nature 619, 801–810 (2023). 10.1038/s41586-023-06311-1

8 Hao, X. et al. TGF-beta1-mediated fibrosis and ion channel remodeling are key mechanisms in producing the sinus node dysfunction associated with SCN5A deficiency and aging. Circ Arrhythm Electrophysiol 4, 397–406 (2011). 10.1161/CIRCEP.110.960807

9 Jones, S. A., Lancaster, M. K. & Boyett, M. R. Ageing-related changes of connexins and conduction within the sinoatrial node. J Physiol 560, 429–437 (2004). 10.1113/jphysiol.2004.072108

10 Tellez, J. O. et al. Ageing-dependent remodelling of ion channel and Ca2+ clock genes underlying sino-atrial node pacemaking. Exp Physiol 96, 1163–1178 (2011). 10.1113/expphysiol.2011.057752

11 Yanni, J., Tellez, J. O., Sutyagin, P. V., Boyett, M. R. & Dobrzynski, H. Structural remodelling of the sinoatrial node in obese old rats. J Mol Cell Cardiol 48, 653–662 (2010). 10.1016/j.yjmcc.2009.08.023

12 Jones, S. A., Boyett, M. R. & Lancaster, M. K. Declining into failure: the age-dependent loss of the L-type calcium channel within the sinoatrial node. Circulation 115, 1183–1190 (2007). 10.1161/CIRCULATIONAHA.106.663070

13 Larson, E. D., St Clair, J. R., Sumner, W. A., Bannister, R. A. & Proenza, C. Depressed pacemaker activity of sinoatrial node myocytes contributes to the age-dependent decline in maximum heart rate. Proc Natl Acad Sci U S A 110, 18011–18016 (2013). 10.1073/pnas.1308477110

14 Zhang, J. et al. MicroRNA-1976 regulates degeneration of the sinoatrial node by targeting Ca(v)1.2 and Ca(v)1.3 ion channels. J Mol Cell Cardiol 134, 74–85 (2019). 10.1016/j.yjmcc.2019.06.018

15 Liu, J. et al. Age-associated abnormalities of intrinsic automaticity of sinoatrial nodal cells are linked to deficient cAMP-PKA-Ca(2+) signaling. Am J Physiol Heart Circ Physiol 306, H1385–1397 (2014). 10.1152/ajpheart.00088.2014

16 Franceschi, C. et al. Inflamm-aging. An evolutionary perspective on immunosenescence. Ann N Y Acad Sci 908, 244–254 (2000). 10.1111/j.1749-6632.2000.tb06651.x

17 Volpato, S. et al. Cardiovascular disease, interleukin-6, and risk of mortality in older women: the women’s health and aging study. Circulation 103, 947–953 (2001). 10.1161/01.cir.103.7.947

18 Soysal, P. et al. Inflammation and frailty in the elderly: A systematic review and meta-analysis. Ageing Res Rev 31, 1–8 (2016). 10.1016/j.arr.2016.08.006

19 Liberale, L., Montecucco, F., Tardif, J. C., Libby, P. & Camici, G. G. Inflamm-ageing: the role of inflammation in age-dependent cardiovascular disease. Eur Heart J 41, 2974–2982 (2020). 10.1093/eurheartj/ehz961

20 Liberale, L. et al. Inflammation, Aging, and Cardiovascular Disease: JACC Review Topic of the Week. J Am Coll Cardiol 79, 837–847 (2022). 10.1016/j.jacc.2021.12.017

21 Spinelli, V. et al. Interleukin-6 Modulates the Expression and Function of HCN Channels: A Link Between Inflammation and Atrial Electrogenesis. Int J Mol Sci 25 (2024). 10.3390/ijms252212212

22 Lee, S. H. et al. Tumor necrosis factor-alpha alters calcium handling and increases arrhythmogenesis of pulmonary vein cardiomyocytes. Life Sci 80, 1806–1815 (2007). 10.1016/j.lfs.2007.02.029

23 El Khoury, N., Mathieu, S. & Fiset, C. Interleukin-1beta reduces L-type Ca2+ current through protein kinase Cɛ activation in mouse heart. J Biol Chem 289, 21896–21908 (2014). 10.1074/jbc.M114.549642

24 Kawada, H. et al. Tumor necrosis factor-alpha downregulates the voltage gated outward K+ current in cultured neonatal rat cardiomyocytes: a possible cause of electrical remodeling in diseased hearts. Circ J 70, 605–609 (2006). 10.1253/circj.70.605

25 Monnerat-Cahli, G. et al. Toll-like receptor 4 activation promotes cardiac arrhythmias by decreasing the transient outward potassium current (Ito) through an IRF3-dependent and MyD88-independent pathway. J Mol Cell Cardiol 76, 116–125 (2014). 10.1016/j.yjmcc.2014.08.012

26 Saraf, A. et al. Functional and molecular effects of TNF-alpha on human iPSC-derived cardiomyocytes. Stem Cell Res 52, 102218 (2021). 10.1016/j.scr.2021.102218

27 De Jesus, N. M. et al. Antiarrhythmic effects of interleukin 1 inhibition after myocardial infarction. Heart Rhythm 14, 727–736 (2017). 10.1016/j.hrthm.2017.01.027

28 Chien, S. J., Liang, C. D., Lin, I. C., Lin, Y. J. & Huang, C. F. Myocarditis complicated by complete atrioventricular block: nine years’ experience in a medical center. Pediatr Neonatol 49, 218–222 (2008). 10.1016/S1875-9572(09)60014-0

29 Rassi, A., Jr., Rassi, A. & Little, W. C. Chagas’ heart disease. Clin Cardiol 23, 883–889 (2000). 10.1002/clc.4960231205

30 Soattin, L. et al. Endurance exercise remodels pulmonary vein sleeve myocytes and promotes a proarrhythmic atrial substrate. Eur Heart J (2026). 10.1093/eurheartj/ehag358

31 Mesirca, P. et al. Intrinsic Electrical Remodeling Underlies Atrioventricular Block in Athletes. Circ Res 129, e1–e20 (2021). 10.1161/CIRCRESAHA.119.316386

32 Tikhomirov, R. et al. Cardiac GR Mediates the Diurnal Rhythm in Ventricular Arrhythmia Susceptibility. Circ Res 134, 1306–1326 (2024). 10.1161/CIRCRESAHA.123.323464

33 Mangoni, M. E. & Nargeot, J. Properties of the hyperpolarization-activated current (I(f)) in isolated mouse sino-atrial cells. Cardiovasc Res 52, 51–64 (2001). 10.1016/s0008-6363(01)00370-4

34 Chandler, N. J. et al. Molecular architecture of the human sinus node: insights into the function of the cardiac pacemaker. Circulation 119, 1562–1575 (2009). 10.1161/CIRCULATIONAHA.108.804369

35 Monfredi, O., Dobrzynski, H., Mondal, T., Boyett, M. R. & Morris, G. M. The anatomy and physiology of the sinoatrial node--a contemporary review. Pacing Clin Electrophysiol 33, 1392–1406 (2010). 10.1111/j.1540-8159.2010.02838.x

36 Mages, S. et al. TACCO unifies annotation transfer and decomposition of cell identities for single-cell and spatial omics. Nat Biotechnol 41, 1465–1473 (2023). 10.1038/s41587-023-01657-3

37 Heras-Bautista, C. O. et al. Cardiomyocytes facing fibrotic conditions re-express extracellular matrix transcripts. Acta Biomater 89, 180–192 (2019). 10.1016/j.actbio.2019.03.017

38 Kubin, T. et al. Oncostatin M is a major mediator of cardiomyocyte dedifferentiation and remodeling. Cell Stem Cell 9, 420–432 (2011). 10.1016/j.stem.2011.08.013

39 Yanai, S. & Endo, S. Functional Aging in Male C57BL/6J Mice Across the Life-Span: A Systematic Behavioral Analysis of Motor, Emotional, and Memory Function to Define an Aging Phenotype. Front Aging Neurosci 13, 697621 (2021). 10.3389/fnagi.2021.697621

40 Flurkey, K., Currer, J. & Harrison, D. in The Mouse in Biomedical Research (eds JG Fox et al.) Ch. 20, 637–673 (Academic Press, 2007).

41 Moghtadaei, M. et al. The impacts of age and frailty on heart rate and sinoatrial node function. J Physiol 594, 7105–7126 (2016). 10.1113/JP272979

42 Ramilowski, J. A. et al. A draft network of ligand-receptor-mediated multicellular signalling in human. Nat Commun 6, 7866 (2015). 10.1038/ncomms8866

43 Sharpe, E. J., Larson, E. D. & Proenza, C. Cyclic AMP reverses the effects of aging on pacemaker activity and If in sinoatrial node myocytes. J Gen Physiol 149, 237–247 (2017). 10.1085/jgp.201611674

44 Vivas, O. et al. Aging Disrupts L-type Ca(2+) Channel Organization and Function in Pacemaker Cells. Circ Res 139, e327894 (2026). 10.1161/CIRCRESAHA.125.327894

45 Drouin, E. Electrophysiologic properties of the adult human sinus node. J Cardiovasc Electrophysiol 8, 254–258 (1997). 10.1111/j.1540-8167.1997.tb00788.x

46 Tsutsui, K. et al. A coupled-clock system drives the automaticity of human sinoatrial nodal pacemaker cells. Sci Signal 11 (2018). 10.1126/scisignal.aap7608

47 Sugita, J. et al. Cardiac macrophages prevent sudden death during heart stress. Nat Commun 12, 1910 (2021). 10.1038/s41467-021-22178-0

48 Al-Othman, S., Wu, Y., Fontanaud, P., Puttur, F. & Conesa, D. Resident cardiac macrophages are not required for normal atrioventricular node conduction. bioRxiv 2026 (2026). 10.64898/2026.02.11.704506

49 Lu, Z. et al. Regulation of HCN2 Current by PI3K/Akt Signaling. Front Physiol 11, 587040 (2020). 10.3389/fphys.2020.587040

50 Catalucci, D. et al. Akt regulates L-type Ca2+ channel activity by modulating Cavalpha1 protein stability. J Cell Biol 184, 923–933 (2009). 10.1083/jcb.200805063

51 Hiraoka, E. et al. TNF-alpha induces protein synthesis through PI3-kinase-Akt/PKB pathway in cardiac myocytes. Am J Physiol Heart Circ Physiol 280, H1861–1868 (2001). 10.1152/ajpheart.2001.280.4.H1861

52 Mesquita, T., Miguel-Dos-Santos, R. & Cingolani, E. Aging and sinus node dysfunction: mechanisms and future directions. Clin Sci (Lond) 139, 577–593 (2025). 10.1042/CS20231025

53 Widjaja, A. A. et al. Inhibition of IL-11 signalling extends mammalian healthspan and lifespan. Nature 632, 157–165 (2024). 10.1038/s41586-024-07701-9

54 Chung, E. S. et al. Randomized, double-blind, placebo-controlled, pilot trial of infliximab, a chimeric monoclonal antibody to tumor necrosis factor-alpha, in patients with moderate-to-severe heart failure: results of the anti-TNF Therapy Against Congestive Heart Failure (ATTACH) trial. Circulation 107, 3133–3140 (2003). 10.1161/01.CIR.0000077913.60364.D2

55 Mann, D. L. et al. Targeted anticytokine therapy in patients with chronic heart failure: results of the Randomized Etanercept Worldwide Evaluation (RENEWAL). Circulation 109, 1594–1602 (2004). 10.1161/01.CIR.0000124490.27666.B2

56 Ridker, P. M. et al. Antiinflammatory Therapy with Canakinumab for Atherosclerotic Disease. N Engl J Med 377, 1119–1131 (2017). 10.1056/NEJMoa1707914

57 Vasilets, L. M. T., A. V., Khlynova, O. V., Vustina, V. V. & Ratanova, E.A. Biomarkers role in the development of sinus node dysfunction. Russian Journal of Cardiology, 43–84 (2012).

