## Supplemental Material for "TNF inhibition reveals a reversible component of age-associated sinoatrial node dysfunction"

### Supplemental Methods

Human cardiac tissue was obtained through the Imperial College Human Tissue Programme. Donor non-failing hearts were obtained under IRAS project ID 189069 (REC reference 16/LO/1568). Tissue collection and use were conducted in accordance with the Human Tissue Act 2004 and relevant institutional approvals, with written informed consent obtained from donors or, where applicable, their next of kin. All animal procedures were conducted in accordance with the UK Animals (Scientific Procedures) Act 1986, under UK Home Office project licence PP3219770, and were approved by the Animal Welfare and Ethical Review Bodies of Imperial College London and the University of Manchester.

#### Human donor hearts

Human sinoatrial node (SAN) tissue samples were collected from donor hearts (provided by NHS Blood and Transplant, UK). Donor and clinical information are provided in **Supplemental Table 1**. Hearts were transported on ice in saline, cardioplegia, or University of Wisconsin (UW) solution, and cold ischemic time defined as the interval from organ retrieval to start of experiments, was less than 7 hours. The SAN was dissected by opening the superior vena cava and isolating tissue along the crista terminalis near the junction with the right atrium, maintaining transmural orientation. For Xenium spatial transcriptomic analysis, dissected tissue was embedded in optimal cutting temperature (OCT) compound, snap-frozen in liquid nitrogen, and stored at -80 °C for preservation.

#### Human SAN slice preparation and culture

Human SAN tissue blocks (approximately 1.5 $cm^3$) were mounted onto a vibratome specimen holder using Histoacryl adhesive (B.Braun) on a 4% agarose base in an ice-cold, oxygenated Tyrode's solution supplemented with 30 mM 2,3-butanedione monoxime. SAN slices (500 µm) were cut parallel to the endocardial surface using a vibratome (7000smz-2 Campden Instruments) at a speed of 0.03 mm/s. Slices were cultured in Medium 199 with Earle's salts (InVitroSys) supplemented with 1% ITS, 2% penicillin–streptomycin, and 20 mg/L ascorbic acid at 37 °C in 95% $O_2$ / 5% $CO_2$ for 48 h. Control slices received vehicle alone, whereas treatment slices were exposed to recombinant human TNF (100 ng/mL; R&Dsystems).[1,2]

#### Mouse studies

Mice were housed five per cage in a temperature-controlled room (22°C) on a 12 h:12 h light:dark cycle with free access to food and water. HCN4/TNF$^{glo}$ transgenic experiments used male FVB/N background mice (initial body weight 20–25 g, 8–12 weeks of age); ageing studies used male C57BL/6NRj mice (Janvier; initial body weight 25-30 g), with young controls at 2 months and aged animals at 22–24 months. Inclusion and exclusion criteria were defined prospectively and no animals were excluded from analysis. Following acclimatisation, animals were allocated to vehicle or treatment groups by simple randomisation using a random-number generator. Investigators were blinded to group allocation during echocardiographic analysis, telemetry analysis, Langendorff perfusion, multielectrode array mapping, sharp microelectrode recordings, whole-cell patch-clamp electrophysiology, RNA sequencing, proteomic profiling, flow cytometry, histology, immunolabelling, qPCR and western blotting.

#### HCN4/TNF$^{glo}$ mouse model

FVB/N-Tg(CAG-EGFP,-TNF)1Kul/J (TNF$^{glo}$) mice (The Jackson Laboratory, JAX strain #028698) were crossed with Hcn4$^{tm1(Cre/ERT2)Anlu}$ (JAX Strain #:024283, HCN4-CreERT2) mice to generate double-positive HCN4/TNF$^{glo}$ mice. The TNF$^{glo}$ transgene contains a CMV-IE enhancer/chicken beta-actin/rabbit beta-globin hybrid promoter (CAG) followed by a loxP-flanked EGFP-polyadenylation cassette positioned upstream of mouse TNF cDNA. HCN4-CreERT2 mice contain a CreERT2 sequence inserted at the Hcn4 start codon, enabling tamoxifen-inducible Cre-mediated recombination in HCN4-expressing cells. HCN4/TNF$^{glo}$ mice were administered intraperitoneal tamoxifen (150 mg/kg body weight) (APExBIO, USA) in soybean oil (Sigma) on two consecutive days. Tamoxifen-induced Cre-mediated recombination excises the loxP-flanked EGFP-polyadenylation cassette, enabling CAG promoter-driven TNF expression in HCN4-expressing cells. Mice were studied six weeks following tamoxifen administration. Genotyping was performed by Transnetyx (Cordova, TN, USA) using real-time PCR with transgene-specific probes. As a control check, heart rate and SAN TNF levels were examined in tamoxifen-treated TNF$^{glo}$ and HCN4-CreERT2 mice and vehicle-treated HCN4/TNF$^{glo}$ mice, with no changes observed in any of these control groups. Controls included TNF$^{glo}$ mice and HCN4-CreERT2 mice injected with tamoxifen on two consecutive days, and HCN4/TNF$^{glo}$ mice injected with soybean oil on two consecutive days.

**Etanercept treatment in aged mice**

Aged mice received etanercept (2.5 mg $kg^{-1}$, i.p., twice weekly for 8 weeks; MedChemExpress) diluted in sterile 0.9% saline; vehicle-treated controls received an equivalent volume of sterile saline. Aged mice entered the study at 22 months of age and were treated for up to 8 weeks. In accordance with the approved animal protocol, animals were not maintained beyond 24 months of age; 24 months therefore represented the prespecified terminal study endpoint. Survival analyses consequently assessed mortality during the 8-week treatment period rather than lifespan beyond 24 months. Terminal physiological and molecular analyses were performed in separate age-matched cohorts of young, aged vehicle-treated and aged etanercept-treated mice (N=8/group), independent of the cohort used for longitudinal frailty and survival analysis.Young (2-month-old) mice and vehicle-treated aged-matched 22-month-old mice served as controls. Frailty was assessed weekly using a validated mouse clinical frailty index.[3-5] The dosing regimen was selected based on prior murine studies demonstrating effective TNF suppression at 1-2.5 mg/kg in chronic inflammatory and ageing-related models, while avoiding the high-dose exposure associated with anti-drug antibody formation and reduced efficacy.

**Justification of dose and regimen**: Previous studies have demonstrated effective suppression of TNF-driven inflammation using etanercept doses between ~0.5-4 mg/kg administered one to three times weekly. In aged mice, weekly administration of 1 mg/kg for 12 months prevented sarcopenia and systemic inflammatory changes,[6] while 1 mg/kg twice weekly reversed age-related TNF-dependent alterations in colonic function.[7] Additional studies demonstrated efficacy using 1-2.5 mg/kg regimens in inflammatory and cardiovascular disease models, including exercise-induced atrial remodelling,[8] autoimmune uveitis,[9] and chronic infection.[10] The selected 2.5 mg/kg dose therefore lies within the established efficacious range reported in mice while remaining substantially below high-dose regimens (>20 mg/kg) associated with anti-drug antibody formation and loss of efficacy.[11,12]

**Murine *in vivo* assessment**

**Transthoracic echocardiography**

Echocardiographic assessments were performed in accordance with modified standard protocols under light isoflurane anaesthesia. Anaesthesia was induced with 3% isoflurane and maintained at 1-1.5% throughout image acquisition. Animals were placed in the supine position on a temperature-controlled platform, with continuous electrocardiogram (ECG) and respiratory rate monitoring.

Imaging was conducted using a Vevo 3100 system (VisualSonics) equipped with a 40 MHz linear-array transducer (MX-550D). All data were analysed offline using VevoLab software (version 5.5.0) with integrated cardiac and vascular analysis packages.

**Left ventricular assessment:** Two-dimensional (B-mode) images were acquired in parasternal long-axis and short-axis views at the level of the papillary muscle. One-dimensional M-mode recordings were obtained at the mid-papillary level. Left ventricular (LV) structure was characterised by measuring LV posterior wall thickness (LVPW), LV internal diameter at end-diastole (LVID, d) and volume. Systolic function was assessed by measuring LV internal diameter at end-diastole and end-systole (LVID,d and LVID,s), enabling calculation of fractional shortening (LV-FS), ejection fraction (EF), stroke volume (SV), and cardiac output (CO).

**Apical views and diastolic function:** Apical four- and five-chamber views were acquired using two-dimensional B-mode imaging at the level of the atrioventricular valves. PW Doppler measurements were performed at the tips of the mitral valve to obtain peak early (E) and atrial (A) diastolic filling velocities.

Mitral valve inflow profiles were further analysed to determine isovolumetric relaxation time (IVRT) and mitral valve deceleration time (MVDT), measured from the onset or peak of the E wave to the return to baseline. Tissue Doppler imaging (TDI) was performed at the mitral annulus to measure early (E′) and late (A′) diastolic myocardial velocities.

#### ECG recordings

Surface ECG recordings were obtained from anaesthetised mice as previously described.[13] Mice were anaesthetised with light isoflurane and placed in a supine position on a heated platform to maintain body temperature. Limb electrodes were positioned subcutaneously to obtain a standard three-lead ECG configuration. Recordings were acquired at baseline and during autonomic blockade following intraperitoneal administration of atropine (1 mg/kg) and propranolol (1 mg/kg). ECG signals were recorded continuously, and parameters were averaged over 100 consecutive cardiac cycles during stable sinus rhythm.

#### Telemetry

Control and HCN4/TNF$^{glo}$ mice (aged 8-12 weeks) were implanted with subcutaneous bio-telemeters (PhysioTel™ ETA-F10, Data Sciences International, DSI), as previously described.[14] Mice were anaesthetised by delivery of isoflurane in 100% oxygen via a nose cone. A 2% isoflurane concentration was used for induction and 1.5% for maintenance of anaesthesia. Animals were shaved and the surgical site was cleaned with iodine solution. Aseptic technique was maintained throughout the procedure. A dorsal paraspinal incision was performed, and blunt dissection was used to create a subcutaneous pocket for the telemetry device in the left flank above the hip. Two further small ventral incisions were made above the positions used for lead placement, and blunt dissection was performed to reach the pectoral muscles. Telemetry leads were tunnelled from the pocket to the chest wall and sutured to the pectoral muscles at right upper and left lower parasternal positions using 6-0 non-absorbable Prolene sutures. Bupivacaine (8 mg/kg made up to 1 mL in saline) was delivered into the subcutaneous pocket and the bio-telemeter was then placed. All incisions were closed with 6-0 absorbable Vicryl sutures. Ten minutes prior to the end of surgery buprenorphine (0.1 mg/kg) was delivered via intraperitoneal injection for pain relief. Another dose of buprenorphine (0.1 mg/kg) was given 6 hours post recovery, with further doses given twice a day for two days post-surgery. Following recovery from anaesthesia, animals were transferred to a warm cage. Animals were allowed to recover in their home cage for a minimum of 1 week prior to the commencement of ECG recordings.

24-hour ECG recordings were taken for mice at baseline and 6 weeks post tamoxifen injection. Data were analysed using Ponemah software (V6.71 DSI). Raw ECG traces were first visually inspected to confirm signal quality. Automated beat detection was then performed using the Ponemah ECG analysis module, with detection parameters optimised for murine ECG morphology. R waves were identified and used to generate beat-to-beat RR interval data, heart rate trends and rhythm annotations. Data were filtered for activity, heart rates outside of physiological range (200-850 BPM), and noise. For each 1-hour epoch, resting heart rate was defined as the median of the retained values. Arrhythmias were identified using Data Insights software (DSI). Candidate arrhythmic events were detected using predefined criteria based on manually defined parameters and subsequently reviewed manually against the raw ECG trace to exclude artefact and confirm arrhythmia classification. Sinus pauses were defined as the absence of a p wave or QRS complex for ≥300ms. Bradycardia was defined by 5 consecutive beats with a heart rate below 300 BPM. Group-level profiles were generated by calculating the mean ± SEM of the individual mice's hourly median heart rates at each Zeitgeber hour. A 24-hour sine curve was fitted to the hourly median heart rate profile using the equation $y = M + A\sin(2\pi t/24 + \phi)$, where $M$ represents the mesor, $A$ the amplitude, $t$ time in hours, and $\phi$ the phase shift. Formal cosinor analysis was performed using a fixed 24-hour period to estimate mesor, amplitude, and acrophase for each condition.

### Murine *ex vivo* assessments

#### Langendorff perfusion

Ten minutes prior to termination by cervical dislocation, heparin was administered via intraperitoneal injection (1000 IU/kg). The heart was excised, lungs and excess tissue removed, and the aorta cannulated with a 22 G Langendorff perfusion cannula. This was secured using 6-0 braided silk suture (Fine Science Tools). The heart then underwent retrograde perfusion with 37°C filtered Kreb's solution (in mmol $L^{-1}$: 118.5 NaCl, 4.75 KCl, 1.19 $MgSO_4 \cdot 7H_2O$, 1.19 $KH_2PO_4 \cdot H_2O$, 1.41 $CaCl_2$, 22 $NaHCO_3$, 11.0 glucose) which was bubbled with a 95% oxygen 5% CO2 gas mixture. Pressure was maintained at 80 mmHg using a physiological pressure transducer and pressure flow regulator (STH pump controller, AD Instruments), calibrated on LabChart 8 (AD Instruments) using a manual sphygmomanometer and connected to a peristaltic pump (Gilson). After 10 minutes to allow the heart to return to a stable rhythm, electrophysiological study was performed. The Mapping Lab EMS64-USB-1003 was used for pseudo-ECG recording and pacing protocols were delivered using the Mapping Lab VCS-3001 Stimulator. The pacing electrode was placed at the right atrial appendage and the threshold for capture was identified and a voltage 3x higher than this level was used for pacing, to ensure adequate capture throughout the experiment. ECG electrodes were placed at the apex and left atrial appendage. Once a satisfactory trace was achieved a pseudo-ECG was recorded. Following this the atria were paced to ascertain the corrected sinus node recovery time (cSNRT), Wenckebach cycle length (WBCL), AV node effective refractory period (AVERP) and atrial effective refractory period (AERP). Pacing protocols were designed based upon those described by Hennis et al.[15] An S1-S11 stimulation protocol was used to test atrial fibrillation (AF) inducibility. The pacing electrode was then moved to the left ventricular free wall, the threshold was re-assessed, and ventricular effective refractory period was tested.

#### Multi-electrode array mapping

Murine preparations encompassing the SAN, right atrial appendage, superior and inferior vena cava, interatrial septum, and adjacent right ventricular tissue were rapidly dissected following humane culling and excision of the heart. Preparations were then superfused with oxygenated Tyrode's solution (in mMol $L^{-1}$: 120.3 NaCl, 4.0 KCl, 1.3 $MgSO_4 \cdot 7H_2O$, 1.2 $NaH_2PO_4 \cdot H_2O$, 1.2 $CaCl_2$, 25.2 $NaHCO_3$, 11.0 glucose) at 37 °C, continuously bubbled with 95% $O_2$/5% $CO_2$. Following a stabilisation period, flexible high-density multi-electrode (64 electrodes with 0.5 mm electrode distance) arrays (MappingLab) were positioned over the SAN region to record unipolar electrograms during spontaneous rhythm and programmed electrical stimulation. A bipolar electrode was introduced at the right atrial appendage for pacing protocols, and spontaneous activity was also evoked by rapid pacing (10 Hz, 3 s).

Electrical signals were acquired at high temporal resolution and analysed offline using EMapScope (MappingLab). Data were bandpass- and notch-filtered to remove baseline drift and mains interference. Local activation times were defined from the steepest negative deflection of each electrogram, from which isochronal activation maps were constructed. Conduction velocity was calculated from spatial gradients of activation times between neighbouring electrodes, enabling quantification of impulse propagation and SAN conduction dynamics.

**Isolation of SAN cells**

SAN pacemaker myocytes were isolated as previously described.[16] Mice were deeply anesthetized by intraperitoneal injection of 0.3 ml of solution containing ketamine (0.1 mg/g, Imalgène, Merial, France) and xylazine (0.01 mg/g, Rompun 2%, Bayer, Germany), followed by a second injection of 0.3 ml of solution containing pentobarbital (150 µL Euthasol Vet, Virbac, France in 10 ml NaCl physiological solution). To avoid intracardiac blood clots, 0.5 ml of NaCl solution containing heparin (25 000 I.U.) was injected. Hearts were removed via thoracotomy when the tail sensitivity test became negative and immersed in a pre-warmed (36 °C) Tyrode solution containing (in mM): 140 NaCl, 5.4 KCl, 1 MgCl2, 1.8 CaCl2, 5.5 D-glucose, and 5 Hepes (adjusted to pH 7.4 with NaOH). SAN tissue was excised and transferred into a low-$Ca^{2+}$ solution, 36 C°, containing (in mM): 140 NaCl, 5.4 KCl, 0.5 MgCl2, 1.2 KH2PO4, 50 taurine, 5.5 D-glucose, 1 mg/mL BSA, and 5 Hepes (adjusted to pH 6.9 with NaOH) for 5 min. Then, enzymatic digestion was carried out for 20-25min at 36 °C in the low-Ca2+ solution containing Liberase-TH (0.15mg/mL; Roche, Mannheim, Germany) and elastase (0.5mg/mL; Worthington, Lakewood, New Jersey, USA). Digestion was stopped by washing the SAN in a "Kraftbrühe" (KB) medium containing (in mM): 100 L-glutamic acid potassium salt, 10 L-Aspartic acid potassium salt, 25 KCl, 80 KOH, 10 KH2PO4, 2 MgSO4, 5 Creatine, 0.5 EGTA, 20 D-glucose, 20 taurine, 1 mg/mL BSA, and 5 Hepes (adjusted to pH 7.2 with KOH). Single myocytes were dissociated from the SAN tissue by manual agitation using a flame-forged Pasteur's pipette in KB solution at 36°C. To recover spontaneous automaticity of the SAN cells, calcium was gradually reintroduced in the cell's storage solution to a final concentration of 1.8mM. SAN cells were then left to rest for 1 h before recordings.

**Patch clamp electrophysiology**

For patch-clamp recordings of SAN myocytes, cells were harvested in custom-made chambers with glass bottoms for cell attachment. Cells were then super-fused with standard Tyrode's solution warmed at 36 °C before recording.

The automaticity of SAN cells was recorded in basal extracellular Tyrode solution by the perforated patch-clamp technique (β-escin at 0.04mg/mL, Sigma Aldrich). $I_f$ was recorded using standard whole-cell variation of the patch-clamp technique in pre-warmed (36 °C) Tyrode's solution. Patch-clamp electrodes, pulled from borosilicate glass using a DMZ-Universal Electrode Puller (Zeitz Instruments, Germany), had a resistance of 3-4 MΩ when filled with an intracellular solution containing (mM): 80 L-Aspartic acid potassium salt, 50 KCl, 1 MgCl2, 5 Hepes, 2 CaCl2, 5 EGTA and 3 ATP-Na+ salt (adjusted to pH 7.2 with KOH). Seal resistances were in the range of 2–5 GΩ.

Voltage-gated L-type $Ca_v1.3$- and $Ca_v1.2$-mediated $Ca^{2+}$ currents ($I_{Cav1.3}$ and $I_{Cav1.2}$) were recorded using standard whole cell patch-clamp configuration in an extracellular solution containing (mM): 135 tetraethylammonium chloride (TEA-Cl), 10 4-aminopyridine (4-AP), 1 MgCl2, 0.03 tetrodotoxin (TTX, Latoxan), 5.5 D-glucose, 2 CaCl2, 10 Hepes, (adjusted to pH = 7.2 with HCl). To selectively block $I_{Cav1.2}$, 1µM of Calciseptine (Latoxan) was added to extracellular solution.

Recording electrodes had a resistance of 3 MΩ when filled with a solution containing (in mM): 125 CsOH, 20 TEA-Cl, 1.2 CaCl2, 5 Mg-ATP, 0.1 Li2-GTP, 5 EGTA, and 10 HEPES (pH adjusted to 7.2 with HCl). Seal resistances were in the range of 2–5 GΩ. Perfusion of pre-warmed (36 °C) experimental solutions was performed by using a multi- MPRE8 heating pen (Cell Micro Controls). Data acquisition was performed by employing an Axopatch 700B patch-clamp amplifier connected to a Digidata 1550B interface (Molecular Devices). Data were analysed using pCLAMP 10.7 (Molecular Devices).

**Sharp microelectrode recordings in human and mouse SAN**
Intracellular action potentials were recorded from intact murine SAN preparations and human SAN slices using sharp microelectrodes fabricated from filamented borosilicate glass capillaries (outer diameter 1.00 mm, inner diameter 0.58 mm; GC100F-10, Harvard Apparatus). Microelectrodes were pulled to a resistance of 20–30 MΩ when filled with 3 M KCl and mounted in a microelectrode holder (1-HL-U, Molecular Devices) containing an Ag/AgCl wire connected to the amplifier headstage. Preparations were superfused with Krebs–Ringer solution at 2 mL/min, continuously gassed with 95% $O_2$ / 5% $CO_2$, and maintained at 37 °C using a temperature-controlled heated inflow perfusion system with feedback regulation (Scientifica). Recordings were obtained using a MultiClamp 700B amplifier coupled to a Digidata 1550B digitiser (Molecular Devices). Signals were sampled at 50 kHz and low-pass filtered at 10 kHz. Data were acquired using Clampex 10 and analysed in LabChart 8 (ADInstruments). For murine experiments, impalements were obtained from the central SAN (SAN-C), peripheral SAN (SAN-P), and adjacent atrial myocardium. A minimum AP amplitude of 25 mV was required for inclusion. Atrial APs were identified by a maximum diastolic potential more negative than −69 mV. Among non-atrial recordings, SAN-C APs were identified using a peak-width criterion of ≥20 ms, with the remaining nodal recordings classified as SAN-P. Spontaneous firing rate, maximum upstroke velocity ($dV/dt_{max}$), diastolic depolarisation slope, maximum diastolic potential, AP amplitude and AP duration at 30%, 50% and 90% repolarisation ($APD_{30}$, $APD_{50}$, and $APD_{90}$) were quantified.

For human SAN slices, impalements were targeted to the subepicardial aspect of the nodal region where slice orientation permitted, reflecting the location of the human SAN beneath the epicardium near the junction of the superior vena cava and right atrium. Precise regional position within the node could not be mapped in the sliced preparation, and the waveform-based criteria used to subclassify murine SAN recordings were therefore not applied. Fresh recordings were benchmarked against published electrophysiological measurements from intact adult human SAN preparations and isolated human SAN myocytes (**Supplemental Table 2**). Spontaneous cycle length, maximum diastolic membrane potential, maximum upstroke velocity, AP amplitude and $APD_{90}$ were quantified. Isoprenaline (0.1 µM) was applied to a subset of freshly prepared slices to assess preserved β-adrenergic responsiveness. Additional recordings obtained from one donor after 12, 24 and 48 h of organotypic culture in the presence of TNF demonstrated persistence of spontaneous electrical activity during culture (**Supplemental Figure 11A-D**). These recordings were not used to infer TNF-specific electrophysiological remodelling because equivalent time-matched control recordings were not available at each interval. Molecular analyses after 48 h were performed using within-donor comparisons between TNF-treated slices and time-matched controls.

**Mouse SAN culture and NF-κB inhibition**
Eight-week-old male C57BL/6 mice were used for ex vivo SAN culture experiments (N=4 per group). Isolated mouse SAN preparations were dissected as described above and individually transferred into MatTek dishes containing 5 mL of pre-warmed Medium 199 and maintained at 37 °C with 5% $CO_2$ for 30 min to recover. Control preparations were maintained in Medium 199 under the same culture conditions for 48 h without additional treatment. To investigate the contribution of NF-κB signalling to TNF-mediated transcriptional remodelling of pacemaker ion channels, NF-

κB signalling was inhibited using BMS-345541 (5 µM; Sigma-Aldrich, Cat# B9935), an allosteric inhibitor of IκB kinase (IKK) α and β. BMS-345541 was dissolved in dimethyl sulfoxide (DMSO). Based on previous studies using BMS-345541 pre-treatment prior to TNF stimulation,[17-19] preparations were pre-incubated overnight for 12 h with either BMS-345541 (5 µM; 0.05% DMSO) or vehicle (0.05% DMSO). Following pre-incubation, TNF-treated preparations were exposed to recombinant mouse TNF (20 ng/mL; PeproTech, Cat# 315-01A), reconstituted in sterile PBS containing 0.1% BSA. At 12 h following TNF addition, 50% of the culture medium was replaced with fresh pre-warmed Medium 199 containing the corresponding BMS-345541 or vehicle treatment and TNF (20 ng/mL). Preparations were harvested 48 h after TNF addition, briefly washed in ice-cold PBS, snap-frozen in liquid nitrogen and stored at −80 °C.

**Conserved NF-κB binding motif analysis.** For *HCN4, HCN1, CACNA1D* and *TBX3*, the 10 kb immediately upstream of the transcription start site together with the full transcript was retrieved for the human (GRCh38) and mouse (GRCm39) reference transcripts ENST00000261917.4 and ENSMUST00000034889.11 (HCN4), ENST00000303230.6 and ENSMUST00000006991.10 (HCN1), ENST00000350061.11 and ENSMUST00000238504.3 (CACNA1D), and ENST00000349155.7 and ENSMUST00000568276.1 (TBX3). Sequences were scanned for NF-κB binding motifs using TRANSFAC positional weight matrices using rVista.[20] Motifs conserved between the aligned human and mouse loci were retained and overlapping predictions within a locus were merged into distinct conserved regions, which were then classified by position as upstream, proximal promoter, intronic or exonic relative to the Ensembl release 116 gene models.

### Molecular and Cellular Profiling

#### Xenium *in situ* spatial transcriptomics analysis of aged versus young human SAN

Given the cellular and spatial heterogeneity of the human SAN and the limited availability of donor tissue, we used Xenium spatial transcriptomics to identify cell-type-specific transcriptional changes and alterations in the local microenvironment associated with ageing. Cell-level analyses provided high-resolution discovery, while key findings were subsequently evaluated using donor-level pseudobulk analyses and, where possible, independent molecular validation.

**Tissue preparation, histology and data acquisition:** Freshly dissected human SAN (hSAN) preparations were snap-frozen, placed in transfer cassettes and immediately frozen in OCT for cryosectioning. On the day of sectioning, cryoblocks were equilibrated to the cryostat chamber (Leica CM1950) temperature. All surfaces and instruments were decontaminated with RNase*Zap* (AM9780, Thermofisher Scientific). MX35 Ultra blades were used to trim blocks at -23 °C. Cryoblocks were sectioned at 10µm. SAN structures were identified with a Masson's Trichrome staining kit (Sigma Aldrich, HT15-1KT) and immunolabelling for Cx43 and $Ca_V1.3$. For Xenium analysis, freshly cut sections were transferred onto proprietary 10x Genomics Xenium sample slides which were stored at -80 °C. Adjacent sections were retained for quality assessment and validation of Xenium findings. On the day of experiment, sections were permeabilised and then subjected to probe hybridisation, ligation and amplification using Xenium Human immuno-oncology panel (380 genes) supplemented with a custom add-on panel of 100 genes, following manufacturer's guidelines (**Supplemental Table 3**). The 100 custom genes were pre-selected and adjusted to fit a predetermined optical budget. Following probe hybridisation, ligation and amplification, slides were imaged and probe decoding and cell segmentation were performed on the Xenium Analyser. Cells were segmented using a proprietary Xenium deep learning model incorporating a boundary stain (**Supplemental Figure 2C).**[21] Transcriptional data were obtained from 272715 cells across 7 donors (**Supplemental Table 4**) and were further visualised with Xenium Explorer software v3.1.0.

**Data processing:** Xenium Spatial transcriptomic datasets were processed to examine tissue molecular architecture and neighbourhood interactions in aged vs young hSAN. Data preprocessing and subsequent analyses were performed using Squidpy (v. 1.6.1),[22] a comprehensive spatial omics data analysis toolkit,[23] CellCharter,[24] and R 4.4.1. packages: Seurat[25] and ggplot2.[26] Data visualisation was performed in Xenium Explorer or R. The dataset provided spatial coordinates and transcriptional profiles for individual cells. Cell clusters were identified across aged and young human SAN datasets jointly and annotated based on dominant cell-type composition to ensure biological coherence across conditions. Initial preprocessing removed low-quality cells and transcripts to minimise technical noise and ensure robust downstream analyses. Cellular quality control metrics were computed, including total transcript counts, unique gene counts per cell, and cellular morphology properties (**Supplemental Figure 2A**). Visualisation of these distributions informed the thresholds for subsequent filtering. Cells with fewer than ten total counts and genes detected in fewer than five cells were excluded. These thresholds were applied to retain only high-quality observations with sufficient transcriptomic and spatial information for downstream analyses.

**Normalisation, dimensionality reduction and downstream analysis:** Total transcript counts per cells were normalised to a uniform scale using Squidpy's integrated normalisation functions. Data distribution was assessed and visualised using Q-Q plot (**Supplemental Figure 2B, left**). Logarithmic transformation was applied to reduce skewness and compress the dynamic range of highly expressed features in overdispersed[27] spatial transcriptomic data. Dimensionality reduction was performed by principal component analysis (PCA), retaining the top 20 components accounting for 82.577% variance as assessed by Scree plot (**Supplemental Figure 2B, right**). Data were visualised using UMAP embeddings, with cells coloured by cluster identity, total transcript counts, and key functional attributes. Cell type annotation was performed using TACCO,[27] with a matched single-cell RNA-sequencing dataset serving as the reference atlas.[28] TACCO performs probabilistic annotation by modelling each spatial transcriptomic profile as a mixture of reference cell type expression signatures and inferring cell type proportions through optimal transport–based alignment between datasets. Prior to annotation, genes shared between the spatial and reference datasets were retained, and datasets were normalised according to the Scanpy preprocessing workflow. The inferred cell type abundance matrix was subsequently used for downstream analysis and visualisation. Annotated cell clusters were further used for cell fractions (%) quantification per group. Because some preparations contained relatively few cells of specific populations, cell-type composition was assessed using the underlying pooled cell counts, preserving the information contained in the number of cells contributing to each proportion. Differences between young and aged SAN datasets were tested using two-sided Fisher's exact tests. Differential expression analysis between aged and young human SAN was performed on transcript counts within each cluster using Mann-Whitney U test with Benjamini-Hochberg correction in line with previously described methods for rare human samples.[29,30] Differential gene expression was used to characterise identified cell populations by known canonical marker gene expression (**Supplemental Figure 2D-F**). To account for donor-level biological replication, raw Xenium transcript counts were additionally aggregated by donor and annotated cell population, followed by differential expression analysis using DESeq2 with Benjamini–Hochberg correction. Concordance between cell-level and donor-level findings was assessed by comparing $\log_2$ fold-change estimates.

**Cellular niche identification, annotation and spatial plot:** Cellular niche prediction was performed using CellCharter by constructing k-nearest neighbour (kNN) graphs across a range of k = 5-20. Cellcharter aggregates information from each cell's own transcriptional identify as well as from its immediate neighbours and second-order neighbours. Cluster stability was assessed across increasing aggregation depths (**Supplemental Figure 3A**). Once an optimal cluster number was defined, cells were assigned to niches, which were annotated against previously defined cell types within the same dataset (**Supplemental Figure 3B**). Spatial niche maps were visualised in CellCharter and Xenium Explorer (**Supplemental Figure 3C**).

**Neighbourhood enrichment analysis:** A kNN graph (k = 6, Squidpy default) was constructed to model ageing-induced changes in cellular neighbourhoods. Neighbourhood enrichment analyses were performed to interrogate the spatial organisation of cell types and their enrichment patterns across conditions using Squidpy's gr.nhood_enrichment module, which calculates a z-score quantifying the enrichment of each cluster's spatial proximity to other clusters. Observed neighbour frequencies were compared with those derived from random permutations (n = 1000), controlling for biases in cluster abundance and spatial distribution. Pairwise cluster interactions were visualised as heatmaps (**Supplemental Figure 3D**), and neighbourhood enrichment scores for pacemaker cells were plotted separately as a bar chart using Squidpy's plotting tools (**Figure 2D**).

**Spatial organisation of pacemaker cardiomyocytes:** Pacemaker cardiomyocyte density was calculated as the number of annotated pacemaker cardiomyocytes per mm² of segmented tissue area. Nearest-neighbour distances and mean distances to the three and five nearest pacemaker cardiomyocytes were computed from cell centroids using the FNN R package.[31] Donor-level values were compared between age groups using Mann–Whitney U tests with Benjamini–Hochberg correction across the four measures.

**Spatial proximity analysis:** Spatial proximity between pacemaker cardiomyocytes (pCMs) and AREG+ macrophages was assessed using cell-centroid coordinates derived from Xenium spatial transcriptomics data.[30,32] Pacemaker cardiomyocytes and AREG+ macrophages were identified according to their annotated cell identities. For each pCM, the Euclidean distance to the nearest AREG+ macrophage was calculated using a nearest-neighbour search implemented with the FNN R package. To assess the robustness of proximity-dependent transcriptional differences to the definition of neighbourhood size, differential expression was examined across distance thresholds of 10, 30, 60, 90, 120 and 150 μm, and overlap between the resulting gene sets was visualised using an UpSet plot (**Supplemental Figure 3F**). A 60 μm threshold was used for the primary proximity analysis as an intermediate distance within the range over which overlapping transcriptional differences were observed. Pacemaker cardiomyocytes located within 60 μm of an AREG+ macrophage were classified as AREG+ macrophage-proximal, whereas those located >60 μm away were classified as AREG+ macrophage-distal. The resulting spatial classification was incorporated into the Seurat object and used to subset pCMs for downstream analysis. Differential gene expression between AREG+ macrophage-proximal and AREG+ macrophage-distal pCMs was assessed at the single-cell level using a two-sided Wilcoxon rank-sum test. P values were corrected for multiple testing using the Benjamini–Hochberg false discovery rate procedure. To validate these findings at the donor level, raw transcript counts from $AREG^+$ macrophage-proximal and macrophage-distal pacemaker cardiomyocytes were aggregated separately for each donor and analysed using DESeq2 with Benjamini–Hochberg correction.

**RNA sequencing**

RNA sequencing and bioinformatic analyses were performed as previously described.[13] Briefly, total RNA was isolated from SAN tissue, and libraries were prepared for bulk RNA sequencing. Sequencing was performed on an Illumina NovaSeq 6000 platform, and reads were aligned to the mouse reference genome. Gene-level counts were generated using standard bioinformatic pipelines, and differential gene expression analysis was performed using DESeq2 (v1.38.3) with Benjamini–Hochberg false discovery rate (FDR) correction. Differentially expressed genes were subsequently subjected to pathway enrichment and network analyses. To evaluate pro-inflammatory signalling within the mouse SAN, significantly upregulated cytokines and their receptors ($log_2FC > 0$, FDR < 0.05) were identified from the KEGG cytokine-cytokine receptor interaction pathway[33] among differentially expressed transcripts.[34]

Signalling pathway activity was inferred from normalised bulk RNA-sequencing counts using PROGENy[35], following $log_2$(count + 1) transformation and application of the mouse-specific model with the top 500 pathway-responsive genes per pathway. Pathway activity scores were compared between young and aged SAN using Wilcoxon rank-sum tests with Benjamini–Hochberg correction. Differential activity was expressed as the difference between aged and young scores.

**Proteomics**

Micro-dissected, snap-frozen SAN biopsies were sent to Biogenity (Aalborg, Denmark) for processing and proteomic analysis. Biogenity contributed to downstream proteomic data analysis. The Technical University of Denmark performed the proteomics sample analysis.

For quantitative proteomic analysis, samples were prepared for liquid chromatography–tandem mass spectrometry (LC-MS/MS) using previously published methodology.[36,37] Tissue was homogenised in detergent-based lysis buffer containing protease and phosphatase inhibitors, followed by acetone precipitation. Protein pellets were resolubilised in guanidinium hydrochloride buffer, reduced with tris(2-carboxyethyl)phosphine and alkylated with chloroacetamide. Protein concentrations were determined using a bicinchoninic acid assay, and 20 µg protein per sample was taken forward for analysis. Proteins were sequentially digested with LysC and trypsin, and the resulting peptides were desalted by solid-phase extraction. Peptides were subsequently labelled using Tandem Mass Tags (TMTpro 18-plex; Thermo Fisher Scientific) to enable multiplexed quantitative analysis.

Following labelling, samples were pooled and subjected to offline high-pH reversed-phase fractionation. Peptides were separated over a 120-minute gradient into 60 fractions, which were combined into 30 final fractions. LC-MS/MS analysis was performed using an Orbitrap Eclipse Tribrid mass spectrometer equipped with a FAIMS Pro interface and coupled to an Evosep One liquid chromatography system (Thermo Fisher Scientific). A 30-samples-per-day acquisition method was employed, with peptides separated over a 44-minute gradient. Data were acquired using an SPS-MS3 TMT workflow with FAIMS compensation voltages of −50 and −70 V. Instrument performance was monitored using a complex cell-lysate quality-control standard.

Raw mass spectrometry data were processed using Proteome Discoverer v2.4 (Thermo Fisher Scientific). Spectra were searched against the *Mus musculus* UniProt reference proteome. Oxidation of methionine and protein N-terminal acetylation were specified as dynamic modifications, whereas cysteine carbamidomethylation and TMT labelling of peptide N-termini and lysine residues were specified as static modifications. Peptide and protein identifications were filtered at a 1% FDR. TMT reporter-ion abundances were normalised to the total peptide abundance within each channel to account for differences in sample loading. Pre-analysis quality control included principal component analysis and hierarchical clustering to assess sample relationships and data integrity. To minimise missing-value-driven variance while retaining condition-specific proteins, proteins were retained if quantified in at least 60% of all samples or in at least 75% of samples within at least one experimental group. Differential protein abundance was assessed using the limma framework, employing parametric linear modelling with moderated analysis of variance to stabilise variance estimated in the small-sample design.[38] P values were adjusted for multiple testing using Benjamini-Hochberg FDR procedure. Proteins not returned by the primary limma analysis but meeting the abundance criterion were assessed separately using exact two-sided Wilcoxon rank-sum tests. Functional annotation and pathway enrichment analyses were performed using Reactome and Gene Ontology (GO) databases to identify biological processes and molecular pathways associated with the observed proteomic changes.[39-41]

**Flow Cytometry**

**Tissue processing and leucocyte stimulation**: Following microdissection, SAN tissue was finely minced and incubated for 1 hour at 37°C in a sample digestion buffer containing: 1 x phosphate buffered saline (PBS) + $Ca^{2+/}Mg^{+}$, 1% Bovine Serum Albumin (BSA), 50 mM HEPES, Collagenase I (450 U/mL), Collagenase XI (125 U/mL), Hyaluronidase (60 U/mL), DNAse I (60 U/mL). Following digestion, the cell suspension was passed through a 50 µM cell strainer and washed with RPMI + 2 mM EDTA. Isolated cells were either used directly for flow cytometric staining or subjected to ex ex *vivo* leucocyte restimulation to assessTNF production. For restimulation, cells were incubated for 4 hours at 37°C/5% $CO_2$ with phorbol 12-myristate 13-acetate (PMA; 50 ng/mL), ionomycin (500 ng/mL) and Brefeldin-A (Golgi inhibitor; 1x) in RPMI supplemented with 10% foetal calf serum (FCS), 2 mM L-glutamine, penicillin (100 U/mL) and streptomycin (100 µg/mL).

**Flow cytometry staining, sample acquisition and analysis**: Isolated cells were stained with LIVE/DEAD™ Fixable Blue Dead Cell Stain Kit (Invitrogen), followed by Fc receptor blockade (TruStain FcX^TM^, Biolegend), followed by surface antigen staining using antibodies detailed in **Supplementary Table 5.** For intracellular antigen staining, cells were first fixed and permeabilised using the Foxp3 / Transcription Factor Staining Buffer Set (eBioscience). Spectral flow cytometric parameter unmixing and data acquisition was conducted using a Cytek Aurora flow cytometer and SpectroFlo® software (Cytek Biosciences). Data were analysed using FlowJo software (Treestar). Absolute cell counts were determined using Precision Count Beads™ (Biolegend).

**Cryosectioning**

Tissues were embedded in OCT compound and snap-frozen in liquid nitrogen and stored at -80 °C. For cryosectioning, tissue blocks were mounted in a cryostat (Leica CM1950) maintained at -20 to -25 °C, and blocks, charged slides (Epredia™ Permafrost™), anti-roll plate, and dissection tools were equilibrated within the chamber for at least 30 min before sectioning. SAN tissue was sectioned perpendicular to the crista terminalis to preserve anatomical orientation within the sectioning plane. Blocks were trimmed at 20 µm until tissue was reached, using the SAN artery and surrounding right atrial tissue as anatomical landmarks to identify the SAN region, before serial 10 µm sections were collected. Sections were mounted onto slides and stored at -80 °C.

**Masson's trichrome staining**

Cryosectioned tissue was thawed at room temperature, and residual OCT was removed by PBS washes. Sections were fixed overnight in Bouin's solution, washed in 70% ethanol, and incubated in celestine blue, and sequentially stained with Weigert's haematoxylin, Biebrich scarlet-acid fuchsin, phosphomolybdic/phosphotungstic acid, and aniline blue using the Masson's trichrome staining kit (Sigma-Aldrich, HT15-1KT). Sections were then treated with 1% acetic acid, dehydrated through graded ethanols, cleared in Histo-Clear II, and mounted with DPX under glass coverslips. Brightfield images were acquired using a Leica microscope. The SAN region was identified by the presence of the SAN artery and selected for analysis. Extracellular matrix fractional area was quantified in Fiji/ImageJ using automated colour thresholding.

**Immunolabelling and image analysis**

Frozen human and mouse SAN cryosections were fixed in 4% paraformaldehyde for 10 min (human) or 30 min (mouse) at room temperature, washed in PBS containing 0.05% Tween-20 (PBS-T), and permeabilised with 0.3% Triton X-100 in PBS for 10 min. Sections were blocked for 1 h in 2% bovine serum albumin (BSA) in PBS-T before incubation overnight at 4 °C with primary antibodies diluted in blocking buffer (**Supplemental Table 6**). Following PBS-T washes, sections were incubated with Alexa Fluor-conjugated secondary antibodies (1:250, Thermo Fisher Scientific) for 3 h at room temperature. Cell membranes were labelled with Alexa Fluor 488-

conjugated wheat germ agglutinin (WGA), and sections were mounted with Vectashield Antifade mounting medium containing DAPI (Vector Laboratories) for nuclear counterstaining. Images were acquired using a Leica STELLARIS 8 confocal microscope using identical acquisition settings for all samples within each experiment.

Fluorescence intensity, positive tissue area, cell counts, and co-localisation were quantified using Fiji/ImageJ (version 1.54p). Colocalisation was assessed using the JACoP (Just Another Colocalisation Plugin) plugin. Manders' overlap coefficients (M1 and M2) were calculated to quantify the degree of fluorescence colocalisation between markers. For TNF and immune-cell marker analyses, Manders' M2 coefficient was used to quantify the proportion of TNF fluorescence colocalising with CD68 or CD8. Threshold-based segmentation was used with thresholds standardised within each experiment and normalised to tissue area. Technical replicates were averaged to generate a single value for each biological replicate, and representative images were selected to reflect typical staining patterns within each experimental group.

**Quantitative real-time PCR**

Quantitative real-time PCR (qPCR) was performed according to previously described methods.[42] Total RNA was extracted from frozen tissue using the RNeasy Mini Kit (Qiagen, #74104), and RNA concentration and purity were assessed by NanoDrop spectrophotometry (Thermo Fisher Scientific). cDNA was synthesised using the High-Capacity RNA-to-cDNA™ Kit (Applied Biosystems, #4387406). qPCR was performed using QuantiTect Primer Assays (Qiagen) and Power SYBR™ Green PCR Master Mix (Applied Biosystems) on a QuantStudio 7 Flex Real-Time PCR System (Applied Biosystems). Relative gene expression was normalised to GAPDH (human) and IPO8 (mouse) calculated using the $2^{-\Delta\Delta Ct}$ method. Primer assay details are provided in **Supplemental Table 7**.

**Western blotting**

Protein was extracted from tissue using the RNeasy flow-through protein precipitation protocol (Qiagen) following RNA extraction. Protein pellets were resuspended in 2x Laemmli sample buffer and heated at 37°C for 10 min before separation on 4-20% Mini-PROTEAN® TGX™ precast gels (Bio-Rad) alongside Precision Plus Protein™ standards (Bio-Rad). Proteins were transferred to 0.2 µm PVDF membranes using the Trans-Blot® Turbo Transfer System (Bio-Rad). Total protein transfer was visualised using stain-free imaging on a ChemiDoc MP Imaging System (Bio-Rad) prior to membrane blocking in 5% milk in TBST for 1 h at room temperature. Membranes were incubated overnight at 4°C with primary antibodies (**Supplemental Table 8**), followed by HRP-conjugated secondary antibodies for 1 h at room temperature. Proteins were detected using Clarity™ western ECL substrate (Bio-Rad) and imaged using a ChemiDoc MP Imaging System (Bio-Rad). Band intensities were quantified using Image Lab software (Bio-Rad) and normalised to stain-free total protein loading. All western blots were performed in duplicate.

**Statistical Analysis**

Statistical analyses were performed using GraphPad Prism v10.0 (GraphPad Software, Inc.) and R v4.4.0. Data are presented as mean ± standard deviation (SD) unless otherwise stated. Exact P values and 95% confidence intervals (CI) are reported for statistical comparisons. Residual diagnostics were examined, and normality was assessed using the Shapiro-Wilk test. For continuous variables, Welch's two-sided t-tests were used for normally distributed data with unequal variances, whereas Mann-Whitney U tests were used for non-normally distributed data. Paired comparisons were analysed using two-sided paired t-tests. Comparisons involving three groups were performed using one-way analysis of variance (ANOVA) followed by Šídák's post hoc multiple-comparison test, or the Kruskal-Wallis test followed by Dunn's multiple-comparison test for non-parametric data. Where missing values precluded standard repeated-measures

ANOVA, mixed-effects models (REML) with Fisher's LSD post hoc test were used. Categorical variables were analysed using $\chi^2$ tests when expected cell counts met standard assumptions; otherwise, Fisher's exact tests were used. Current-voltage relationships were analysed using linear mixed-effects models, with global fixed effects assessed by Wald $\chi^2$ tests and voltage-specific between-genotype differences assessed using two-sided Wald contrasts with Holm adjustment across test voltages. Statistical significance was defined as a two-sided $P < 0.05$.

Differential expression analysis of bulk RNA-sequencing datasets was performed using DESeq2 with Benjamini-Hochberg FDR correction. Xenium spatial transcriptomic data were analysed using Seurat and Squidpy. Cell type-specific differential expression was performed on counts using Mann-Whitney *U* tests with Benjamini-Hochberg correction for multiple testing. Differential protein abundance from mass spectrometry-based proteomic data was assessed using the limma framework with Benjamini-Hochberg FDR correction. N denotes independent biological replicates (donors or animals); n denotes cells, impalements or individual measurements, as specified. Sample sizes, statistical tests, post hoc comparisons, and exact P values are provided in the corresponding figure legends.

### Supplemental Tables

**Supplemental Table 1. Demographic and clinical characteristics of human donor cohorts**

| Experimental cohort | Donor ID | Age (years) | Sex | Donor type | Cause of death | Reported medical history |
|---|---|---|---|---|---|---|
| **Xenium spatial transcriptomics - Aged** | DN01 | 61 | F | DBD | Hypoxic brain damage | Hypertension |
| | DN02 | 73 | F | DBD | Intracranial haemorrhage | EBV$^+$, CMV$^+$, |
| | DN03 | 76 | F | DBD | Intracranial haemorrhage | Toxoplasma$^+$ |
| | DN04 | 65 | M | DBD | Intracranial haemorrhage | EBV$^+$, CMV$^+$, hypotension |
| | DN05 | 72 | F | DCD | Intracranial haemorrhage | EBV$^+$, Dilated LV, impaired systolic function, EF 41% |
| | DN06 | 80 | M | DBD | Hypoxic brain damage | EBV$^+$, CMV$^+$, Toxoplasma$^+$, hypertension, EF 55% |
| **Xenium spatial transcriptomics - Young** | DN07 | 36 | F | DBD | Intracranial haemorrhage | EBV$^+$, hypertension, hypotension, mitral valve regurgitation, LV hypertrophy, aortic stenosis |
| | DN08 | 16 | F | DBD | Hypoxic brain damage | EBV$^+$, CMV$^+$, |
| | DN09 | 27 | F | DBD | Intracranial haemorrhage | - |
| | DN10 | 27 | M | DBD | Intracranial haemorrhage | - |
| | DN11 | 33 | F | DBD | Subarachnoid haemorrhage | EBV$^+$, CMV$^+$, Toxoplasma$^+$, anterolateral ST elevation |
| ***Figure 1, 2, Supplemental Figure 1-3*** | DN12 | 29 | M | DCD | Intracranial haemorrhage | EBV$^+$, CMV$^+$, pulmonary disease |
| **SAN slices** | DN13 | 59 | F | DCD | Intracranial thrombosis | EBV$^+$, CMV$^+$, EF 56% |
| | DN14 | 58 | M | DCD | Intracranial thrombosis | EBV$^+$, CMV$^+$ |
| | DN15 | 30 | F | DBD | Intracranial haemorrhage | EBV$^+$, EF 41% |
| | DN16 | 46 | M | DBD | Intracranial haemorrhage | EBV$^+$, EF 50% myotonic dystrophy, previous cardiac arrest |
| | DN17 | 36 | F | DBD | Intracranial haemorrhage | EBV$^+$, EF 52%, treated inflammatory episode |
| | DN18 | 76 | F | DCD | Intracranial haemorrhage | EF 55%, hypertension, previous RV failure |

| | | | | | | |
|---|---|---|---|---|---|---|
| ***Figure 6, Supplememtal Figure 11*** | DN19 | 69 | M | DBD | Subarachnoid haemorrhage | Asthma, basal cell carcinoma, prostate cancer |
| **qPCR additional donors** | DN20 | 64 | M | DCD | Intracranial haemorrhage | $EBV^+$ |
| **- Aged** | DN21 | 64 | M | DBD | Intracranial thrombosis | $EBV^+$, $CMV^+$ |
| | DN22 | 78 | F | DBD | Intracranial haemorrhage | $EBV^+$, $TOXO^+$, PMH breast cancer, COPD |
| | DN23 | 61 | F | DBD | Intracranial haemorrhage | EF 65%, COPD |
| | DN24 | 74 | F | DCD | Intracranial haemorrhage | $EBV^+$, $CMV^+$ |
| | DN25 | 67 | M | DBD | Intracranial haemorrhage | $EBV^+$, $CMV^+$, $E.Coli^+$ |
| **- Young** | DN26 | 45 | F | DBD | Subarachnoid haemorrhage | $EBV^+$, Influenza |
| | DN27 | 36 | F | DBD | Intracranial haemorrhage | $EBV^+$, EF 52%, treated inflammatory episode |
| | DN28 | 21 | F | DCD | Hypoxic brain injury | $EBV^+$, Hypertension |
| | DN29 | 30 | M | DCD | Hypoxic brain injury | Hypertension |
| | DN30 | 44 | F | DBD | Intracranial haemorrhage | $EBV^+$, $CMV^+$, Hypertension |
| ***Supplemental Figure 4C, 10F*** | DN31 | 34 | F | DBD | Intracranial haemorrhage | $EBV^+$, $CMV^+$ |

Summary of donor age, sex, donor classification, cause of death, reported medical history, tissue-retrieval information and allocation to the human SAN ageing or slice cohorts. Donors were classified as donation after brainstem death (DBD) or donation after circulatory death (DCD). Reported medical history reflects information available from donor records and was not uniformly complete. Dashes indicate that no relevant information was reported. EBV, Epstein–Barr virus; CMV, cytomegalovirus; EF, ejection fraction; LV, left ventricle; RV, right ventricle

**Supplemental Table 2. Electrophysiological characteristics of freshly prepared adult human sinoatrial node tissue compared with published human recordings**

| Parameter | Tsutsui et al., 2018 | Verkerk et al., 2007 | Drouin et al., 1997 | Current study |
|---|---|---|---|---|
| **Preparation** | Isolated SAN myocytes; whole-cell patch clamp | Isolated SAN myocytes; whole-cell patch clamp | Intact SAN preparation; sharp microelectrode | 500 µm SAN slices; sharp microelectrode |
| **Donors (N)**<br>**Samples (n)** | 2<br>4 spontaneously active cells | 1<br>3 cells | 4<br>4 preparations | 7<br>18 impalements |
| **Maximum diastolic/resting membrane potential (mV)** | −50.1 ± 2.3 | −61.7 ± 4.3 | −62.5 ± 4.7 | −45.3 ± 10.1 |
| **Spontaneous cycle length (ms)** | 1698 ± 482 | 828 ± 15 | 2300 ± 300 | 2177 ± 1257 |

| | | | | |
|---|---|---|---|---|
| **Maximum upstroke velocity (V/s)** | — | 4.6 ± 1.2 | 5.4 ± 2.2 | 1.36 ± 1.05 |
| **Action potential amplitude (mV)** | — | 78.0 ± 4.5 | 72.0 ± 2.9 | 34.9 ± 12.7 |
| **$APD_{90}$ (ms)** | — | 143.5 ± 34.9 | 308.7 ± 15.5 | 527.8 ± 129.6 |

Values are mean ± SEM for Tsutsui et al.[43] and Verkerk et al.[44] and mean ± SD for Drouin et al.[45] and the current study, as reported in the respective studies. Current-study values are calculated across fresh impalements; the corresponding spontaneous cycle length reported in the Results as median [IQR] was 2291 [1211–3502] ms. Maximum diastolic potential and resting membrane potential are presented in the same row to facilitate comparison but reflect the terminology used in the respective studies. Dashes indicate parameters that were not reported. $APD_{90}$, action potential duration at 90% repolarisation; SAN, sinoatrial node.

**Supplemental Table 3. Custom 100-gene Xenium add-on panel for spatial transcriptomic profiling of the human SAN**

| Gene | Pathway | Ensembl ID |
|---|---|---|
| HCN1 | Pacemaker myocytes | ENSG00000164588 |
| HCN4 | Pacemaker myocytes | ENSG00000138622 |
| CACNA1D | Pacemaker myocytes | ENSG00000157388 |
| TBX3 | Pacemaker myocytes | ENSG00000135111 |
| TBX5 | Pacemaker myocytes | ENSG00000089225 |
| SLC8A1 | Pacemaker myocytes | ENSG00000183023 |
| BMP5 | Pacemaker myocytes | ENSG00000112175 |
| PDE1A | Pacemaker myocytes | ENSG00000115252 |
| TRPM3 | Pacemaker myocytes | ENSG00000083067 |
| GJA1 | Atrial myocytes | ENSG00000152661 |
| NPPA | Atrial myocytes | ENSG00000175206 |
| MYL7 | Atrial myocytes | ENSG00000106631 |
| PLN | Atrial myocytes | ENSG00000198523 |
| TACC2 | Atrial myocytes | ENSG00000138162 |
| MYH6 | Atrial myocytes | ENSG00000197616 |
| TTN | Atrial myocytes | ENSG00000155657 |
| TNNT2 | Atrial myocytes | ENSG00000118194 |
| HCN2 | Ion channel and related genes | ENSG00000099822 |
| SCN5A | Ion channel and related genes | ENSG00000183873 |
| CACNA1C | Ion channel and related genes | ENSG00000151067 |
| CACNA1G | Ion channel and related genes | ENSG00000006283 |
| CACNA1H | Ion channel and related genes | ENSG00000196557 |
| KCNA4 | Ion channel and related genes | ENSG00000182255 |
| KCND2 | Ion channel and related genes | ENSG00000184408 |
| KCND3 | Ion channel and related genes | ENSG00000171385 |
| KCNA5 | Ion channel and related genes | ENSG00000130037 |
| KCNH2 | Ion channel and related genes | ENSG00000055118 |
| KCNQ1 | Ion channel and related genes | ENSG00000053918 |
| KCNJ2 | Ion channel and related genes | ENSG00000123700 |
| KCNJ12 | Ion channel and related genes | ENSG00000184185 |
| KCNJ3 | Ion channel and related genes | ENSG00000162989 |
| KCNJ5 | Ion channel and related genes | ENSG00000120457 |
| ATP2A2 | Ion channel and related genes | ENSG00000174437 |
| CASQ2 | Ion channel and related genes | ENSG00000118729 |
| GJD3 | Ion channel and related genes | ENSG00000183153 |
| GJA5 | Ion channel and related genes | ENSG00000265107 |
| GJC1 | Ion channel and related genes | ENSG00000182963 |
| CHRM2 | Ion channel and related genes | ENSG00000181072 |
| ADRB1 | Ion channel and related genes | ENSG00000043591 |
| ADRB2 | Ion channel and related genes | ENSG00000169252 |

| | | |
|---|---|---|
| GATA5 | Girk4 | ENSG00000130700 |
| AKT3 | Girk4 | ENSG00000117020 |
| GATA6 | Girk4 | ENSG00000141448 |
| LGALS3 | Galectin-3 | ENSG00000131981 |
| IL11 | Fibroblasts | ENSG00000095752 |
| COL3A1 | Fibroblasts | ENSG00000168542 |
| COL1A2 | Fibroblasts | ENSG00000164692 |
| POSTN | Fibroblasts | ENSG00000133110 |
| PI16 | Fibroblasts | ENSG00000164530 |
| COL6A3 | Fibroblasts | ENSG00000163359 |
| MMP2 | Fibroblasts | ENSG00000087245 |
| RGS4 | Myofibroblasts | ENSG00000117152 |
| COLEC11 | Myofibroblasts | ENSG00000118004 |
| MYH11 | Smooth muscle | ENSG00000133392 |
| CALD1 | Smooth muscle | ENSG00000122786 |
| MYL9 | Smooth muscle | ENSG00000101335 |
| CNN1 | Smooth muscle | ENSG00000130176 |
| VWF | Endothelial cells | ENSG00000110799 |
| PECAM1 | Endothelial cells | ENSG00000261371 |
| PTPRB | Endothelial cells | ENSG00000127329 |
| ADAMTS9 | Endothelial cells | ENSG00000163638 |
| FABP4 | Adipocytes | ENSG00000170323 |
| RETN | Adipocytes | ENSG00000104918 |
| GPAM | Adipocytes | ENSG00000119927 |
| FASN | Adipocytes | ENSG00000169710 |
| TUBB3 | Neurons | ENSG00000258947 |
| NRXN1 | Neurons | ENSG00000179915 |
| MAP2 | Neurons | ENSG00000078018 |
| SYN1 | Neurons | ENSG00000008056 |
| TNFAIP6 | RNAseq | ENSG00000123610 |
| TNFSF13B | RNAseq | ENSG00000102524 |
| TRPV3 | RNAseq | ENSG00000167723 |
| ACKR2 | RNAseq | ENSG00000144648 |
| ACKR3 | RNAseq | ENSG00000144476 |
| FOXO1 | RNAseq | ENSG00000150907 |
| NLRP3 | NLRP3 inflammasome | ENSG00000162711 |
| RELA | TNF pathway | ENSG00000173039 |
| TNFRSF1A | TNF pathway | ENSG00000067182 |
| TRAF2 | TNF pathway | ENSG00000127191 |
| PIK3R1 | TNF pathway | ENSG00000145675 |
| TRADD | TNF pathway | ENSG00000102871 |
| MAPK8 | TNF pathway | ENSG00000107643 |
| MAPK9 | TNF pathway | ENSG00000050748 |
| MAPK10 | TNF pathway | ENSG00000109339 |
| NFKBIA | TNF pathway | ENSG00000100906 |
| NFKB1 | TNF pathway | ENSG00000109320 |
| IKBKB | TNF pathway | ENSG00000104365 |
| IKBKG | TNF pathway | ENSG00000269335 |
| CHUK | TNF pathway | ENSG00000213341 |
| PYCARD | NLRP3 inflammasome | ENSG00000103490 |
| CASP1 | NLRP3 inflammasome | ENSG00000137752 |
| TLR1 | TLR signalling | ENSG00000174125 |
| TLR2 | TLR signalling | ENSG00000137462 |
| TLR6 | TLR signalling | ENSG00000174130 |
| RAC1 | TLR signalling | ENSG00000136238 |
| MAP2K1 | TLR signalling | ENSG00000169032 |
| TLR3 | TLR signalling | ENSG00000164342 |
| MAP2K2 | TLR signalling | ENSG00000126934 |

| | | |
|---|---|---|
| MAPK1 | TLR signalling | ENSG00000100030 |
| MAPK3 | TLR signalling | ENSG00000102882 |

Gene are grouped by biological category and pre-selected based on relevance to SAN function, cardiac fibrosis, and inflammatory signalling pathways, to supplement the Xenium Human Immuno-Oncology panel (380 genes). Ensembl ID indicated the references transcript used for probe design.

**Supplemental Table 4. Number of cells from each donor following Xenium cell segmentation.**

| Donor | Segmented cells |
|---|---|
| H100 | 57,675 |
| H111 | 67,240 |
| DN40 | 38,744 |
| DN35 | 18,215 |
| DN36 | 21,568 |
| DN37 | 44,332 |
| DN39 | 24,941 |
| **Total** | **272,715** |
| **Mean ± SD** | **38,959 ± 18,745** |

**Supplemental Table 5. Antibodies and reagents used for mouse flow cytometry analysis.**

| Marker / Reagent | Fluorophore | Clone | Supplier | Catalogue No. |
|---|---|---|---|---|
| LIVE/DEAD™ Fixable Blue Dead Cell Stain Kit | NA | NA | Invitrogen | L34961 |
| MHCII | BUV395 | 2G9 | BD Biosciences | 569244 |
| TCR delta | BUV563 | GL3 | BD Biosciences | 748993 |
| CD19 | BUV661 | eBio1D3 | Invitrogen | 376-0193-80 |
| CD11c | BUV737 | HL3 | BD Biosciences | 612796 |
| CD4 | BUV805 | RM4-5 | Invitrogen | 368-0042-82 |
| Ly6C/Ly6G | eFluor 450 | RB6-8C5 | Invitrogen | 48-5931-82 |
| CD45 | BV510 | 30-F11 | BioLegend | 103137 |
| CD11b | BV711 | M1/70 | BioLegend | 101241 |
| TCR beta | BV785 | H57-597 | BioLegend | 109249 |
| Eomes | PE-Cy7 | Dan11mag | Invitrogen | 25-4875-80 |
| CD25 | PE-Cy5 | PC61 | BioLegend | 102010 |
| CCR2 | PE | SA203G11 | BioLegend | 150609 |
| CD206 | PE-Dazzle™ 594 | C068C2 | BioLegend | 141731 |
| F4/80 | Alexa Fluor 647 | BM8 | BioLegend | 123122 |
| CD44 | Alexa Fluor 700 | IM7 | BioLegend | 103026 |
| CD8α | APC-Fire™ 750 | 53-6.7 | BioLegend | 100766 |

| Marker / Reagent | Fluorophore | Clone | Supplier | Catalogue No. |
|---|---|---|---|---|
| TNF | BV650 | MP6-XT22 | Biolegend | 50633 |
| CD64 | APC | X54-5/7.1 | Biolegend | 139305 |
| Brefeldin-A solution, 1000x | NA | NA | eBioscience | 00-4506 |
| Phorbol 12-myristate 13-acetate | NA | NA | Sigma-Aldrich | P8139 |
| Ionomycin | NA | NA | Sigma-Aldrich | I9657 |
| Foxp3 / Transcription Factor Staining Buffer Set | NA | NA | eBioscience | 00-5523-00 |
| Precision Count Beads™ | NA | NA | BioLegend | 424902 |

Fluorophore-conjugated antibodies, clones, suppliers, and catalogue numbers used for immunophenotyping of mouse immune cell populations.

**Supplemental Table 6. Primary and secondary immunolabelling antibodies.**

| Target | Host species | Supplier | Catalogue No. | Primary dilution | Secondary antibody |
|---|---|---|---|---|---|
| HCN4 | Guinea pig | Alomone Labs | APC-052-GP | 1:100 | Alexa Fluor™ 647 goat anti-guinea pig IgG |
| $Ca_v1.3$ | Mouse | Novus Biologicals | NBP2-12893 | 1:200 | Alexa Fluor™ 680 goat anti-mouse IgG |
| Connexin 43 (Cx43) | Rabbit | Sigma-Aldrich | C6219 | 1:500 | Alexa Fluor™ 546 goat anti-rabbit IgG |
| CD45 | Rat | Bio-Rad | MCA1957 | 1:200 | Alexa Fluor™ 546 goat anti-rat IgG |
| TNF | Rabbit | Abcam | ab6671 | 1:100 | Alexa Fluor™ 594 goat anti-rabbit IgG |
| CD68 | Rat | Thermo Fisher | 14-0688-82 | 1:200 | Alexa Fluor™ 546 goat anti-rat IgG |
| Amphiregulin (AREG) | Rabbit | Proteintech | 16036-1-AP | 1:200 | Alexa Fluor™ 680 goat anti-rabbit IgG |
| CD4 | Rabbit | Abcam | ab183685 | 1:200 | Alexa Fluor™ 680 goat anti-rabbit IgG |
| CD8 | Rat | Invitrogen | 14-0081 | 1:200 | Alexa Fluor™ 546 goat anti-rat IgG |
| TNFR1 | Mouse | Proiteintech | 60192-1-Ig | 1:200 | Alexa Fluor™ 680 goat anti-mouse IgG |
| TNFR2 | Rabbit | Proiteintech | 19272-1-AP | 1:200 | Alexa Fluor™ 546 goat anti-rabbit IgG |
| NF-kB p65 | *594-conjugated* | Proiteintech | CL594-80979 | 1:200 | Alexa Fluor™ 594-conjugated |
| Cleaved caspase 3 | Rabbit | Proteintech | 25128-1-AP | 1:200 | Alexa Fluor™ 680 goat anti-rabbit IgG |

Host species, suppliers, catalogue numbers, primary antibody dilutions, and Alexa Fluor-conjugated secondary antibodies used for immunolabelling.

**Supplemental Table 7. Human and mouse qPCR primer assays**

| Human (Hs) | Catalogue No. | Mouse (Mm) | Catalogue No. |
|---|---|---|---|
| Hs_GAPDH_1_SG | QT00079247 | Mm_IPO8_1_SG | QT00291977 |
| Hs_HCN4_1_SG | QT00038108 | Mm_HCN4_1_SG | QT00268660 |
| Hs_HCN1_1_SG | QT0048020 | Mm_HCN1_1_SG | QT00172130 |
| Hs_CACNA1D_1_SG | QT00091812 | Mm_CACNA1D_1_SG | QT00112238 |
| Hs_SCN5A_1_SG | QT00012684 | Mm_SCN5A_1_SG | QT00136714 |
| Hs_KCNH2_2_SG | QT01003254 | Mm_KCNH2_1_SG | QT00105574 |
| Hs_TNF_1_SG | QT00029162 | Mm_KCNJ5_1_SG | QT00286216 |
| Hs_NFKB1_1_SG | QT00063791 | Mm_TNF_1_SG | QT00104006 |
| Hs_NLRP3_1_SG | QT00029771 | Mm_NFKBIA_1_SG | QT00134421 |
| Hs_IL6_1_SG | QT00083720 | Mm_NLRP3_1_SG | QT01552033 |
| Hs_CCL2_1_SG | QT00212730 | Mm_IL6_1_SG | QT00098875 |
| Hs_TBX3_1_SG | QT00022484 | Mm_CCL2_1_SG | QT00167832 |
| | | Mm_TBX3_1_SG | QT00147042 |

Qiagen QuantiTect primer assay IDs and catalogue numbers.

**Supplemental Table 8. Primary and secondary antibodies used for western blotting**

| Antibody | Host species | Supplier | Catalogue No. | Dilution |
|---|---|---|---|---|
| HCN4 | Rabbit | Alomone Labs | APC-052 | 1:500 |
| HCN1 | Rabbit | Alomone Labs | APC-056 | 1:500 |
| KCNH2 | Rabbit | Alomone Labs | OSP00150W | 1:500 |
| TNF | Rabbit | Alomone Labs | ALX-210-909 | 1:500 |
| Connexin 43 | Mouse | Invitrogen | ACI-201 | 1:500 |
| Connexin 45 | Mouse | Invitrogen | 41-4580 | 1:250 |
| HRP-conjugated anti-rabbit IgG | Goat | Cell Signaling Technology | 7074 | 1:3000 |
| HRP-conjugated anti-mouse IgG | Goat | Santa Cruz | Sc-516102 | 1:3000 |

Host species, suppliers, catalogue numbers, and working dilutions of antibodies used for western blot analysis.

**Supplemental Table 9. Voltage-specific analysis of pacemaker ionic currents in ageing and HCN4/TNF$^{glo}$ SAN myocytes.**

| Test voltage (mV) | Young, mean±SEM (pA/pF) | Aged, mean±SEM (pA/pF) | Šídák-adjusted P |
|---|---|---|---|
| −55 | −1.09±0.30 | −0.90±0.09 | >.9999 |
| −65 | −1.79±0.50 | −0.98±0.21 | >.9999 |
| −75 | −4.20±0.84 | −2.76±0.44 | 0.9999 |
| −85 | −9.53±1.28 | −5.43±0.78 | 0.8044 |
| −95 | −17.00±1.89 | −8.89±1.18 | 0.0543 |
| −105 | −25.29±2.34 | −12.54±1.86 | 0.00015 |
| −115 | −32.46±2.21 | −16.30±2.14 | $4.32\times10^{-7}$ |
| −125 | −42.00±3.86 | −20.84±2.78 | $8.16\times10^{-12}$ |
| −135 | −50.33±3.93 | −25.20±3.69 | $<1\times10^{-15}$ |
| **$I_f$** | | | |
| **Test voltage (mV)** | **Control, mean±SEM (pA/pF)** | **HCN4/TNF$^{glo}$, mean±SEM (pA/pF)** | **Šídák-adjusted P** |
| −135 | −42.79±3.43 | −24.21±2.40 | 0.0003 |
| −125 | −34.11±2.42 | −20.09±1.91 | 0.0002 |
| −115 | −27.05±1.82 | −16.64±1.61 | 0.0005 |
| −105 | −21.22±1.41 | −13.27±1.34 | 0.001 |
| −95 | −15.11±1.05 | −9.29±1.05 | 0.0018 |
| −85 | −9.39±0.76 | −5.49±0.71 | 0.0033 |
| −75 | −4.93±0.45 | −2.64±0.41 | 0.0028 |
| −65 | −2.47±0.24 | −1.20±0.19 | 0.0009 |
| −55 | −1.41±0.17 | −0.77±0.10 | 0.0209 |
| −45 | −1.31±0.16 | −0.76±0.09 | 0.0391 |
| −35 | −1.21±0.16 | −0.76±0.10 | 0.2039 |
| **Total $I_{Ca,L}$** | | | |
| **Test voltage (mV)** | **Control, mean±SEM (pA/pF)** | **HCN4/TNF$^{glo}$, mean±SEM (pA/pF)** | **Šídák-adjusted P** |
| −55 | −0.45±0.04 | −0.37±0.03 | 0.7839 |
| −50 | −0.39±0.05 | −0.33±0.03 | 0.9993 |
| −45 | −0.51±0.11 | −0.33±0.04 | 0.8749 |
| −40 | −1.11±0.21 | −0.44±0.05 | 0.0527 |
| −35 | −2.14±0.34 | −0.65±0.08 | 0.0028 |
| −30 | −3.61±0.53 | −1.12±0.13 | 0.0012 |
| −25 | −5.09±0.65 | −1.86±0.21 | 0.0007 |
| −20 | −6.64±0.72 | −2.94±0.29 | 0.0005 |
| −10 | −8.54±0.65 | −5.61±0.41 | 0.0061 |
| 0 | −8.77±0.49 | −7.61±0.42 | 0.7082 |
| 10 | −6.91±0.40 | −7.53±0.43 | 0.9952 |

| | | | |
|---|---|---|---|
| 20 | −4.30±0.44 | −6.19±0.42 | 0.0452 |
| 40 | −0.71±0.39 | −1.98±0.34 | 0.228 |
| 50 | 0.99±0.37 | −0.19±0.29 | 0.2038 |
| 60 | 2.62±0.39 | 1.04±0.27 | 0.022 |
| $I_{Cav1.2}$ | | | |
| **Test voltage (mV)** | **Control, mean±SEM (pA/pF)** | **HCN4/TNF$^{glo}$, mean±SEM (pA/pF)** | **Šídák-adjusted P** |
| −55 | 0.03±0.04 | 0.04±0.02 | >.9999 |
| −50 | 0.11±0.07 | 0.02±0.02 | 0.986 |
| −45 | 0.26±0.09 | 0.02±0.04 | 0.3228 |
| −40 | 0.13±0.13 | −0.04±0.05 | 0.9824 |
| −35 | −0.11±0.21 | −0.10±0.08 | >.9999 |
| −30 | −0.83±0.25 | −0.40±0.11 | 0.8685 |
| −25 | −1.66±0.29 | −0.83±0.17 | 0.25 |
| −20 | −2.87±0.36 | −1.60±0.24 | 0.0738 |
| −10 | −4.50±0.37 | −3.49±0.39 | 0.6496 |
| 0 | −5.26±0.35 | −5.02±0.48 | >.9999 |
| 10 | −4.39±0.36 | −5.22±0.48 | 0.9445 |
| 20 | −3.32±0.34 | −4.36±0.41 | 0.5958 |
| 40 | −1.54±0.27 | −2.08±0.29 | 0.9509 |
| 50 | −0.79±0.24 | −1.29±0.24 | 0.9147 |
| 60 | −0.41±0.28 | −0.77±0.22 | 0.9965 |
| $I_{Cav1.3}$ | | | |
| **Test voltage (mV)** | **Control, mean±SEM (pA/pF)** | **HCN4/TNF$^{glo}$, mean±SEM (pA/pF)** | **Šídák-adjusted P** |
| −55 | −0.50±0.05 | −0.39±0.04 | 0.6934 |
| −50 | −0.53±0.08 | −0.32±0.04 | 0.3386 |
| −45 | −0.85±0.10 | −0.33±0.05 | 0.0016 |
| −40 | −1.37±0.22 | −0.36±0.06 | 0.0021 |
| −35 | −2.22±0.34 | −0.48±0.08 | 0.0007 |
| −30 | −3.05±0.46 | −0.66±0.11 | 0.0005 |
| −25 | −3.77±0.55 | −0.99±0.15 | 0.0008 |
| −20 | −4.12±0.58 | −1.31±0.18 | 0.0015 |
| −10 | −4.24±0.55 | −2.16±0.24 | 0.0245 |
| 0 | −3.58±0.43 | −2.81±0.31 | 0.9135 |
| 10 | −2.56±0.32 | −2.70±0.34 | >.9999 |
| 20 | −1.15±0.29 | −2.11±0.35 | 0.4575 |

| | | | |
|---|---|---|---|
| 40 | 0.75±0.29 | 0.01±0.29 | 0.7162 |
| 50 | 1.66±0.29 | 1.03±0.29 | 0.869 |
| 60 | 2.82±0.33 | 1.79±0.28 | 0.2844 |

Current densities and voltage-specific comparisons for the current–voltage relationships shown in Figures 3C and 5C. Data are mean±SEM and current density is expressed as pA/pF. Current–voltage relationships were analysed using mixed-effects models with group, test voltage and their interaction, followed by Šídák-adjusted multiple comparisons between groups at individual voltages. For young versus aged *I*f (Figure 3C): young, n=16 cells/N=6 mice; aged, n=16/N=5. For control vs HCN4/TNF$^{glo}$ currents $I_f$: control, n=46 cells/N=9 mice; HCN4/TNF$^{glo}$, n=54/N=13. For total $I_{Ca,L}$: 27/N=5 versus 35/N=6; $I$Ca$_V$1.2: 23/N=4 versus 27/N=5; and $I$Ca$_V$1.3: 23/N=4 versus 29/N=5. *n* denotes cells and *N* denotes mice. Adjusted two-sided *P* values are shown.

### Supplemental Results

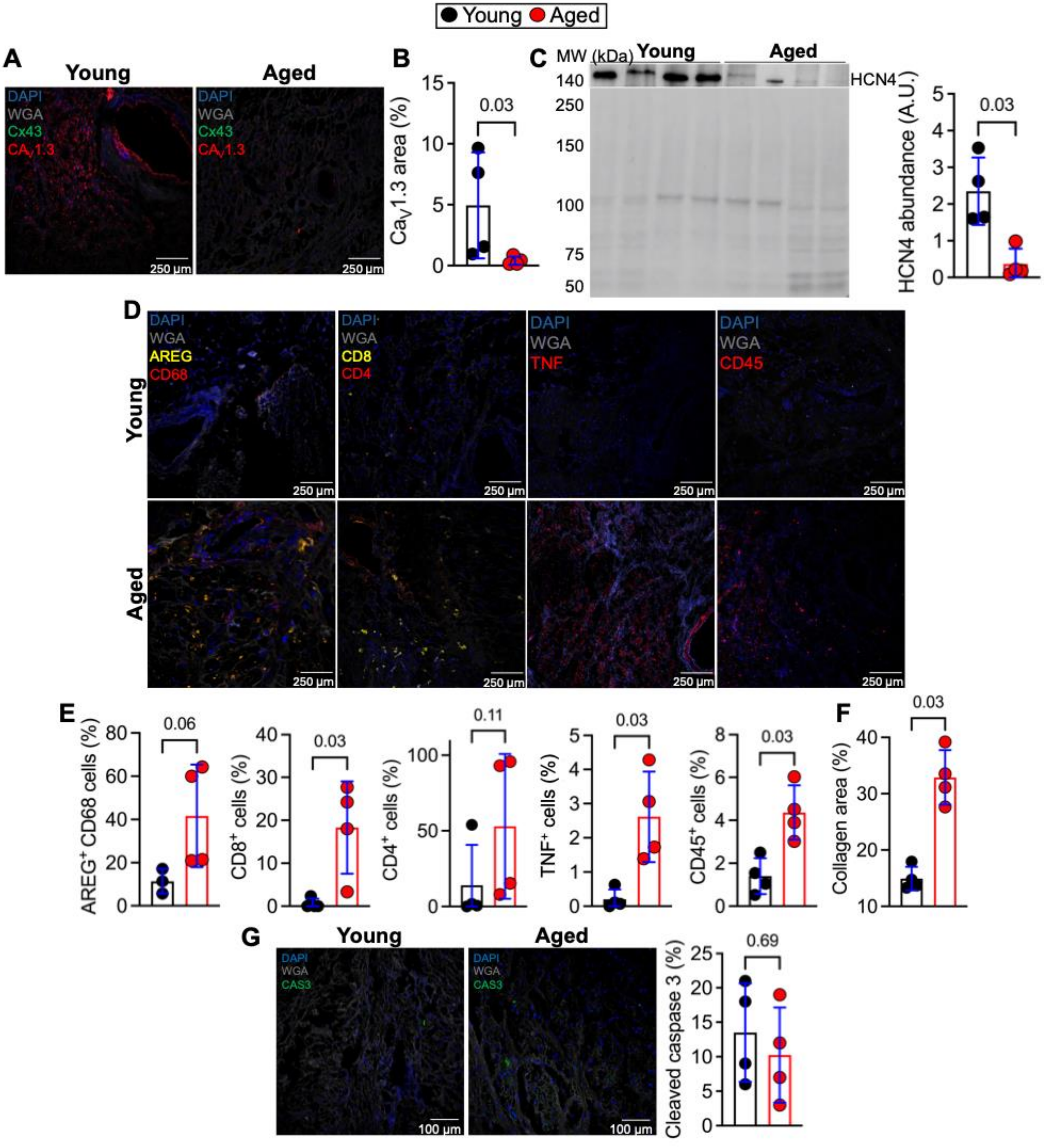

**Supplemental Figure 1. Identification and immune characterisation of the human SAN in young and aged donors (A)** Representative immunofluorescence images used to identify human SAN region in donor hearts. Images acquired at 10× magnification show nuclei (DAPI, blue), cardiomyocyte membranes (wheat germ agglutinin, WGA, grey), connexin 43 (Cx43, green), and $Ca_v1.3$ (red). Cx43-negative, $Ca_v1.3$-enriched regions were used to define the SAN region. Scale bars, 250 µm. **(B)** Quantification of the percentage of $Ca_v1.3^+$ area in the SAN from young (N=4) and aged (N=4) donors. **(C)** Representative western blot and quantification of stain-free total protein normalised HCN4 protein abundance in SAN tissue from young (N=4) and aged (N=4) donors. **(D)** Representative immunofluorescence images acquired at 10x showing amphiregulin (AREG, yellow) and CD68 (red), CD8 (yellow) and CD4 (red), TNF (red), and CD45 (red), together with WGA (grey) and DAPI (blue), in young and aged human SAN tissue. Scale bars, 250 µm. **(E)** Quantification of the percentage of $AREG^+$ $CD68^+$ macrophages, $CD8^+$ T cells, $CD4^+$ T cells, $CD45^+$ leukocytes, and $TNF^+$ cells in young (N=4) and aged (N=4) donors. **(F)** Quantification of collagen area fraction in SAN tissue from young (N=4) and aged (N=4) donors from Masson's trichrome-stained sections. **(G)** Representative immunofluorescence images of human SAN tissue from young (N=4) and aged (N=4) donors showing cleaved caspase-3 (CAS3, green), WGA (grey) and nuclei (DAPI, blue). Quantification of $CAS3^+$ cells in young and aged SAN tissue is shown alongside. Scale bars, 100 µm. Data are presented as mean ± SD with individual donors. P-values computed by Mann–Whitney U test (B, C, E, F, G).

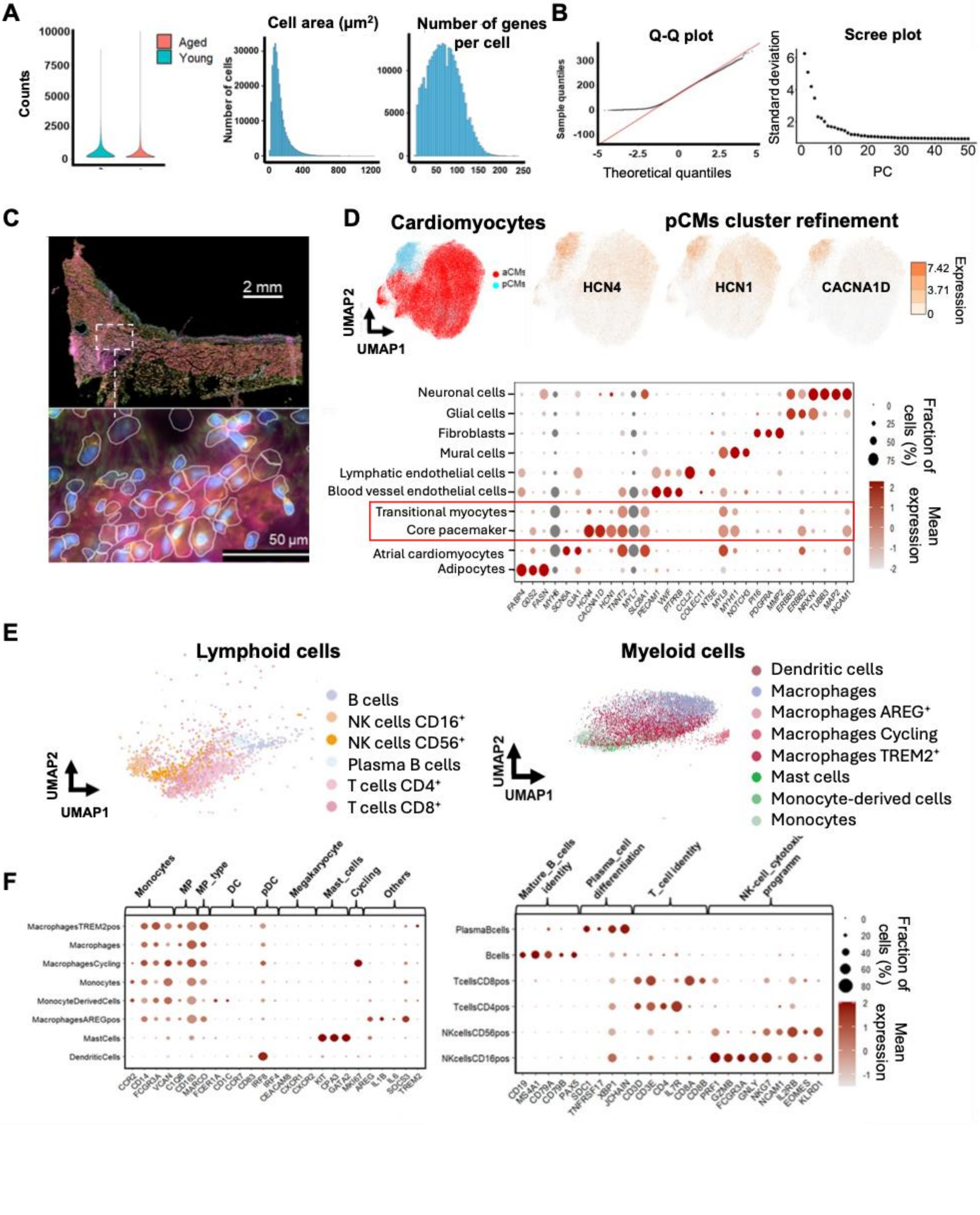
A
Counts
Aged
Young
Cell area (µm²)
Number of cells
Number of genes per cell
B
Q-Q plot
Sample quantiles
Theoretical quantiles
Scree plot
Standard deviation
PC
C
2 mm
50 µm
D
Cardiomyocytes
aCMs
pCMs
UMAP2
UMAP1
pCMs cluster refinement
HCN4
HCN1
CACNA1D
Expression
Neuronal cells
Glial cells
Fibroblasts
Mural cells
Lymphatic endothelial cells
Blood vessel endothelial cells
Transitional myocytes
Core pacemaker
Atrial cardiomyocytes
Adipocytes
Fraction of cells (%)
Mean expression
E
Lymphoid cells
B cells
NK cells CD16⁺
NK cells CD56⁺
Plasma B cells
T cells CD4⁺
T cells CD8⁺
UMAP2
UMAP1
Myeloid cells
Dendritic cells
Macrophages
Macrophages AREG⁺
Macrophages Cycling
Macrophages TREM2⁺
Mast cells
Monocyte-derived cells
Monocytes
F
Monocytes
MP
MP_type
DC
pDC
Megakaryocyte
Mast_cells
Cycling
Others
MacrophagesTREM2pos
Macrophages
MacrophagesCycling
Monocytes
MonocyteDerivedCells
MacrophagesAREGpos
MastCells
DendriticCells
Mature_B_cells identity
Plasma_cell differentiation
T_cell identity
NK-cell_cytotoxic programm
PlasmaBcells
Bcells
TcellsCD8pos
TcellsCD4pos
NKcellsCD56pos
NKcellsCD16pos
Fraction of cells (%)
Mean expression

**Supplemental Figure 2. Quality control and cell-type annotation of human SAN Xenium spatial transcriptomic datasets (A)** Quality control metrics for Xenium datasets derived from young (N=3) and aged (N=4) human SAN samples. Left: distribution of total transcripts detected per sample. Middle: distribution of segmented cell area (µm²). Right: distribution of genes detected per cell. **(B)** Assessment of data distribution and dimensionality reduction. Left: quantile–quantile (Q–Q) plot comparing observed and theoretical distributions of genes detected per cell. The black dotted line represents observed values, and the red line indicates the expected normal distribution. Right: scree plot showing the contribution of individual principal components to overall dataset variance. **(C)** Representative Xenium spatial transcriptomic image of the human SAN demonstrating transcript localisation and cell segmentation. Upper panel: overview of the SAN region analysed. Lower panel: high-magnification view showing segmented cell boundaries and transcript distribution. Scale bars as indicated. **(D)** Identification and annotation of cardiomyocyte populations. Left: UMAP embedding of cardiomyocytes classified as atrial cardiomyocytes (aCMs, red) and pacemaker cardiomyocytes (pCMs, cyan). Upper right: feature plots demonstrating expression of canonical pacemaker markers HCN4, HCN1, and CACNA1D. Lower right: dot plot showing marker genes used for annotation of non-immune cell populations. Dot size represents the percentage of cells expressing each gene and colour indicates mean expression. **(E)** UMAP embeddings of immune cell populations identified within the human SAN. Left: lymphoid cell populations, including B cells, plasma cells, NK cells, and $CD4^+$ and $CD8^+$ T-cell subsets. Right: myeloid cell populations, including macrophages, monocytes, monocyte-derived cells, dendritic cells, mast cells, and macrophage-state subclusters. **(F)** Dot plots showing canonical marker genes used for immune cell annotation. Left: myeloid cell populations. Right: lymphoid cell populations. Dot size represents the percentage of cells expressing each gene and colour indicates mean expression.

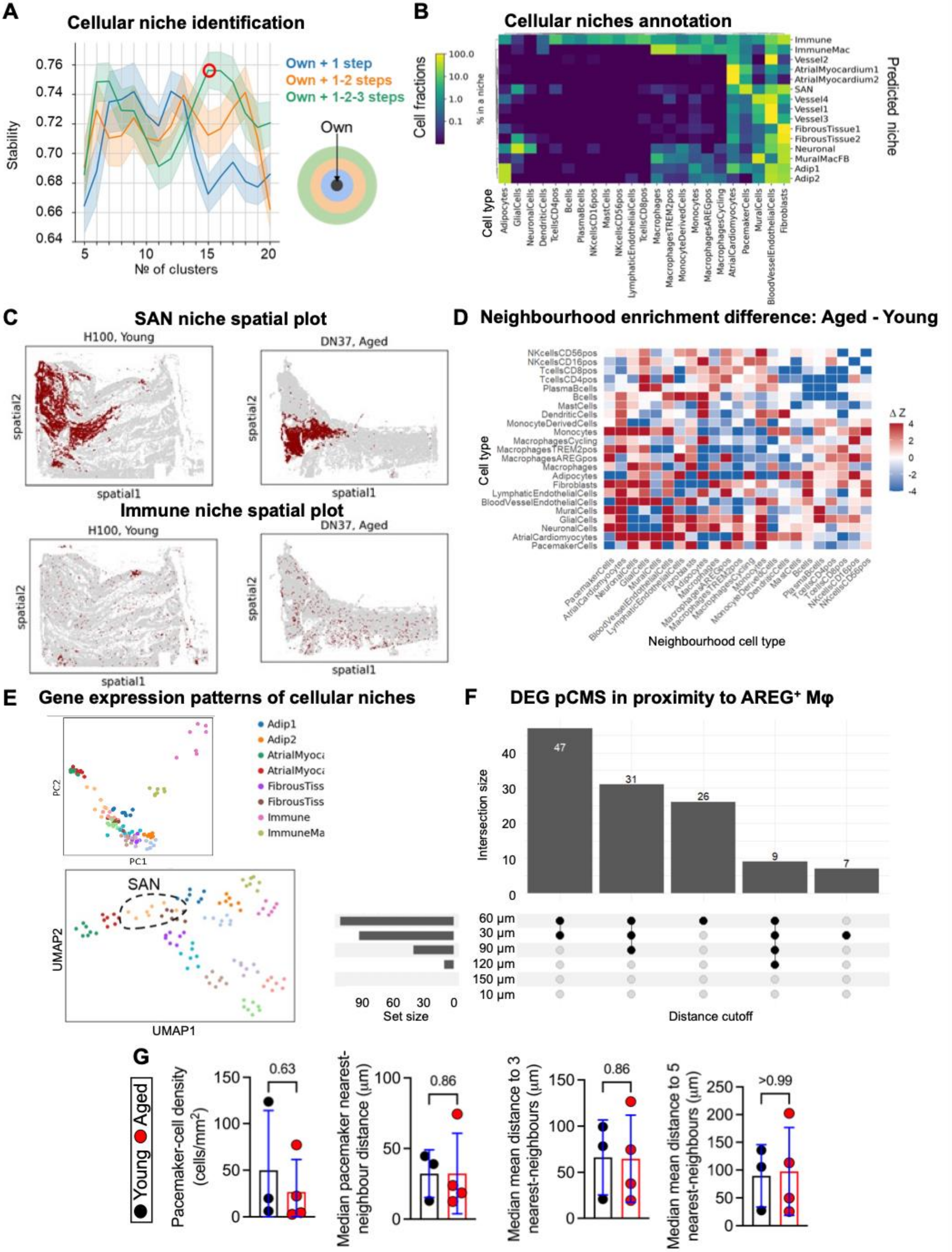

A
Cellular niche identification
Own + 1 step
Own + 1-2 steps
Own + 1-2-3 steps
Own
Stability
№ of clusters
B
Cellular niches annotation
Cell fractions
% in a niche
Predicted niche
Cell type
C
SAN niche spatial plot
H100, Young
DN37, Aged
Immune niche spatial plot
spatial1
spatial2
D
Neighbourhood enrichment difference: Aged - Young
Cell type
Neighbourhood cell type
ΔZ
E
Gene expression patterns of cellular niches
PC1
PC2
UMAP1
UMAP2
SAN
F
DEG pCMS in proximity to AREG⁺ Mφ
Intersection size
Set size
Distance cutoff
G
Young
Aged
Pacemaker-cell density (cells/mm²)
Median pacemaker nearest-neighbour distance (µm)
Median mean distance to 3 nearest-neighbours (µm)
Median mean distance to 5 nearest-neighbours (µm)

**Supplemental Figure 3. Identification, annotation, and spatial characterisation of cellular niches in the human SAN (A)** Spatial niche identification and cluster stability analysis. Cluster stability was assessed across increasing neighbourhood aggregation depths, incorporating information from a cell and its first-order neighbours (blue), first- and second-order neighbours (orange), and broader tissue context including first-, second-, and third-order neighbours (green). The optimal clustering solution was selected based on maximal cluster stability (red circle). Schematic illustrating neighbourhood aggregation depths is shown on the right. **(B)** Annotation of spatial niches based on previously identified cell populations. Heatmap showing the percentage contribution of each cell type to individual spatial niches, with colour intensity indicating the proportion of cells within each niche. **(C)** Representative spatial localisation of cellular niches within young and aged human SAN tissue. Top: spatial distribution of the SAN niche in representative young and aged samples. Bottom: spatial distribution of the Immune niche in representative young and aged samples. Cells belonging to the indicated niche are shown in red. **(D)** Differences in permutation-derived spatial neighbourhood-enrichment Z-scores between aged and young human SAN sections. These scores describe relative spatial associations within the profiled tissue. Colour intensity represents the difference in neighbourhood enrichment Z-scores ($\Delta Z$) between aged and young tissues. Red indicates increased spatial association in aged human SAN, whereas blue indicates reduced spatial association relative to young human SAN. **(E)** Dimensionality reduction of pseudobulk spatial niche profiles. Top: principal component analysis (PCA) of donor-level cellular niches. Bottom: UMAP embedding of cellular niches identified across human SAN samples. Each point represents an individual donor niche. The SAN niche is highlighted by the dashed black outline. **(F)** Sensitivity analysis of cell-level differential expression in pacemaker cardiomyocytes classified by proximity to $AREG^+$ macrophages across distance thresholds of 10, 30, 60, 90, 120 and 150 µm.. Bar plots indicate the number of genes shared between distance thresholds, while connected dots indicate the threshold combinations contributing to each gene set intersection. **(G)** Pacemaker cardiomyocyte density (cells per $mm^2$ of tissue), nearest-neighbour distance between pacemaker cardiomyocytes, and mean distance to the three and five nearest pacemaker cardiomyocyte neighbours, in young (N=3) and aged (N=4) human SAN donors. Data are presented as mean ± SD, with individual data points representing individual donors. P-values computed by Mann–Whitney U test.

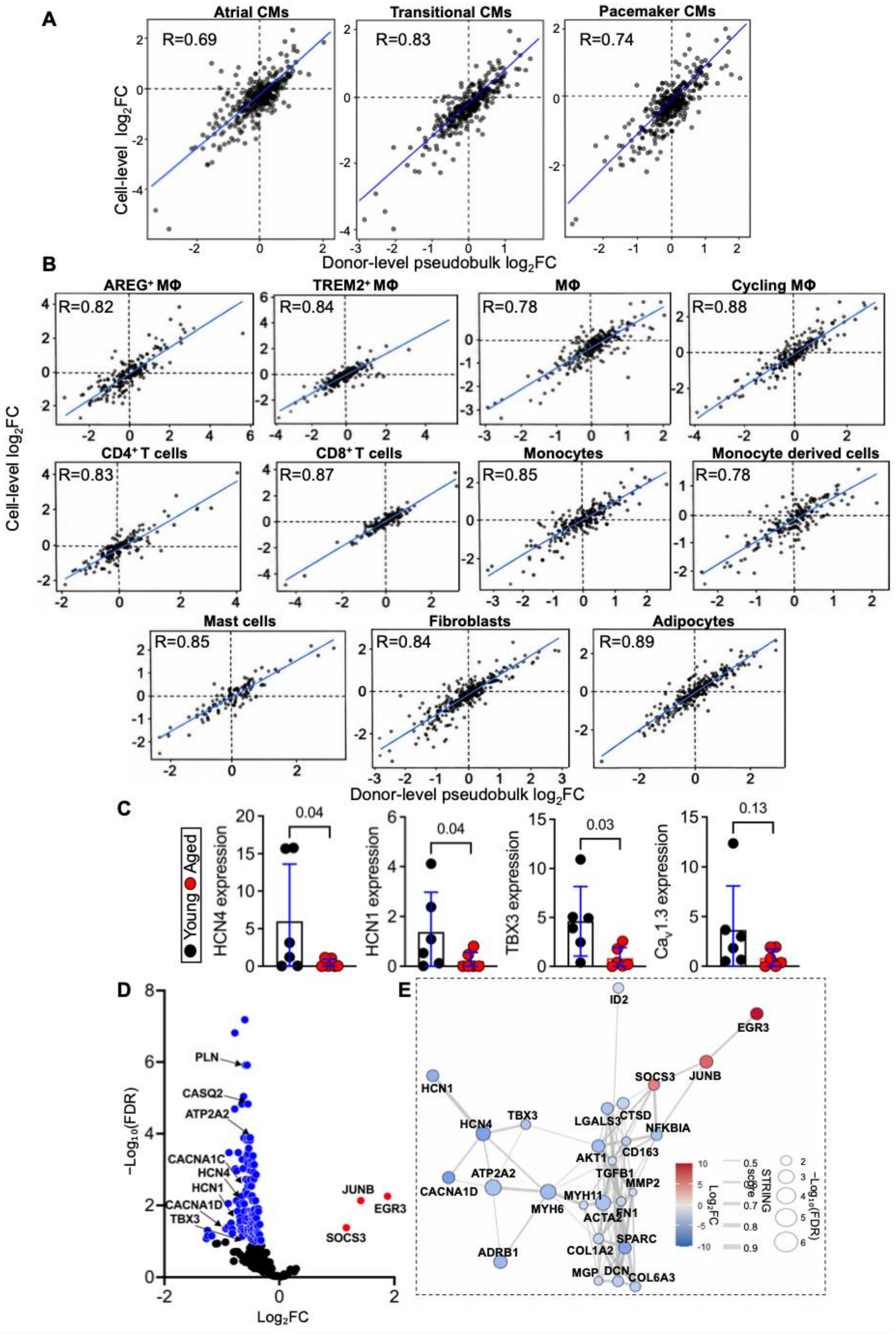

A
Atrial CMs
Transitional CMs
Pacemaker CMs
R=0.69
R=0.83
R=0.74
Cell-level log2FC
Donor-level pseudobulk log2FC
B
AREG+ MΦ
TREM2+ MΦ
MΦ
Cycling MΦ
R=0.82
R=0.84
R=0.78
R=0.88
CD4+ T cells
CD8+ T cells
Monocytes
Monocyte derived cells
R=0.83
R=0.87
R=0.85
R=0.78
Mast cells
Fibroblasts
Adipocytes
R=0.85
R=0.84
R=0.89
Cell-level log2FC
Donor-level pseudobulk log2FC
C
Young
Aged
HCN4 expression
0.04
HCN1 expression
0.04
TBX3 expression
0.03
CaV1.3 expression
0.13
D
−Log10(FDR)
Log2FC
PLN
CASQ2
ATP2A2
CACNA1C
HCN4
HCN1
CACNA1D
TBX3
JUNB
EGR3
SOCS3
E
ID2
EGR3
JUNB
SOCS3
HCN1
HCN4
TBX3
LGALS3
CTSD
NFKBIA
AKT1
CD163
TGFB1
MMP2
ATP2A2
CACNA1D
MYH6
MYH11
ACTA2
FN1
COL1A2
SPARC
ADRB1
MGP
DCN
COL6A3
Log2FC
STRING score
−Log10(FDR)

**New Supplemental Figure 4. Donor-level analysis of age-associated transcriptional changes and macrophage-proximity-associated remodelling in the human SAN. (A)** Comparison of age-associated $\log_2$ fold-change estimates obtained from cell-level Wilcoxon analysis (y-axis) and donor-level pseudobulk DESeq2 analysis (x-axis) in atrial (aCMs), transitional (tCMs) and pacemaker cardiomyocytes (pCMs) from young (N=3) and aged (N=4) human SAN donors. Raw Xenium transcript counts were aggregated by donor and cell population for pseudobulk analysis. Each point represents one gene, R denotes the Pearson correlation coefficient, dashed lines indicate zero fold change, and the blue line represents the ordinary least-squres (OLS) linear regression fit. **(B)** Equivalent comparison for immune and stromal populations, comprising $AREG^+$ macrophages, $TREM2^+$ macrophages, macrophages, cycling macrophages, $CD4^+$ T cells, $CD8^+$ T cells, monocytes, monocyte-derived cells, mast cells, fibroblasts and adipocytes. **(C)** Relative mRNA expression of *HCN4, HCN1, TBX3, CACNA1D* in human SAN tissue from young (N=6) and aged (N=6) donors, normalised to GAPDH and calculated using the $2^{-\Delta\Delta Ct}$ method. **(D)** Volcano plot showing donor-level pseudobulk differential expression between pacemaker cardiomyocytes located within 60 μm of an $AREG^+$ macrophage and those located more than 60 μm away. The x-axis shows $\log_2$ fold change (proximal relative to distal) and the y-axis $-\log_{10}$(FDR). Significantly upregulated and downregulated genes are distinguished by colour and selected genes are labelled. **(E)** STRING protein–protein interaction network of proximity-associated differentially expressed genes identified by donor-level pseudobulk analysis. Node colour denotes $\log_2$ fold change, node size $-\log_{10}$(FDR) and edge thickness STRING interaction confidence. Data are presented as mean ± SD with individual donors; P-values computed using the Mann–Whitney U test (C). Differential expression in A, B and D used DESeq2 with Benjamini–Hochberg FDR correction; cell-level estimates in A and B were obtained using Mann-Whitney U tests with Benjamini–Hochberg FDR correction.

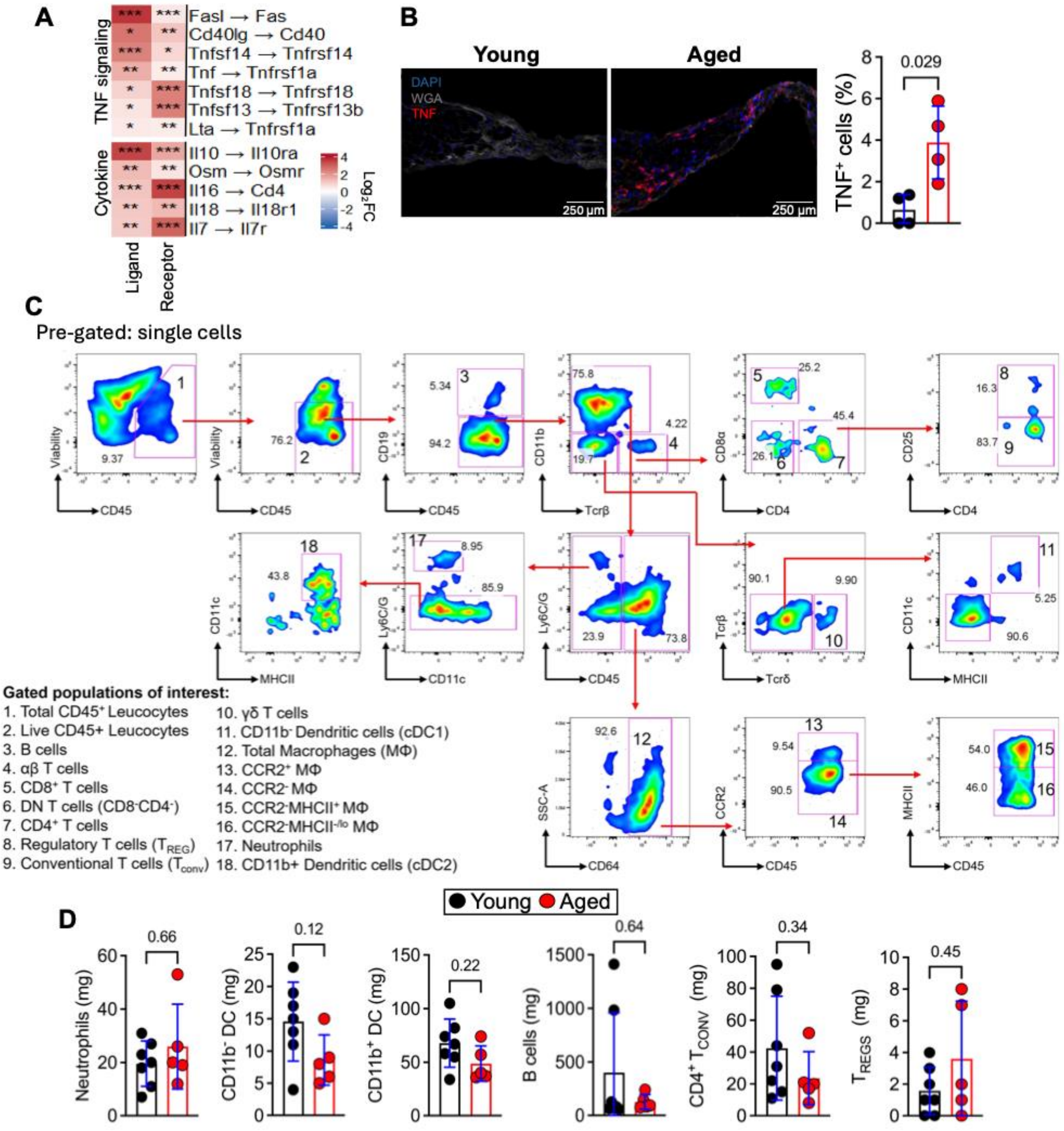

A
TNF signaling
Cytokine
Fasl → Fas
Cd40lg → Cd40
Tnfsf14 → Tnfrsf14
Tnf → Tnfrsf1a
Tnfsf18 → Tnfrsf18
Tnfsf13 → Tnfrsf13b
Lta → Tnfrsf1a
Il10 → Il10ra
Osm → Osmr
Il16 → Cd4
Il18 → Il18r1
Il7 → Il7r
Ligand
Receptor
Log2FC
B
Young
Aged
DAPI
WGA
TNF
250 µm
TNF+ cells (%)
0.029
C
Pre-gated: single cells
Viability
CD45
CD19
CD11b
Tcrβ
CD8α
CD4
CD25
CD11c
MHCII
Ly6C/G
Tcrδ
SSC-A
CD64
CCR2
Gated populations of interest:
1. Total CD45+ Leucocytes
2. Live CD45+ Leucocytes
3. B cells
4. αβ T cells
5. CD8+ T cells
6. DN T cells (CD8-CD4-)
7. CD4+ T cells
8. Regulatory T cells (TREG)
9. Conventional T cells (Tconv)
10. γδ T cells
11. CD11b- Dendritic cells (cDC1)
12. Total Macrophages (MΦ)
13. CCR2+ MΦ
14. CCR2- MΦ
15. CCR2-MHCII+ MΦ
16. CCR2-MHCII-/lo MΦ
17. Neutrophils
18. CD11b+ Dendritic cells (cDC2)
D
Young
Aged
Neutrophils (mg)
0.66
CD11b- DC (mg)
0.12
CD11b+ DC (mg)
0.22
B cells (mg)
0.64
CD4+ TCONV (mg)
0.34
TREGS (mg)
0.45

**Supplemental Figure 5. Immune cell remodelling and heightened pro-inflammatory ligand–receptor signalling in the aged murine SAN. (A)** Ligand–receptor interaction analysis demonstrating age-associated alterations in cytokine and TNF-family signalling pathways within the SAN. Heatmap colour represents $log_2FC$ of ligand and receptor expression in aged relative to young SAN tissue. Asterisks denote differential expression following Benjamini–Hochberg FDR correction ($^*P < 0.05$, $^{**}P < 0.005$, $^{***}P < 0.0005$). **(B)** Representative immunofluorescence images acquired at 20x showing TNF (red), WGA (grey), and DAPI (blue). Scale bars, 250 µm. Quantification of the percentage of $TNF^+$ cells in young (N=4) and aged (N=4) mice. **(C)** Representative flow cytometry gating strategy for identification of murine SAN leukocyte populations. Pseudocolour plots show sequential gating of single-cell suspensions, with arrows indicating the gating hierarchy. Values denote the percentage of cells within the parent population. **(D)** Quantification of neutrophils, $CD11b^+$ dendritic cells, $CD11b^-$ dendritic cells, B cells, $CD4^+$ T cells, and regulatory T cells in SAN tissue from young (N=7) and aged (N=5) mice. Data are presented as mean ± SD with individual biological replicates. P-values computed by Mann–Whitney U test (B, D).

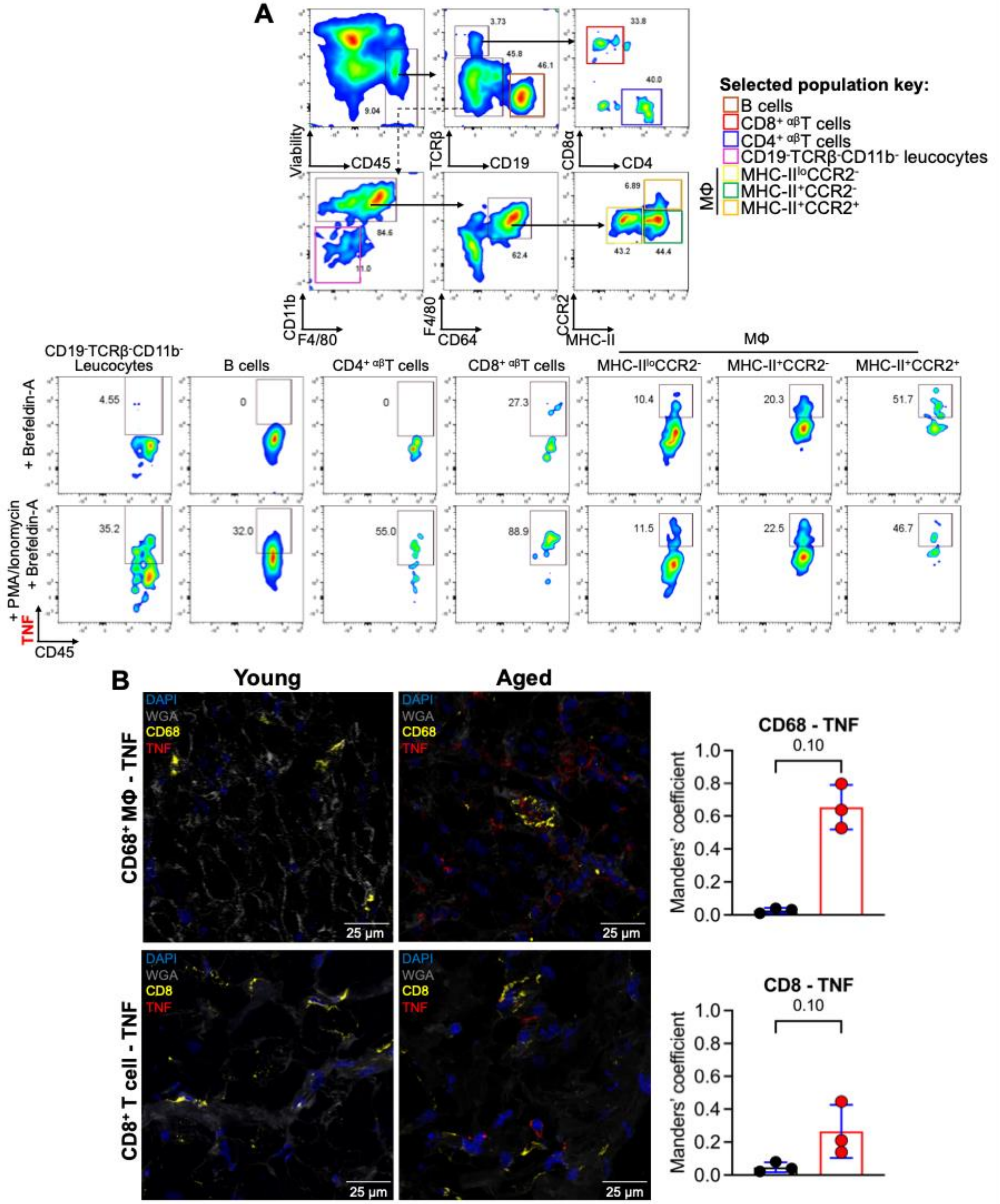
A
Viability
CD45
TCRβ
CD19
CD8α
CD4
CD11b
F4/80
CD64
CCR2
MHC-II
Selected population key:
B cells
CD8+ αβT cells
CD4+ αβT cells
CD19-TCRβ-CD11b- leucocytes
MHC-IIloCCR2-
MHC-II+CCR2-
MHC-II+CCR2+
MΦ
CD19-TCRβ-CD11b- Leucocytes
B cells
CD4+ αβT cells
CD8+ αβT cells
MHC-IIloCCR2-
MHC-II+CCR2-
MHC-II+CCR2+
+ Brefeldin-A
+ PMA/Ionomycin + Brefeldin-A
TNF
CD45
B
Young
Aged
CD68+ MΦ - TNF
CD8+ T cell - TNF
DAPI
WGA
CD68
CD8
TNF
25 µm
CD68 - TNF
CD8 - TNF
0.10
Manders' coefficient

**New Supplementary Figure 6. Cellular sources of TNF in the murine sinoatrial node. (A)** Flow cytometric analysis of TNF production by leucocyte populations isolated from murine SAN tissue from two mice (7 weeks old; N = 1 pooled biological sample). Representative gating shows identification of live $CD45^+$ leucocytes and the parent populations analysed, with TNF production under unstimulated conditions (Brefeldin A alone) and following stimulation with phorbol 12-myristate 13-acetate (PMA) and ionomycin in the presence of Brefeldin A. MΦ, macrophages. **(B)** Representative immunofluorescence images of murine SAN tissue from young (N = 3) and aged (N = 3) mice showing CD68 (yellow, upper panels) or CD8 (yellow, lower panels), TNF (red), WGA (grey) and nuclei (DAPI, blue). Colocalisation of TNF with $CD68^+$ macrophages and $CD8^+$ T cells was quantified using Manders' overlap coefficient (M2), representing the proportion of TNF fluorescence colocalising with CD68 or CD8. Scale bars, 25 μm. Data are presented as mean ± SD with individual biological replicates. P-Values computed by Mann–Whitney U test (B).

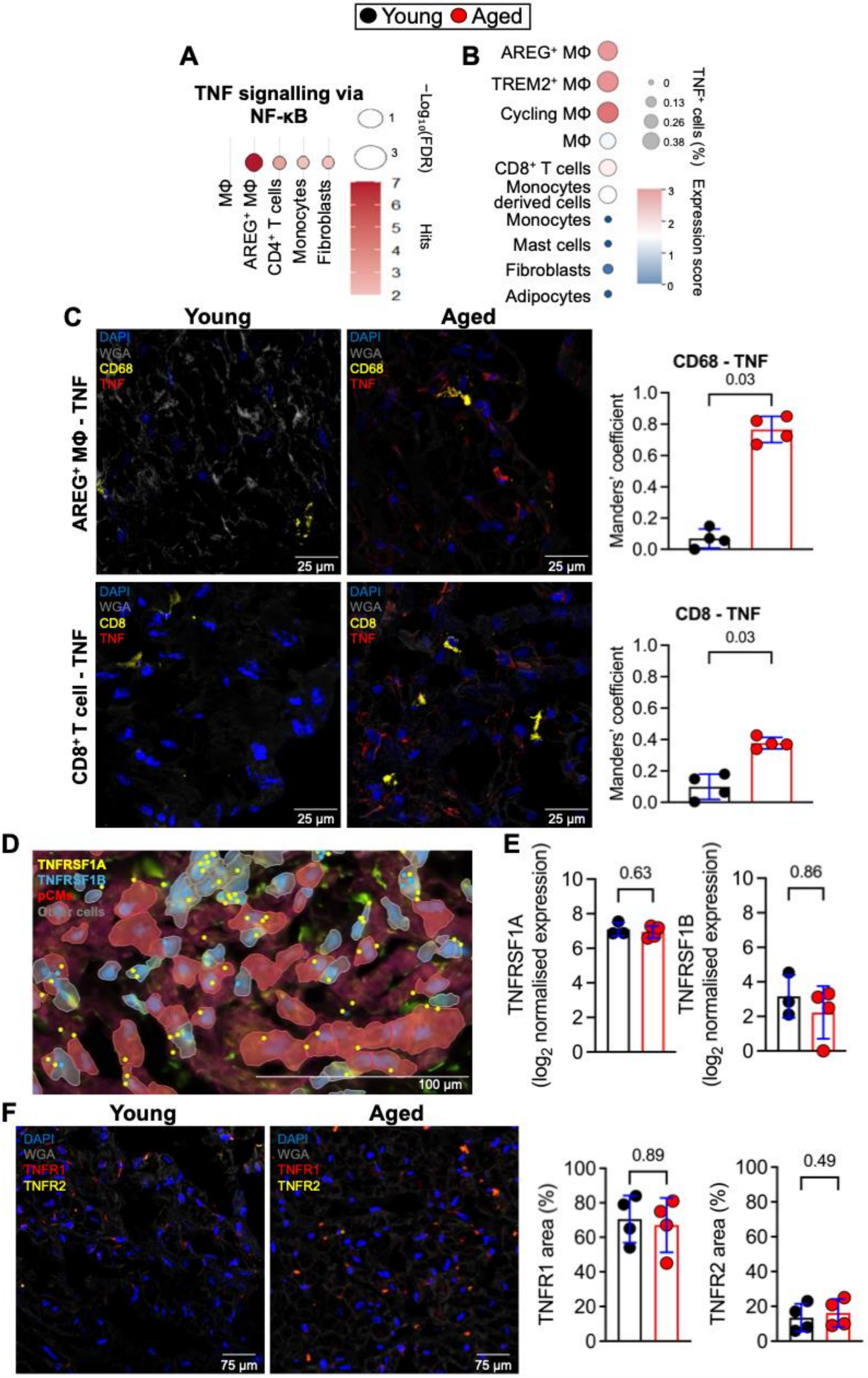

Young Aged
A
TNF signalling via NF-κB
−Log10(FDR)
1
3
Hits
7
6
5
4
3
2
MΦ
AREG+ MΦ
CD4+ T cells
Monocytes
Fibroblasts
B
AREG+ MΦ
TREM2+ MΦ
Cycling MΦ
MΦ
CD8+ T cells
Monocytes derived cells
Monocytes
Mast cells
Fibroblasts
Adipocytes
TNF+ cells (%)
0
0.13
0.26
0.38
Expression score
3
2
1
0
C
Young
Aged
AREG+ MΦ - TNF
DAPI
WGA
CD68
TNF
25 μm
CD68 - TNF
0.03
Manders' coefficient
1.0
0.8
0.6
0.4
0.2
0.0
CD8+ T cell - TNF
DAPI
WGA
CD8
TNF
25 μm
CD8 - TNF
0.03
Manders' coefficient
D
TNFRSF1A
TNFRSF1B
pCMs
Other cells
100 μm
E
0.63
0.86
TNFRSF1A (log2 normalised expression)
TNFRSF1B (log2 normalised expression)
10
8
6
4
2
0
F
Young
Aged
DAPI
WGA
TNFR1
TNFR2
75 μm
0.89
0.49
TNFR1 area (%)
TNFR2 area (%)
100
80
60
40
20
0

**New Supplemental Figure 7. TNF sources and TNF receptors in the human SAN. (A)** Gene set enrichment analysis of TNF signalling via NF-κB across immune cell populations in aged relative to young human SAN, using the Gene Ontology and Hallmark pathway databases. Dot colour denotes the number of genes (hits) associated with each gene set and dot size represents $-\log_{10}$(FDR). **(B)** Dot plot showing TNF expression across cell populations identified by Xenium spatial transcriptomics in aged (N=4) human SAN tissue. Dot size indicates the percentage of TNF-positive cells within each cell population and dot colour denotes mean TNF transcript counts. **(C)** Representative immunofluorescence images of human SAN tissue from young (N=4) and aged (N=4) donors showing CD68 (yellow, upper panels) or CD8 (yellow, lower panels), TNF (red), wheat germ agglutinin (WGA, grey) and nuclei (DAPI, blue). Colocalisation of TNF with $CD68^{+}$ macrophages and $CD8^{+}$ T cells was quantified using Manders' overlap coefficient (M2), representing the proportion of TNF fluorescence colocalising with CD68 or CD8. Scale bars, 25 µm. **(D)** Representative Xenium spatial transcriptomic map showing pacemaker cardiomyocytes (pCMs, red), TNFRSF1A expression (yellow), TNFRSF1B expression (blue) and other cell populations (grey) in human SAN tissue. Scale bar, 100 µm. **(E)** Donor-level expression of TNFRSF1A and TNFRSF1B in pacemaker cardiomyocytes from young (N=3) and aged (N=4) human SAN donors. **(F)** Representative immunofluorescence images of human SAN tissue from young (N=4) and aged (N=4) donors showing TNFR1 (red), TNFR2 (yellow), WGA (grey) and nuclei (DAPI, blue). Quantification of TNFR1- and TNFR2-positive area is shown alongside. Scale bar, 75 µm. Data are presented as mean ± SD with individual donors. P-values were computed using the Mann–Whitney U test (C, E, F).

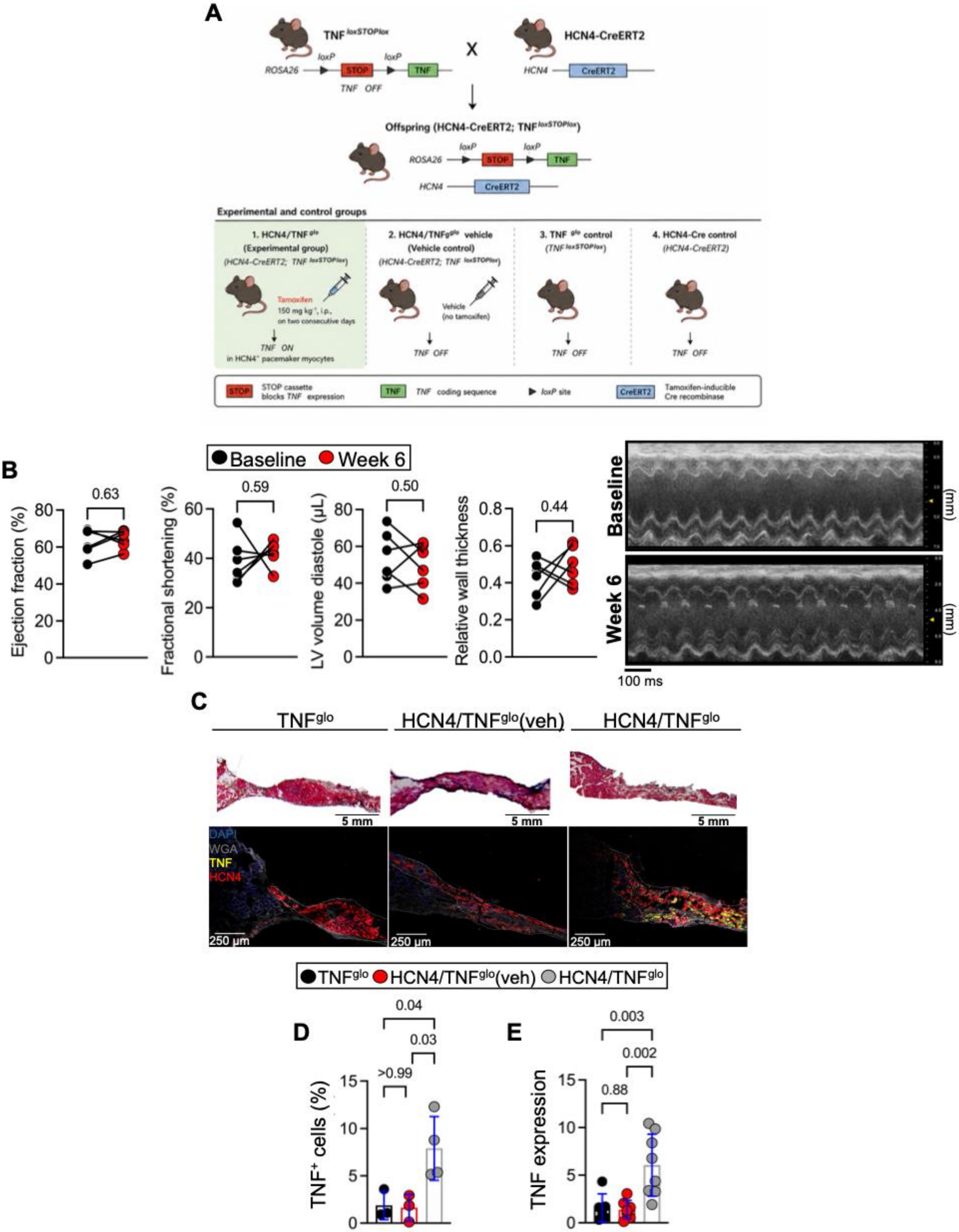
A
TNF loxSTOPlox
HCN4-CreERT2
Offspring (HCN4-CreERT2; TNF loxSTOPlox)
Experimental and control groups
1. HCN4/TNF glo (Experimental group)
2. HCN4/TNFglo vehicle (Vehicle control)
3. TNF glo control
4. HCN4-Cre control
TNF ON
TNF OFF
B
Baseline
Week 6
Ejection fraction (%)
Fractional shortening (%)
LV volume diastole (μL)
Relative wall thickness
0.63
0.59
0.50
0.44
100 ms
C
TNFglo
HCN4/TNFglo(veh)
HCN4/TNFglo
5 mm
250 μm
DAPI
WGA
TNF
HCN4
D
E
TNF+ cells (%)
TNF expression
0.04
0.03
>0.99
0.003
0.002
0.88

**Supplemental Figure 8. Validation of HCN4/TNF$^{glo}$ mouse model and preservatioin of ventricular systolic function. (A)** Schematic of the HCN4/TNF$^{glo}$ mouse model. HCN4-CreERT2 mice were crossed with TNF$^{loxSTOPlox}$ mice to generate HCN4/TNF$^{glo}$ offspring. Tamoxifen induced Cre-mediated excision of the loxP-flanked STOP cassette, resulting in selective TNF expression in HCN4$^{+}$ pacemaker myocytes. Experimental and control groups used throughout the study shown. **(B)** Quantification of ejection fraction, fractional shortening, relative wall thickness and left ventricular volume at baseline and 6 weeks post tamoxifen (N=6). Representative parasternal short-axis echocardiographic images from a single HCN4 TNF$^{glo}$ mouse at baseline and 6 weeks post tamoxifen injection. **(C)** Representative Masson's trichrome staining and immunofluorescence images acquired at 20x of SAN tissue from TNF$^{glo}$ controls, vehicle-injected HCN4/TNF$^{glo}$ mice, and tamoxifen-injected HCN4/TNF$^{glo}$ mice. Immunofluorescence images show TNF (yellow), HCN4 (red), WGA (grey), and DAPI (blue). Scale bars, 5 mm and 250 µm. **(D)** Quantification of the percentage of TNF$^{+}$ cells in SAN tissue from TNF$^{glo}$ controls (N=3), vehicle-injected HCN4/TNF$^{glo}$ mice (N=3), and tamoxifen-injected HCN4/TNF$^{glo}$ mice (N=4). **(E)** Relative TNF mRNA expression in SAN tissue from TNF$^{glo}$ controls (N=8), vehicle-injected HCN4/TNF$^{glo}$ mice (N=8), and tamoxifen-injected HCN4/TNF$^{glo}$ mice (N=8). Gene expression was normalised to IPO8 and calculated using the $2^{-\Delta\Delta Ct}$ method. Data are presented as mean ± SD with individual biological replicates. P-values computed by paired Students t-test (B) or one-way ANOVA with Šídák's multiple-comparisons test (D, E).

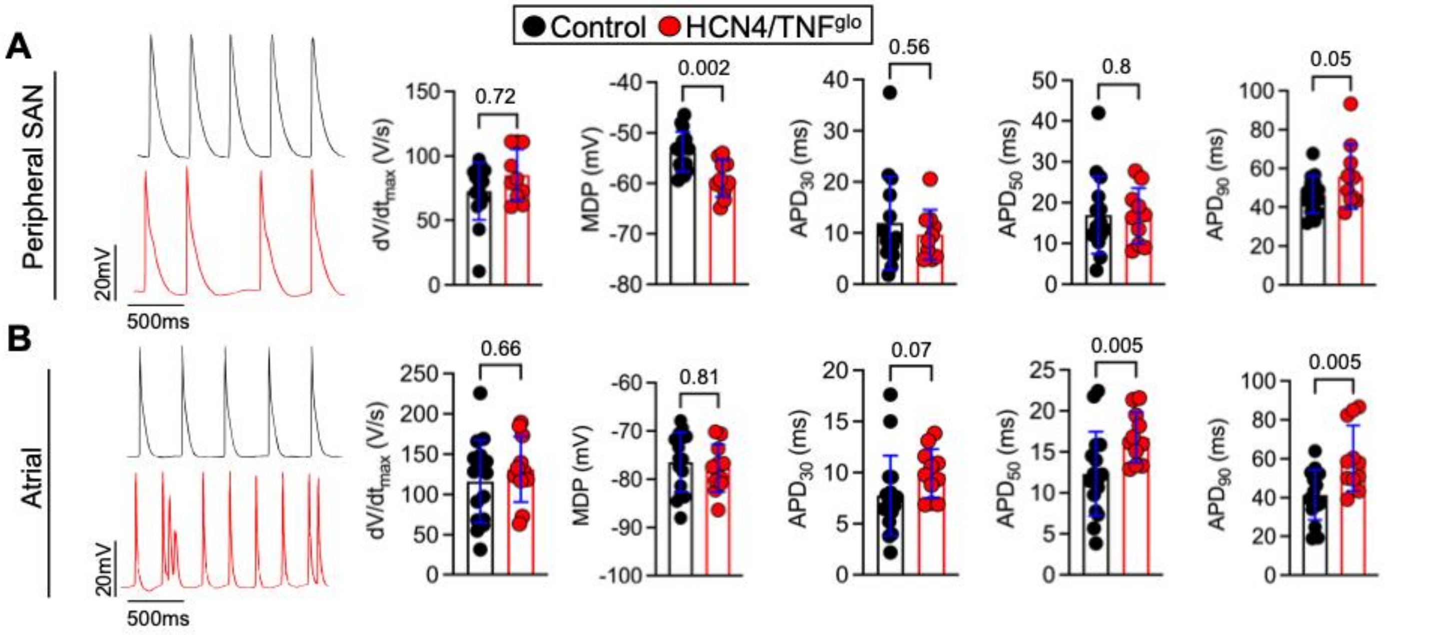


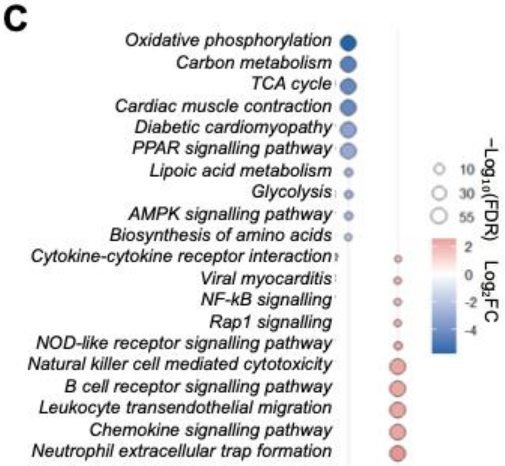


E

| Protein | Control | TNF | FC | FDR |
|---|---|---|---|---|
| TNF | 26.33 ± 31.55 | 913.25 ± 258.15 | ↑ 34.7 | 0.02857 |
| HCN4 | 639.10 ± 182.65 | 127.08 ± 52.64 | ↓ 5.2 | 0.0000331 |
| HCN1 | 542.23 ± 182.13 | 112.13 ± 48.06 | ↓ 4.9 | 0.000421 |
| CD68 | 135.05 ± 25.22 | 498.88 ± 137.46 | ↑ 3.7 | 0.02857 |
| CD4 | 112.35 ± 10.88 | 757.13 ± 276.15 | ↑ 6.4 | 0.000042 |
| NF-κB1 | 3816.80 ± 502.86 | 8051.83 ± 1147.86 | ↑ 2.1 | 0.000578 |
| $Ca_V1.2$ | 558.30 ± 137.96 | 413.15 ± 43.58 | 0 | 0.34286 |
| Cavα2δ2 | 1246.35 ± 385.40 | 719.05 ± 76.45 | ↓ 1.7 | 0.024966 |
| CCL6 | 450.25 ± 293.57 | 4332.50 ± 2395.15 | ↑ 9.6 | 0.000179 |
| CCL9 | 802.75 ± 396.17 | 4893.75 ± 2486.70 | ↑ 6.1 | 0.02857 |
| IL-16 | 7073.25 ± 1120.39 | 22228.50 ± 6220.32 | ↑ 3.1 | 0.000465 |
| IL-33 | 487.20 ± 86.61 | 859.68 ± 73.18 | ↑ 1.8 | 0.0003426 |

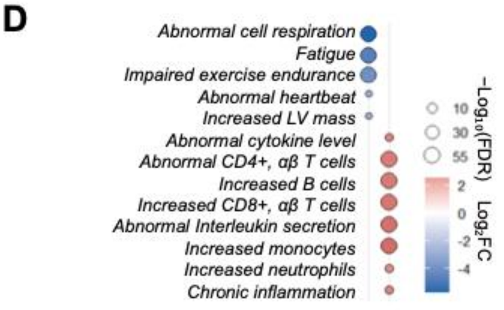


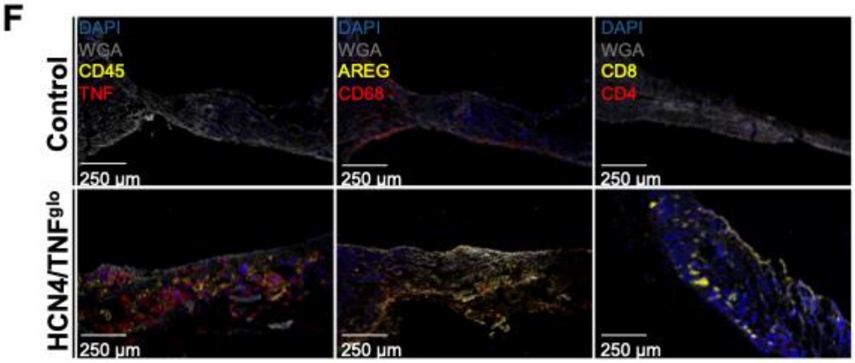


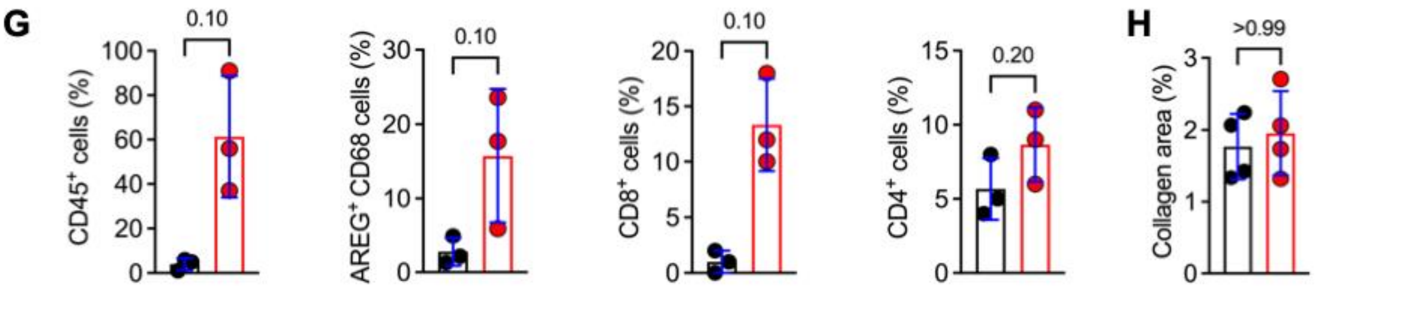

**Supplemental Figure 9. Electrophysiological, proteomic and immune remodelling in HCN4/TNF$^{glo}$ mice. (A, B)** Representative sharp microelectrode action potential recordings from **(A)** SAN peripheral (SAN-p) cells and **(B)** atrial cardiomyocytes in control (N=15) and HCN4/TNF$^{glo}$ (N=10) mice. Quantification of maximum upstroke velocity ($dV/dt_{max}$), maximum diastolic potential (MDP), and action potential duration at 30%, 50%, and 90% repolarisation ($APD_{30}$, $APD_{50}$, $APD_{90}$). Each dot represents the mean value obtained from averaged impalements (control, (A) n=48, (B) n=50; HCN4/TNF$^{glo}$, (A) n=49, (B) n=44) **(C)** GSEA in KEGG database and **(D)** mammalian phenotype databases based on differentially protein abundance in SAN tissue from control (N=4) and HCN4/TNF$^{glo}$ (N=4) mice. Dot colour denotes $\log_2$FC and dot size represents $-\log_{10}$(FDR). **(E)** Table showing normalised protein abundance (mean ± SD), fold-change (FC), and false discovery rate (FDR) for selected proteins in SAN tissue from control (N=4) and HCN4/TNF$^{glo}$ (N=4) mice. **(F)** Representative immunofluorescence images acquired at 20x of SAN tissue from control and HCN4/TNF$^{glo}$ mice showing CD45 (yellow) and TNF (red), AREG (yellow) and CD68 (red), or CD8 (yellow) and CD4 (red), together with WGA (grey) and DAPI (blue). Scale bars, 250 μm. **(G)** Quantification of the percentage of CD45$^{+}$ leukocytes, AREG$^{+}$ CD68$^{+}$ macrophages, CD8$^{+}$ T cells, CD4$^{+}$ T cells from control (N=3) and HCN4/TNF$^{glo}$ (N=3). **(H)** Quantification of collagen area fraction in SAN tissue from vehicle-injected HCN4/TNF$^{glo}$ mice (N=4), and tamoxifen-injected HCN4/TNF$^{glo}$ mice (N=4). Data are presented as mean ± SD with individual biological replicates. P-values computed by nested t-test (A, B), permutation testing with Benjamini–Hochberg correction (C, D), limma with Benjamini–Hochberg correction (E), or Mann-Whitney U test (G, H).

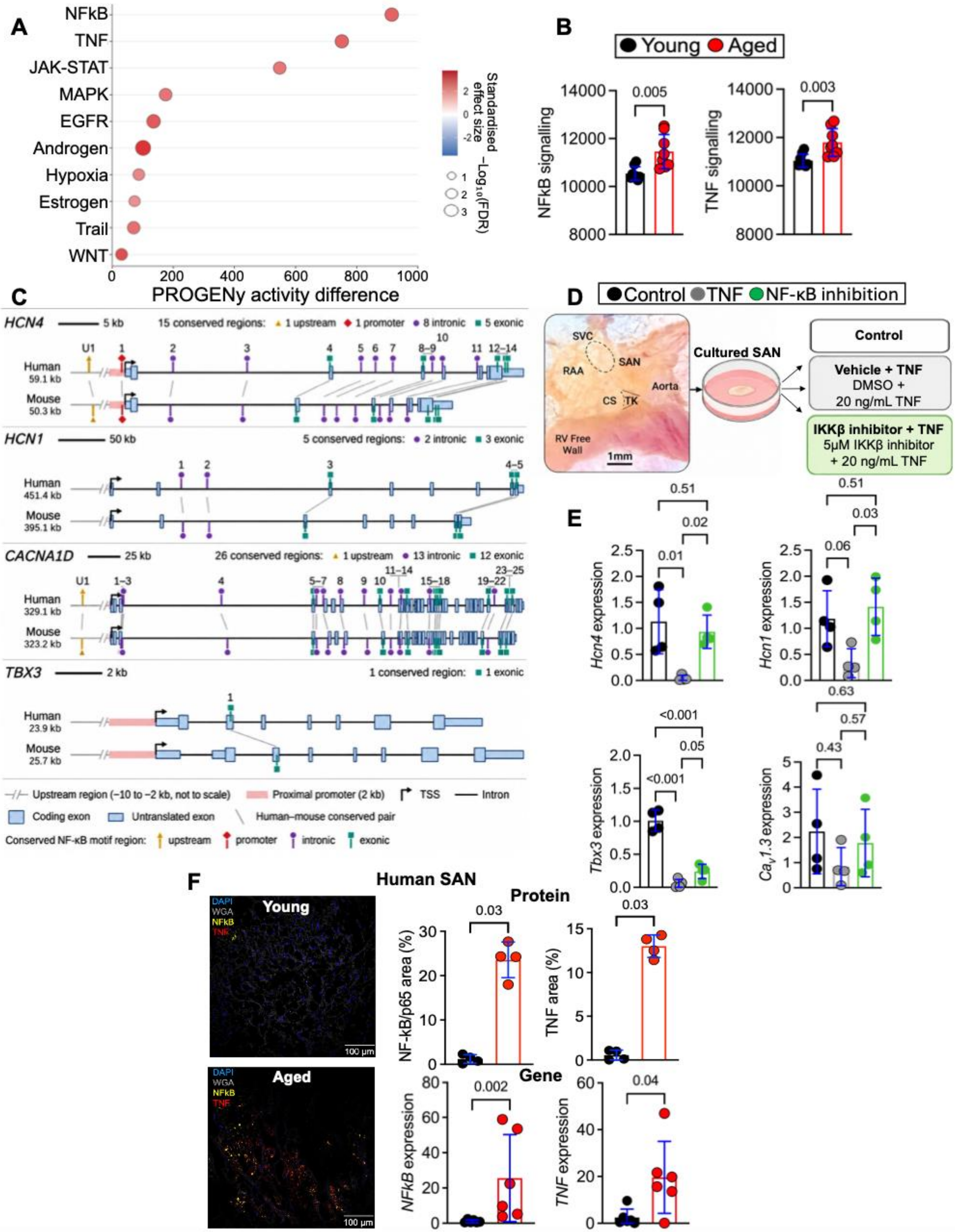
A
NFkB
TNF
JAK-STAT
MAPK
EGFR
Androgen
Hypoxia
Estrogen
Trail
WNT
0
200
400
600
800
1000
PROGENy activity difference
Standardised effect size
2
0
-2
−Log₁₀(FDR)
1
2
3
B
Young
Aged
NFkB signalling
TNF signalling
14000
12000
10000
8000
0.005
0.003
C
HCN4
5 kb
15 conserved regions:
1 upstream
1 promoter
8 intronic
5 exonic
Human
59.1 kb
Mouse
50.3 kb
HCN1
50 kb
5 conserved regions:
2 intronic
3 exonic
Human
451.4 kb
Mouse
395.1 kb
CACNA1D
25 kb
26 conserved regions:
1 upstream
13 intronic
12 exonic
Human
329.1 kb
Mouse
323.2 kb
TBX3
2 kb
1 conserved region:
1 exonic
Human
23.9 kb
Mouse
25.7 kb
Upstream region (−10 to −2 kb, not to scale)
Proximal promoter (2 kb)
TSS
Intron
Coding exon
Untranslated exon
Human–mouse conserved pair
Conserved NF-κB motif region:
upstream
promoter
intronic
exonic
D
Control
TNF
NF-κB inhibition
SVC
SAN
RAA
Aorta
CS
TK
RV Free Wall
1mm
Cultured SAN
Control
Vehicle + TNF
DMSO +
20 ng/mL TNF
IKKβ inhibitor + TNF
5μM IKKβ inhibitor
+ 20 ng/mL TNF
E
Hcn4 expression
0.51
0.02
0.01
Hcn1 expression
0.51
0.03
0.06
Tbx3 expression
<0.001
0.05
<0.001
Cav1.3 expression
0.63
0.57
0.43
F
Human SAN
Young
Aged
DAPI
WGA
NFkB
TNF
100 μm
Protein
NF-kB/p65 area (%)
0.03
TNF area (%)
0.03
Gene
NFkB expression
0.002
TNF expression
0.04

**New Supplemental Figure 10. TNF/NF-κB signalling in the aged SAN and the effect of IKKβ inhibition on pacemaker channel expression. (A)** Differential PROGENy signalling pathway activity in aged (N=8) relative to young (N=8) mouse SAN tissue. The x-axis represents the difference in inferred pathway activity (aged − young), with positive values indicating higher predicted activity in aged SAN. Point size is proportional to $-\log_{10}$(FDR) and colour denotes the standardised effect size. **(B)** PROGENy activity estimates for NF-κB and TNF signalling in young (N=8) and aged (N=8) mouse SAN tissue. Individual points represent biological replicates. **(C)** Conserved predicted NF-κB binding motifs at the human (hg38) and mouse (mm39) HCN4, HCN1, CACNA1D and TBX3 loci. For each gene, 10 kb upstream of the transcription start site (TSS) and the full transcript were scanned with TRANSFAC NF-κB binding matrices, and overlapping predictions conserved between species were merged into distinct regions. Schematics show the upstream region (−10 to −2 kb, fixed width, not to scale), the proximal promoter (2 kb), coding and untranslated exons, and introns, with arrows marking the TSS. Markers denote conserved regions classified as upstream, promoter, intronic or exonic, and lines connect corresponding regions in human and mouse. Gene models are from Ensembl release 116. **(D)** Schematic of the ex vivo mouse SAN culture and treatment protocol. Isolated preparations were pre-incubated for 12 h with the IKKβ inhibitor BMS-345541 (5 µM) or vehicle (0.05% DMSO), and preparations were then exposed to recombinant mouse TNF (20 ng/mL) for 36 h. Control preparations were cultured in Medium 199 for 48 h. **(E)** Relative mRNA expression of *Hcn4, Hcn1, Cacna1d* and *Tbx3* in control (N = 4), TNF-exposed preparations treated with vehicle (N=4) or IKKβ inhibitor (N=4), normalised to Ipo8 and calculated using the $2^{-\Delta\Delta Ct}$ method. **(F)** Representative immunofluorescence images of human SAN tissue from young (N=4) and aged (N=4) donors showing NF-κB p65 (yellow), TNF (red), WGA (grey) and nuclei (DAPI, blue). Quantification of the percentage of tissue area positive for NF-κB p65 and TNF immunoreactivity in young and aged SAN tissue is shown alongside. Scale bars, 100 µm. Relative mRNA expression of *NFKB1* and *TNF* in human SAN tissue from young (N=6) and aged (N=6) donors, normalised to GAPDH and calculated using the $2^{-\Delta\Delta Ct}$ method. Data are presented as mean ± SD with individual biological replicates. P-values were computed using the Wilcoxon rank-sum test with Benjamini–Hochberg correction (A, B), one-way ANOVA with Šídák's multiple-comparisons test (E) or the Mann–Whitney U test (F).

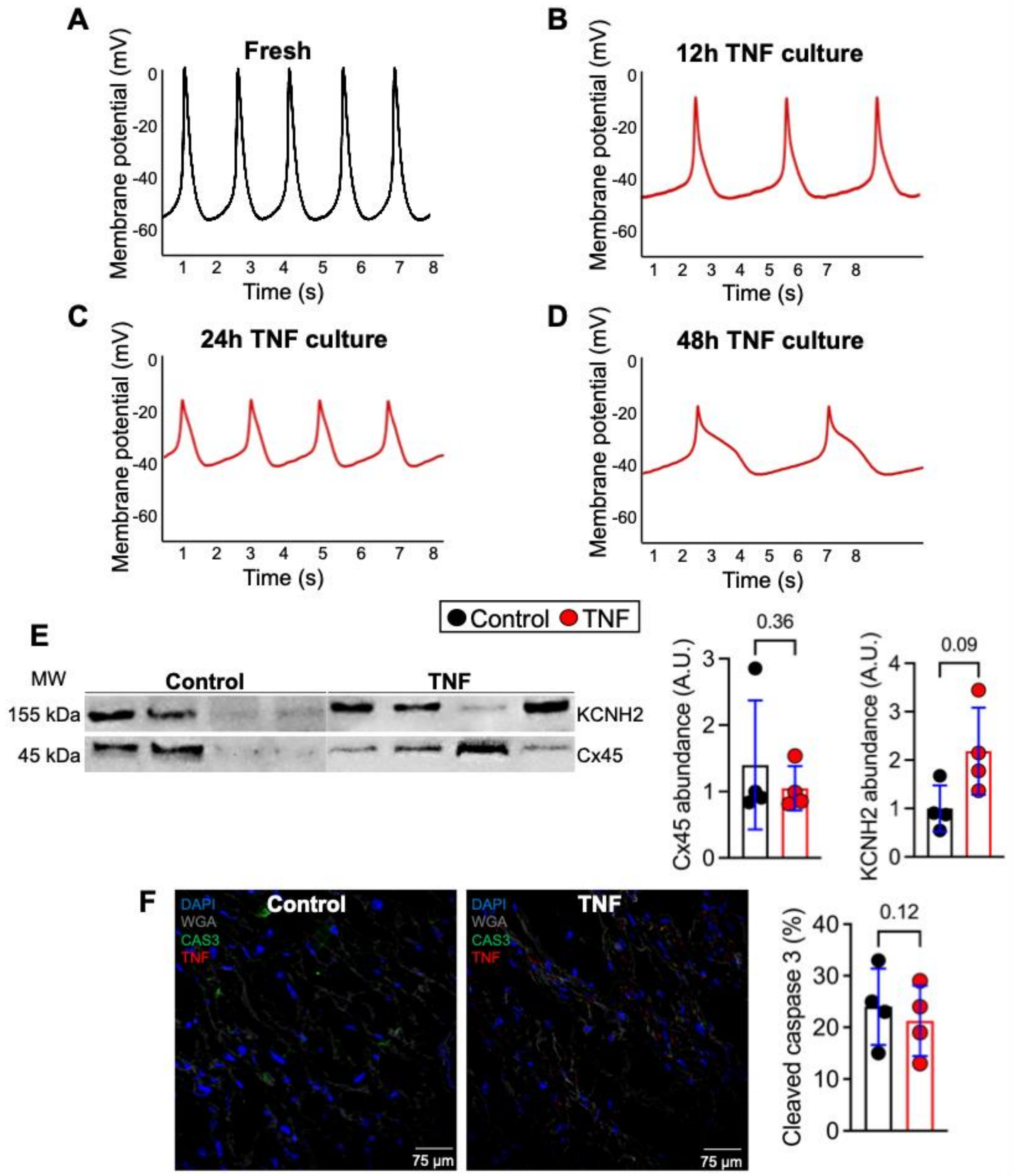
A
Fresh
Membrane potential (mV)
0
-20
-40
-60
1 2 3 4 5 6 7 8
Time (s)
B
12h TNF culture
Membrane potential (mV)
Time (s)
C
24h TNF culture
Membrane potential (mV)
Time (s)
D
48h TNF culture
Membrane potential (mV)
Time (s)
Control TNF
E
MW
Control
TNF
155 kDa
45 kDa
KCNH2
Cx45
Cx45 abundance (A.U.)
0.36
KCNH2 abundance (A.U.)
0.09
F
DAPI
WGA
CAS3
TNF
Control
TNF
75 µm
75 µm
Cleaved caspase 3 (%)
0.12

**Supplemental Figure 11. Human SAN slices remain spontaneously active during organotypic culture and show TNF-dependent molecular remodelling. (A–D)** Representative spontaneous action potential recordings from a single human SAN donor obtained **(A)** fresh and after **(B)** 12 h, **(C)** 24 h and **(D)** 48 h of organotypic culture in the presence of TNF. These recordings demonstrate persistence of spontaneous electrical activity throughout the culture period. Changes in action potential morphology over time were not used to infer TNF-specific electrophysiological remodelling because equivalent time-matched control recordings were not available at each interval. **(E)** Representative western blots and quantification of stain-free total protein-normalised Cx45 and KCNH2 protein abundance in control (N=4) and TNF-treated (N=4) human SAN slices after 48 h. **(F)** Representative immunofluorescence images of paired human SAN slices cultured for 48 h under control conditions or with TNF, showing cleaved caspase-3 (CAS3, green), TNF (red), WGA (grey) and nuclei (DAPI, blue). Quantification of $CAS3^+$ cells in paired control and TNF-treated slices from four donors is shown alongside. Scale bars, 75 µm. Data are presented as mean ± SD with individual donors. P-values computed by paired Student's t-test (E, F).

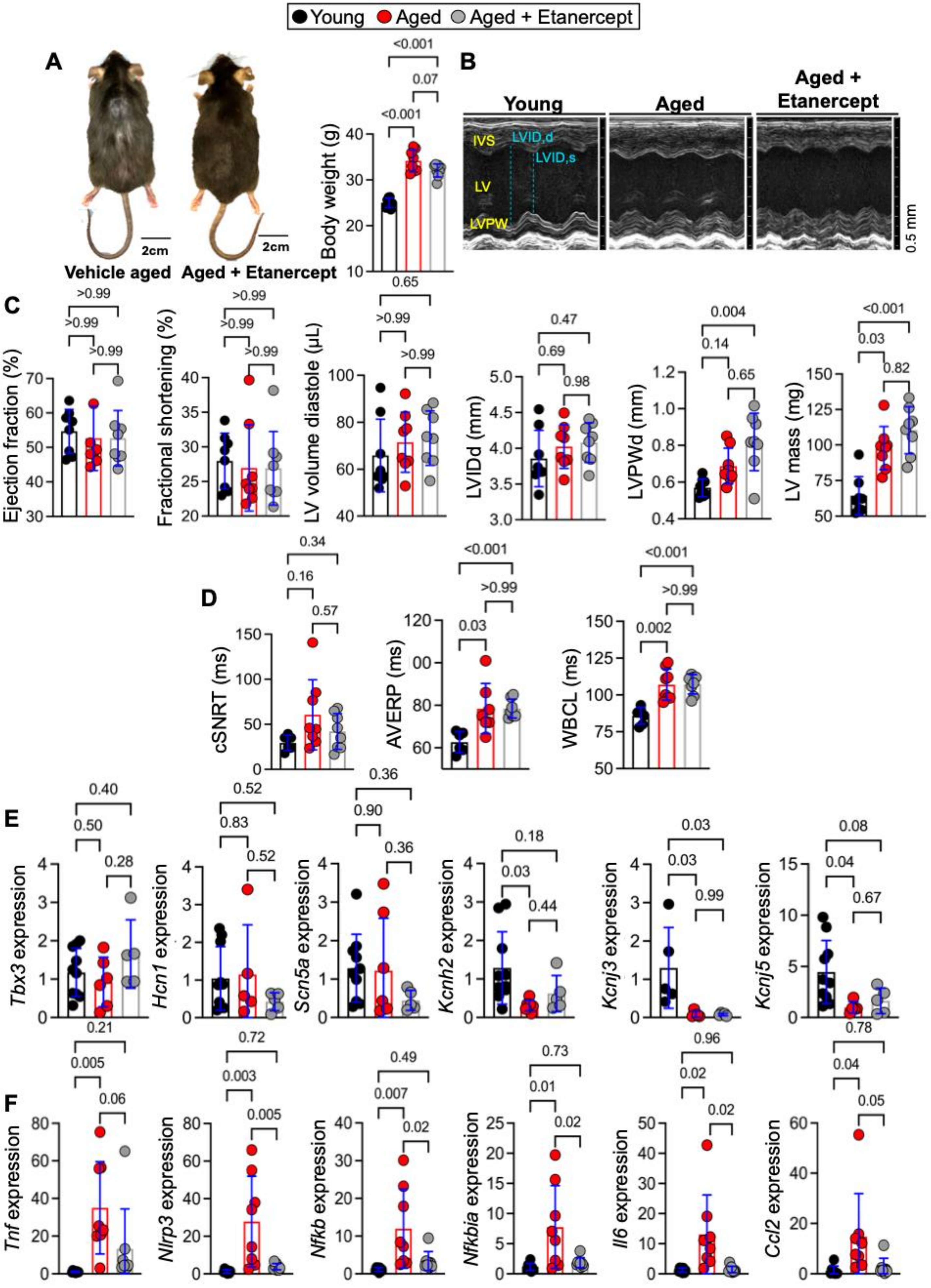

Young
Aged
Aged + Etanercept
A
2cm
Vehicle aged
Aged + Etanercept
Body weight (g)
B
IVS
LVID,d
LVID,s
LV
LVPW
0.5 mm
C
Ejection fraction (%)
Fractional shortening (%)
LV volume diastole (μL)
LVIDd (mm)
LVPWd (mm)
LV mass (mg)
D
cSNRT (ms)
AVERP (ms)
WBCL (ms)
E
Tbx3 expression
Hcn1 expression
Scn5a expression
Kcnh2 expression
Kcnj3 expression
Kcnj5 expression
F
Tnf expression
Nlrp3 expression
Nfkb expression
Nfkbia expression
Il6 expression
Ccl2 expression

**Supplemental Figure 12. Etanercept attenuates age-associated frailty and SAN remodelling. (A)** Representative images of aged mice treated with vehicle or etanercept. Vehicle-treated aged mice exhibited signs of frailty including alopecia, tumour burden, and reduced grooming, whereas etanercept-treated mice showed improved physical appearance. Scale bars, 2 cm. Body weight quantified in young (N=8), aged (N=8), and aged + etanercept mice (N=8). **(B)** Representative M-mode echocardiographic images from young, aged, and aged + etanercept mice. The young panel highlights anatomical landmarks including the interventricular septum (IVS), left ventricle (LV), and left ventricular posterior wall (LVPW), together with measurements of left ventricular internal diameter during diastole (LVID,d) and systole (LVID,s). **(C)** Quantification of ejection fraction, fractional shortening, left ventricular (LV) volume, LV internal diameter (LVID), LV posterior wall (LVPW) thickness, and LV mass in young (N=8), aged (N=8), and aged + etanercept mice (N=8). **(D)** Quantification of corrected sinoatrial node recovery time (cSNRT), atrioventricular effective refractory period (AVERP), and Wenckebach cycle length (WBCL) in Langendorff-perfused hearts from young (N=8), aged (N=8), and aged + etanercept mice (N=8). **(E)** Relative mRNA expression of pacemaker and ion channel transcripts in SAN tissue from young (N=8), aged (N=8), and aged + etanercept mice (N=8). **(F)** Relative mRNA expression of inflammatory transcripts in liver tissue from young (N=8), aged (N=8), and aged + etanercept mice (N=8). Gene expression was normalised to IPO8 and calculated using the $2^{-\Delta\Delta Ct}$ method. Data are presented as individual biological replicates with mean ± SD. P-values computed by one-way ANOVA with Šídák's multiple-comparisons test.

### References


1 Siwik, D. A., Chang, D. L. & Colucci, W. S. Interleukin-1beta and tumor necrosis factor-alpha decrease collagen synthesis and increase matrix metalloproteinase activity in cardiac fibroblasts in vitro. *Circ Res* **86**, 1259–1265 (2000). https://doi.org/10.1161/01.res.86.12.1259

2 Saraf, A. *et al.* Functional and molecular effects of TNF-alpha on human iPSC-derived cardiomyocytes. *Stem Cell Res* **52**, 102218 (2021). https://doi.org/10.1016/j.scr.2021.102218

3 Howlett, S. E., Rutenberg, A. D. & Rockwood, K. The degree of frailty as a translational measure of health in aging. *Nat Aging* **1**, 651–665 (2021). https://doi.org/10.1038/s43587-021-00099-3

4 Moghtadaei, M. *et al.* The impacts of age and frailty on heart rate and sinoatrial node function. *J Physiol* **594**, 7105–7126 (2016). https://doi.org/10.1113/JP272979

5 Whitehead, J. C. *et al.* A clinical frailty index in aging mice: comparisons with frailty index data in humans. *J Gerontol A Biol Sci Med Sci* **69**, 621–632 (2014). https://doi.org/10.1093/gerona/glt136

6 Sciorati, C. *et al.* Pharmacological blockade of TNFalpha prevents sarcopenia and prolongs survival in aging mice. *Aging (Albany NY)* **12**, 23497–23508 (2020). https://doi.org/10.18632/aging.202200

7 Patel, B. A. *et al.* The TNF-alpha antagonist etanercept reverses age-related decreases in colonic SERT expression and faecal output in mice. *Sci Rep* **7**, 42754 (2017). https://doi.org/10.1038/srep42754

8 Aschar-Sobbi, R. *et al.* Increased atrial arrhythmia susceptibility induced by intense endurance exercise in mice requires TNFalpha. *Nat Commun* **6**, 6018 (2015). https://doi.org/10.1038/ncomms7018

9 Busch, M. *et al.* Effects of systemic and intravitreal TNF-alpha inhibition in experimental autoimmune uveoretinitis. *Invest Ophthalmol Vis Sci* **54**, 39–46 (2013). https://doi.org/10.1167/iovs.12-10138

10 Yang, J. *et al.* Risk Assessment of Etanercept in Mice Chronically Infected With Toxoplasma gondii. *Front Microbiol* **9**, 2822 (2018). https://doi.org/10.3389/fmicb.2018.02822

11 Yi, H. *et al.* Induced production of anti-etanercept antibody in collagen-induced arthritis. *Mol Med Rep* **9**, 2301–2308 (2014). https://doi.org/10.3892/mmr.2014.2127

12 Yoshitaka, T. *et al.* Etanercept administration to neonatal SH3BP2 knock-in cherubism mice prevents TNF-alpha-induced inflammation and bone loss. *J Bone Miner Res* **29**, 1170–1182 (2014). https://doi.org/10.1002/jbmr.2125

13 Soattin, L. *et al.* Endurance exercise remodels pulmonary vein sleeve myocytes and promotes a proarrhythmic atrial substrate. *Eur Heart J* (2026). https://doi.org/10.1093/eurheartj/ehag358

14 D'Souza, A. *et al.* A circadian clock in the sinus node mediates day-night rhythms in Hcn4 and heart rate. *Heart Rhythm* **18**, 801–810 (2021). https://doi.org/10.1016/j.hrthm.2020.11.026

15 Hennis, K. *et al.* In vivo and ex vivo electrophysiological study of the mouse heart to characterize the cardiac conduction system, including atrial and ventricular vulnerability. *Nat Protoc* **17**, 1189–1222 (2022). https://doi.org/10.1038/s41596-021-00678-z

16 Mangoni, M. E. & Nargeot, J. Properties of the hyperpolarization-activated current (If) in isolated mouse sino-atrial cells. *Cardiovascular Research* **52**, 51–64 (2001). https://doi.org/10.1016/s0008-6363(01)00370-4

17 Fredriksson, L. *et al.* Diclofenac inhibits tumor necrosis factor-alpha-induced nuclear factor-kappaB activation causing synergistic hepatocyte apoptosis. *Hepatology* **53**, 2027–2041 (2011). https://doi.org/10.1002/hep.24314

18 Du, Z. *et al.* Inhibition of type I interferon-mediated antiviral action in human glioma cells by the IKK inhibitors BMS-345541 and TPCA-1. *J Interferon Cytokine Res* **32**, 368–377 (2012). https://doi.org/10.1089/jir.2012.0002
19 Ortis, F. *et al.* Differential usage of NF-kappaB activating signals by IL-1beta and TNF-alpha in pancreatic beta cells. *FEBS Lett* **586**, 984–989 (2012). https://doi.org/10.1016/j.febslet.2012.02.021
20 Loots, G. G. & Ovcharenko, I. rVISTA 2.0: evolutionary analysis of transcription factor binding sites. *Nucleic Acids Res* **32**, W217–221 (2004). https://doi.org/10.1093/nar/gkh383
21 Kotah, J. M. *et al.* Beyond the nuclear border: single-cell analysis of in situ sequenced human brain tissue using cellular features. *Commun Biol* **8**, 1089 (2025). https://doi.org/10.1038/s42003-025-08518-6
22 Palla, G. *et al.* Squidpy: a scalable framework for spatial omics analysis. *Nat Methods* **19**, 171–178 (2022). https://doi.org/10.1038/s41592-021-01358-2
23 Genomics, x. *Xenium Analyzer*, <https://www.10xgenomics.com/platforms/xenium?utm> (
24 Varrone, M., Tavernari, D., Santamaria-Martinez, A., Walsh, L. A. & Ciriello, G. CellCharter reveals spatial cell niches associated with tissue remodeling and cell plasticity. *Nat Genet* **56**, 74–84 (2024). https://doi.org/10.1038/s41588-023-01588-4
25 Hao, Y. *et al.* Integrated analysis of multimodal single-cell data. *Cell* **184**, 3573–3587 e3529 (2021). https://doi.org/10.1016/j.cell.2021.04.048
26 Wickham, H. *ggplot2: Elegant Graphics for Data Analysis*. (Springer-Verlag New York, 2016).
27 Mages, S. *et al.* TACCO unifies annotation transfer and decomposition of cell identities for single-cell and spatial omics. *Nat Biotechnol* **41**, 1465–1473 (2023). https://doi.org/10.1038/s41587-023-01657-3
28 Kanemaru, K. *et al.* Spatially resolved multiomics of human cardiac niches. *Nature* **619**, 801–810 (2023). https://doi.org/10.1038/s41586-023-06311-1
29 Katsuki, T. *et al.* Inhibition of Scarb1 on Endothelial Cells Attenuates Pressure Overload-Induced Heart Failure Progression. *JACC Basic Transl Sci* **10**, 101308 (2025). https://doi.org/10.1016/j.jacbts.2025.05.003
30 Li, X. *et al.* Ventricular assist device unloading reverses microvascular senescence in single ventricle disease. *Nature Cardiovascular Research* **5**, 262–280 (2026). https://doi.org/10.1038/s44161-026-00790-x
31 FNN: Fast Nearest Neighbor Search Algorithms and Applications v. 1.1.4.1 (2024).
32 Li, Y. *et al.* SOAR elucidates biological insights and empowers drug discovery through spatial transcriptomics. *Sci Adv* **11**, eadt7450 (2025). https://doi.org/10.1126/sciadv.adt7450
33 Kanehisa, M. & Goto, S. KEGG: kyoto encyclopedia of genes and genomes. *Nucleic Acids Res* **28**, 27–30 (2000). https://doi.org/10.1093/nar/28.1.27
34 Ramilowski, J. A. *et al.* A draft network of ligand-receptor-mediated multicellular signalling in human. *Nat Commun* **6**, 7866 (2015). https://doi.org/10.1038/ncomms8866
35 Schubert, M. *et al.* Perturbation-response genes reveal signaling footprints in cancer gene expression. *Nat Commun* **9**, 20 (2018). https://doi.org/10.1038/s41467-017-02391-6
36 Kahnert, K. *et al.* Proteomics couples electrical remodelling to inflammation in a murine model of heart failure with sinus node dysfunction. *Cardiovasc Res* **120**, 927–942 (2024). https://doi.org/10.1093/cvr/cvae054
37 Linscheid, N. *et al.* Quantitative proteome comparison of human hearts with those of model organisms. *PLoS Biol* **19**, e3001144 (2021). https://doi.org/10.1371/journal.pbio.3001144

38 Ritchie, M. E. *et al.* limma powers differential expression analyses for RNA-sequencing and microarray studies. *Nucleic Acids Res* **43**, e47 (2015). https://doi.org/10.1093/nar/gkv007
39 Yu, G., Wang, L. G., Han, Y. & He, Q. Y. clusterProfiler: an R package for comparing biological themes among gene clusters. *OMICS* **16**, 284–287 (2012). https://doi.org/10.1089/omi.2011.0118
40 Yu, G., Wang, L. G., Yan, G. R. & He, Q. Y. DOSE: an R/Bioconductor package for disease ontology semantic and enrichment analysis. *Bioinformatics* **31**, 608–609 (2015). https://doi.org/10.1093/bioinformatics/btu684
41 Yu, G. & He, Q. Y. ReactomePA: an R/Bioconductor package for reactome pathway analysis and visualization. *Mol Biosyst* **12**, 477–479 (2016). https://doi.org/10.1039/c5mb00663e
42 Tikhomirov, R. *et al.* Cardiac GR Mediates the Diurnal Rhythm in Ventricular Arrhythmia Susceptibility. *Circ Res* **134**, 1306–1326 (2024). https://doi.org/10.1161/CIRCRESAHA.123.323464
43 Tsutsui, K. *et al.* A coupled-clock system drives the automaticity of human sinoatrial nodal pacemaker cells. *Sci Signal* **11** (2018). https://doi.org/10.1126/scisignal.aap7608
44 Verkerk, A. O. *et al.* Pacemaker current (I(f)) in the human sinoatrial node. *Eur Heart J* **28**, 2472–2478 (2007). https://doi.org/10.1093/eurheartj/ehm339
45 Drouin, E. Electrophysiologic properties of the adult human sinus node. *J Cardiovasc Electrophysiol* **8**, 254–258 (1997). https://doi.org/10.1111/j.1540-8167.1997.tb00788.x
